# GLOM–TGITO: multiscale protein modelling and multiview landscape-guided sequence discovery

**DOI:** 10.64898/2026.10.22.760886

**Authors:** Yuchen Xiong, Yosef Kristian Tirta, Shiyu Wang, Zhenquan Hu, Wenjing Ma, Shuguang Yuan

**Affiliations:** Xiamen University Malaysia, Sepang, Selangor 43900, Malaysia; AlphaMol Science Ltd., Shenzhen 518055, China; School of Artificial Intelligence, The Chinese University of Hong Kong, Shenzhen, Shenzhen 518172, China; AlphaMol Science Ltd., Allschwil CH-4123, Switzerland; College of Life Sciences, Zhejiang University, Hangzhou 310027, China

**Author notes:** Correspondence: Wenjing Ma; Shuguang Yuan.

**Keywords:** protein design, fluorescent proteins, protein language models, fitness landscapes, offline model-based optimization, inference-time optimization

## Abstract

**Motivation:** Offline protein design must seek high-activity variants while retaining support for their predicted performance. The challenge is to explore multiple promising regions without confining the search to average-performing sequences. We test whether multiscale representations and local evidence can guide this search.

**Results:** GLOM–TGITO links a global–local fluorescence predictor to multiview landscape-guided optimization. Across 31,434 measured ppluGFP2 sequences, GLOM achieved random/depth-holdout RMSEs of 0.170/0.247 versus 0.247/0.324 for the strongest SaProt-650M controls. Averaged across three training seeds, 96.3%/97.3% of prediction-ranked Top 100 test variants were measured bright (log_10_ fluorescence ≥4.2). Search from 900 peak regions yielded 2,594 distinct candidate sequences. The selected Top 100 showed 2.9–8.2% lower mean representation deviation than four comparator pipelines; 38 versus 11–16 met joint novelty–reliability filters. Deviation identified measured test errors >0.1 (AUROC 0.773). The first molecular-dynamics run of the selected Top 30 identified smaller surface hydrophobic patches in 20 candidates relative to their starts, with representative variants showing more persistent chromophore contacts or narrower torsional fluctuations.

**Availability and implementation:** Code, model weights and reproducibility files are available at https://github.com/shycakj/GLOM-TGITO-multiscale-protein-modelling-and-multiview-landscape-guided-sequence-discovery.

**Supplementary information:** Supplementary methods, figures, tables and reproducibility records are provided.

## 1. Introduction

Protein engineering often starts with a collection of measured variants and a much larger set of sequences that have never been tested. The practical question is which variants to make next. For fluorescent proteins, one goal is to obtain brighter reporters from a limited set of measurements. Fluorescence depends on chromophore formation and its environment within the folded scaffold, so substitutions can affect brightness through both local changes and interactions with the rest of the protein (Sarkisyan et al., 2016; Gonzalez Somermeyer et al., 2022). Measuring every combination is impractical. Computational design must therefore learn from the available assay and use that knowledge to identify promising new variants.

Predictive models provide a way to learn these sequence– fluorescence relationships. Assay-trained models learn the effects of substitutions, while protein language models supply residue representations learned from large sequence collections (Alley et al., 2019; Rives et al., 2021; Hsu et al., 2022). Structure-aware models such as SaProt add information about local structural environments (Su et al., 2024), and pretrained representations can support protein engineering with relatively few measurements (Biswas et al., 2021). Applying these features to a particular mutational library still requires learning how its substitutions affect the measured phenotype. For GFP, the dependence of fluorescence on both local and wider context motivates a model that combines explicit mutation information with whole-protein, regional and key-site representations.

Once trained, such a predictor can be used to search for improved sequences. Model-guided evolution and adaptive sequence search use predicted activity to propose new variants (Wu et al., 2019; Sinai et al., 2020). We formulate sequence design as offline black-box optimization: the search for high-fluorescence sequences uses a fixed dataset of measured sequence–fluorescence pairs, without access to new evaluations of the experimental objective. A common difficulty in such offline black-box optimization is that the model may overestimate the activity of sequences with few similar training examples, leading the search towards candidates with high predicted scores but poor actual performance (Brookes et al., 2019; Trabucco et al., 2021). Accuracy on ordinary test variants does not establish reliability for inputs selected to maximize the predictor (Fannjiang et al., 2022; Greenman et al., 2025). The challenge is to seek high fluorescence while retaining support for the prediction, moving beyond average-performing sequences and exploring multiple promising regions.

The measured variants can guide this search as well as train the predictor. In a learned representation, they define a neighbourhood around a proposed sequence. Their measured fluorescence indicates whether that neighbourhood contains bright proteins, while their corresponding predictions show where the model tends to overestimate brightness. These two sources of information can distinguish a promising region with experimental support from one that appears favourable mainly to the predictor. The neighbourhood also depends on which representation is used: whole-protein, regional and key-site features group variants differently. Retaining these intermediate features therefore provides several views of the same measured landscape, from which the search can draw guidance as a sequence changes.

We implement this approach in GLOM–TGITO, a unified framework comprising a Global–Local Organization Model (GLOM) and Terrain-Gated Inference-Time Optimization (TGITO). GLOM learns fluorescence from SaProt-aligned residue features and explicit mutation embeddings, combining a global branch with regional and key-site branches linked by reciprocal cross-attention. Its intermediate representations supply views in which TGITO builds paired measured and predicted landscapes. TGITO starts from multiple supported peak regions and optimizes amino-acid probabilities through the frozen predictor. Predicted brightness and terrain guidance jointly drive the search; a dynamic gate updates the contributing views using local experimental support and overprediction risk. Final discrete sequences are ranked by a composite confidence score and selected under brightness and diversity constraints.

We evaluate this integrated framework on the measured ppluGFP2 landscape. Prediction benchmarks test how well GLOM learns fluorescence, and comparisons between complete discovery pipelines assess the candidates produced by search and selection. To evaluate support for novel sequences, we validate representation deviation against measured prediction errors and examine it alongside predicted brightness, terrain risk and sequence novelty after candidate batches are fixed. GLOM reduces prediction error relative to the matched baselines, while the complete pipeline selects candidates with lower mean representation deviation. Biophysical analysis of the selected Top 30 connects the designed sequences to changes in surface exposure and chromophore dynamics. These complementary effects suggest routes to fluorescent reporters through surface engineering and control of the chromophore environment.

## 2. Materials and methods

### 2.1. Design task and fluorescence data

Given measured sequences {(*x*_*i*_, *y*_*i*_)}, we seek unmeasured, high-fluorescence variants across multiple promising regions, with local experimental evidence supporting their predictions. The target *y* is log_10_ fluorescence. Search operates offline: the training measurements and fitted model remain fixed while the input sequence changes. The full task includes initialization, optimization and selection of the final batch.

The ppluGFP2 dataset contains 31,480 records and 31,434 unique 221-residue sequences after exact deduplication (Gonzalez Somermeyer et al., 2022). Residues are indexed without the initiating methionine. The sequence-hash random split contains 22,003 training, 3,144 validation, 3,143 calibration and 3,144 test variants. The mutation-depth protocol trains, validates and calibrates on 13,723, 1,715 and 1,716 sequences with at most two substitutions; its 14,280 test variants have 3–13 substitutions. Exact duplicates do not cross partitions. The random split is not homology-disjoint, and historical label exposure in feature-panel and policy selection is documented in Supplementary Sections S4 and S9.

We use fluorescence thresholds of 4.2 and 4.35 for the screening analyses. These cutoffs define bright variants on this assay scale. Landscape colours show the empirical percentile of displayed fluorescence among the 22,003 training labels. Blue changes to red at the training median (4.1303), so red denotes the upper half of this training distribution. Elevation is linear in log_10_ fluorescence.

### 2.2 Framework overview

GLOM–TGITO shares a representation between prediction and search (Fig. 1). GLOM learns whole-sequence, regional and key-site summaries. Eleven intermediate feature views and one fixed sequence projection organize twelve memories containing the same measured training variants. TGITO first finds supported peak regions in these views, then optimizes amino-acid probabilities using the predictor and locally interpolated experimental evidence. The gate selects six contributing views for each state. Final hard sequences are ranked by the native confidence score and filtered for brightness and diversity. Reliability analyses are applied after these candidate identities and rankings have been fixed.

**Figure 1.**
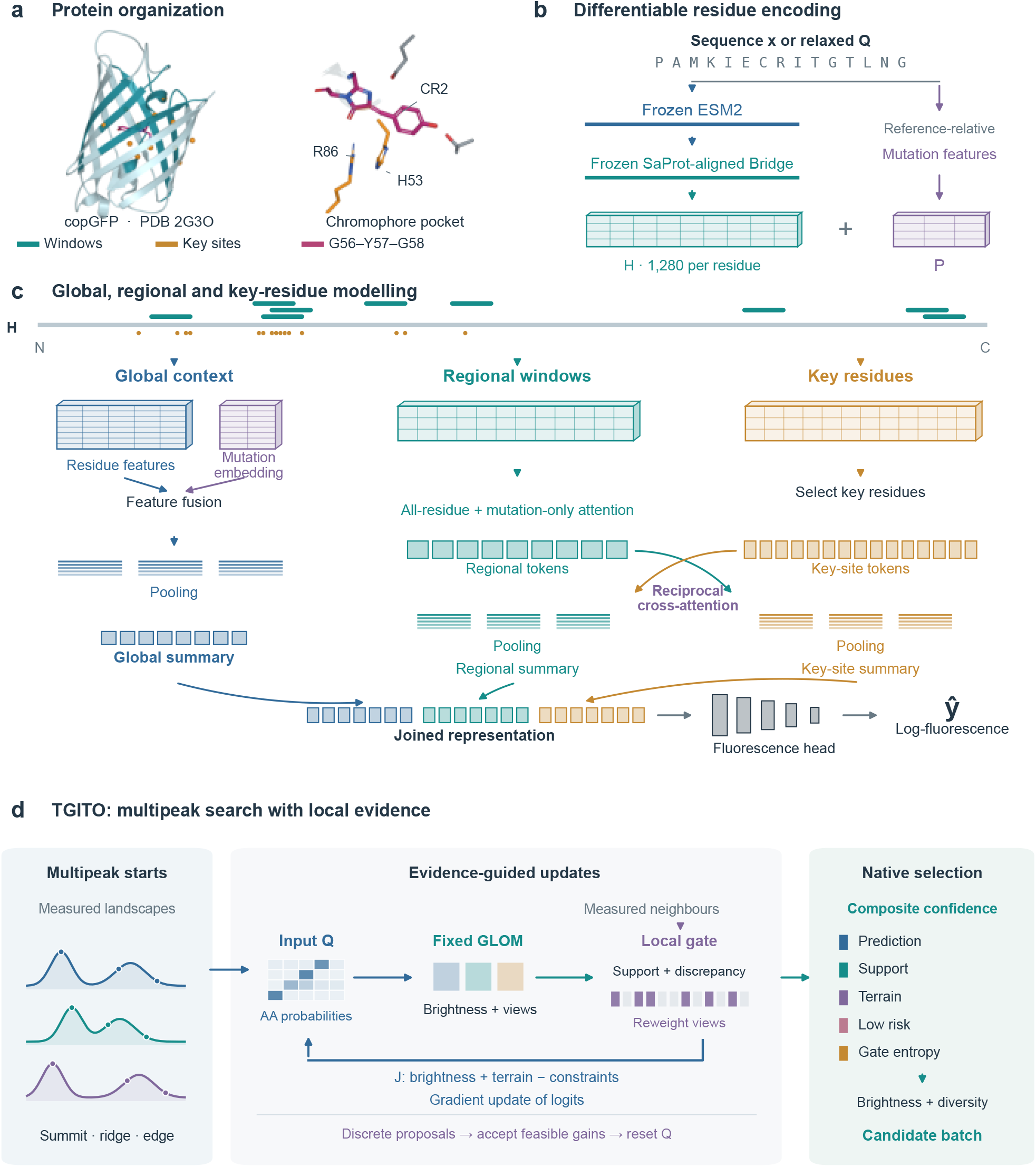
One representation supports fluorescence prediction and sequence discovery. **A**. Experimental reference structure (2G3O, chain C); windows, teal; key sites, amber; mature chromophore, magenta. Coordinates supply biological context. **B**. Frozen ESM2 and a frozen, SaProt-aligned Bridge map hard or relaxed sequences to 1,280-dimensional residue features *H*. Reference-relative mutation features *P* accompany *H* into GLOM. **C**. Explicit mutation features have a learned embedding in the global branch and define the regional mutation mask. Regional and key-site tokens exchange information through reciprocal cross-attention. Mean, maximum and attention pooling summarize each branch. Branch intermediates supply terrain views, while the joined representation feeds the fluorescence head. **D**. Multipeak starts enter a repeated input-update loop. The fixed GLOM queries measured neighbours; local support and prediction–measurement discrepancy determine the dynamic view gate. Gradients of the joint objective update amino-acid logits. Accepted feasible discrete proposals reset the relaxed input. Native confidence and diversity criteria select the final batch. Neighbourhood retrieval and gradients use the original feature spaces; UMAP displays their geometry. Feature and landscape glyphs are schematic; numerical settings are in Supplementary Methods S1–S4 and Table S1. **Alt text:** GFP structural regions and key sites guide a multiscale predictor. Frozen ESM2 and a SaProt-aligned Bridge supply residue features alongside mutation features. Global, regional and key-site branches predict fluorescence and supply landscape views. TGITO updates sequences from multiple peaks using local measured evidence, then selects candidates by confidence, brightness and diversity.

**Figure 2.**
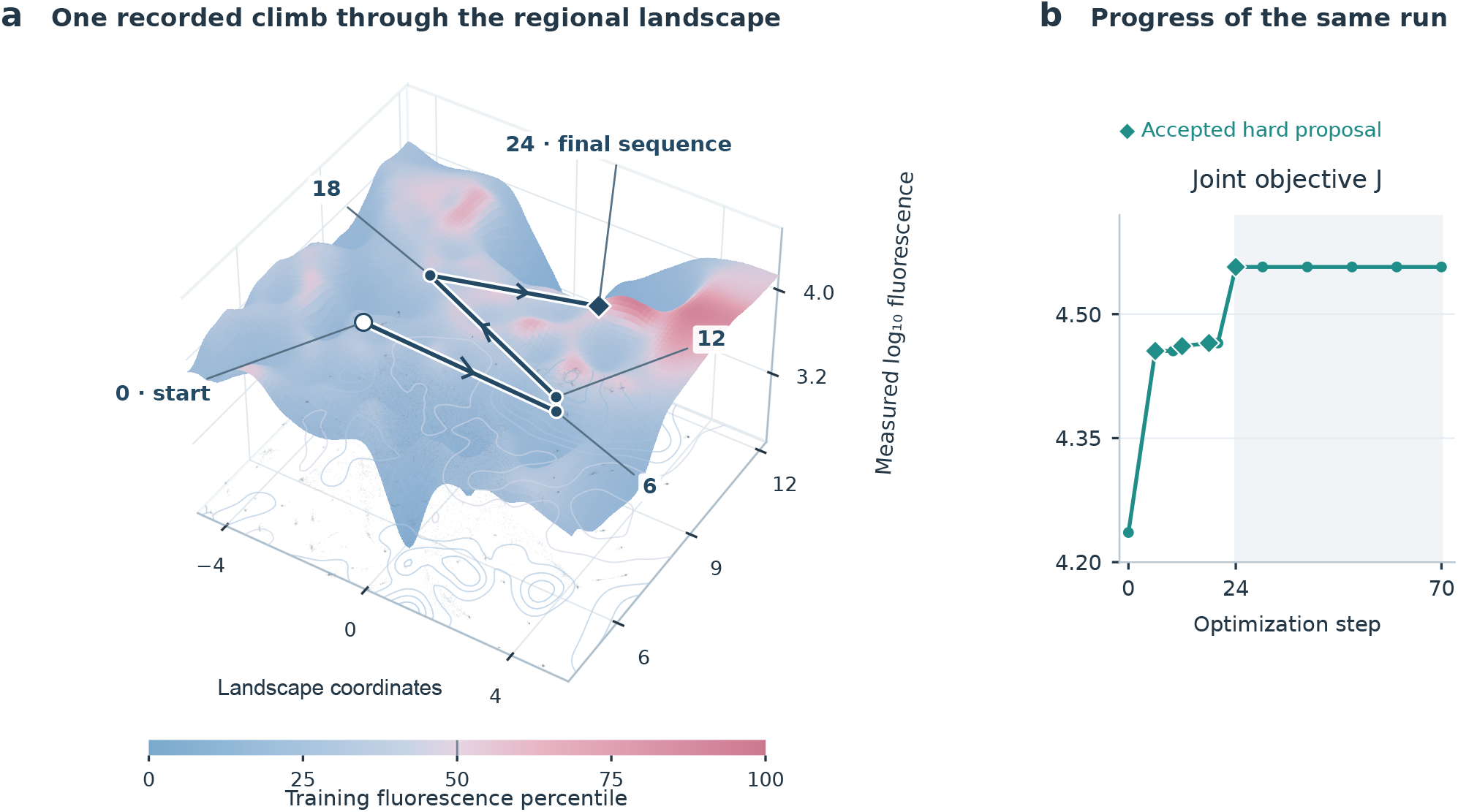
A recorded TGITO climb and its optimization progress. **A**. A regional landscape from the 22,003-sequence training memory, displayed over two UMAP coordinates. Colours encode training-label percentiles; red begins at the median (4.1303). The path connects initialization and four accepted-proposal checkpoints (steps 0, 6, 12, 18 and 24) of context 0054, selected by native Top 6 rank 1. Markers retain the logged neighbour-weighted coordinates of the relaxed queries; labels give optimization steps. Elevation follows the measured display surface, not candidate fluorescence or the joint objective. **B**. Joint objective *J* evaluated on the hard sequence at all twelve saved checkpoints. Diamonds mark accepted proposals; the final hard sequence persists from step 24 through 70. The objective includes terrain and seed-departure terms, so its changes need not track brightness alone. Other views, intermediate states and gate weights appear in Supplementary Figs S3 and S10. **Alt text:** Landscape height shows measured training fluorescence; blue-to-red colour shows its percentile. A path joins initialization and accepted checkpoints at steps 6, 12, 18 and 24. The adjacent joint-objective trace rises overall and plateaus from step 24 through 70.

### 2.3. Residue representation and mutation inputs

A sequence is represented by *Q* ∈ [0, 1]^221*×*20^, with one-hot rows for discrete evaluation or softmax probabilities during optimization. Frozen ESM2-150M supplies 640-dimensional residue states (Lin et al., 2023). A three-layer Bridge predicts probabilities over twenty Foldseek 3Di symbols plus a padding token and forms a 128-dimensional expected structural-token embedding (van Kempen et al., 2024). Concatenating the ESM2 state, expected structural-token embedding and 512-dimensional Bridge hidden state, followed by projection, gives a 1,280-dimensional residue representation *H*.

Bridge pretraining combines 3Di cross-entropy with cosine alignment to SaProt residue embeddings (Su et al., 2024). SaProt thus supplies representation-level supervision; inference and input optimization use the frozen, differentiable ESM2– Bridge pathway. The 3Di alphabet encodes local structural environments. Four reference-relative channels additionally encode mutation probability and the probability of a hydrophobic, polar or charged replacement. Their learned 4 →24 →24 projection gives the mutation embedding; the unchanged reference residue contributes zero to these channels (Supplementary Section S1).

### 2.4. Multiscale organization and supervised training

The global branch projects *H* to 256 dimensions, fuses the 24-dimensional mutation embedding and pools over the whole sequence. The regional branch represents nine ten-residue windows, combining attention pools over all residues and over mutation-masked positions in each window. The key-site branch retains fifteen individually selected residue tokens. These historically assay-informed panels are fixed for the present experiments; PDB 2G3O locates them relative to the chromophore (Supplementary Fig. S1).

Reciprocal four-head cross-attention lets regional tokens query the original key-site tokens and key-site tokens query the original regional tokens. Residual normalization and feed-forward updates precede pooling. The three 256-dimensional summaries form *z* = [*G*; *R*_*f*_; *K*_*f*_], a 768-dimensional representation passed to the fluorescence head. GLOM is trained using standardized Huber loss, AdamW and validation-Spearman checkpointing, with three training seeds. Layer dimensions, losses and optimizer settings are specified in Supplementary Sections S1 and S6.

### 2.5. Measured memories and dynamic terrain guidance

Twelve views comprise three global pools, two regional pools, five window descriptors, flattened key-site states and a fixed projected-sequence view. Each memory contains the same 22,003 random-split training observations. UMAP layouts define candidate peak regions and display the landscapes; online optimization instead uses cosine neighbourhoods in the original feature spaces (McInnes et al., 2018). Calibration variants assess source quality without entering the measured memory.

Local measured and predicted brightness distributions define relative brightness, bright-neighbour support and prediction– measurement disagreement. A positive local prediction coupled to weaker experimental evidence raises terrain risk. For each query and view, the 32 nearest memory rows are interpolated to form the local potential, support, risk and live overprediction gap. The potential weights measured and predicted relative brightness by 0.7 and 0.3. Calibration quality, support, risk and gap determine the dynamic gate over the six highest-scoring sources. Source identities and weights are recomputed at every relaxed or discrete evaluation (Supplementary Section S2).

Gradients flow through the original feature similarities and interpolated evidence within the selected neighbourhoods. Gate authority is detached, while source membership and regional mutation masks are hard selections. The displayed terrains provide a readable view of the evidence encountered by this computation.

### 2.6. Multipeak optimization and native batch selection

Source-balanced selection with uniform source priors retains 900 consolidated peak regions. Summit, ridge and edge roles define 2,700 search tasks using 1,308 distinct starting sequences; a sequence can serve more than one role or region. Each task has its own random seed. All positions are mutable, with a final budget of two substitutions relative to that task’s starting sequence.

TGITO optimizes amino-acid logits for 70 steps at fixed softmax temperature 0.8. Its objective combines predicted brightness and gated terrain potential, penalizes excessive substitutions and brightness loss, encourages departure from the start and includes a small entropy term. Every six steps, gradient-informed single- and paired-site substitutions generate up to 64 hard proposals. An accepted proposal resets the relaxed state around the new discrete sequence. The endpoint is the best eligible hard state, must differ from its start and must be absent from training, validation and calibration. The full objective and fixed hyperparameters are given in Supplementary Sections S3–S4.

The native confidence score combines normalized prediction, support, terrain potential, low risk and gate entropy with weights 0.35, 0.25, 0.20, 0.15 and 0.05. Confidence is computed before exact sequence deduplication. Greedy selection follows this ranking and requires predicted brightness ≥4.31 and pairwise Hamming distance ≥2. Top 6, Top 30 and Top 100 are nested prefixes under the same constraints. Hamming distance enforces sequence diversity.

### 2.7. Prediction and complete-discovery comparators

Supervised controls include ridge, Hamming kNN, frozen ESM encoders, ESM2 LoRA, a matched Transformer and four SaProt-650M configurations (Supplementary Sections S6 and S13). The SaProt controls use frozen mean-pooled embeddings with an MLP, mean-pool LoRA, and two CLS LoRA adaptations of the official fluorescence recipe (SaProtHub, 2026). The CLS inputs contain amino acids alone or predicted 3Di from the same frozen Bridge. All four SaProt controls start from the general pretrained encoder and fit new heads or adapters on our partitions; fluorescence-task pretrained weights are not used.

Complete discovery comparators are GLOM–AdaLead, GLOM–random, GLOM–ProtHMSO and ESM650–ProtHMSO (using an ESM2-650M predictor) (Sinai et al., 2020; Wang et al., 2026). They share 2,700 distinct bright training starts, a two-substitution radius, and the final brightness/diversity constraints. The baselines use predictor-based endpoint ranking. The comparison evaluates each complete procedure, including its own initialization and ranking. Counted fitness forwards are matched task by task at 1,476,063 per method. TGITO additionally uses 189,000 backward steps; total computation is not matched (Supplementary Section S5).

### 2.8. Accuracy, deviation and reliability evaluation

Prediction is assessed by RMSE, MAE, Pearson and Spearman correlations and *R*^2^. Measured brightness among prediction-ranked Top 6/30/100 test variants assesses screening yield. Neural results report means and sample SDs across three seeds; paired test-row bootstrap intervals quantify fixed-model error differences.

For a generated sequence, representation deviation measures how far its learned state lies from the observed training neighbourhood. We L2-normalize *z*, calculate Euclidean distance to its 32nd nearest training neighbour and divide by the high-prediction calibration Q95 distance, 0.5837557863 (Sun et al., 2022). Thus *D* = 1 marks the calibration Q95 boundary, with larger values indicating greater deviation from the training neighbourhood. This continuous descriptor is the primary candidate-support measure, accompanied by Q85–Q99 exceedances, brightness, terrain risk and sequence novelty (Fig. 4).

We first test whether deviation distinguishes measured prediction errors. Candidate rescoring and matched-risk analyses use the mean of two other GLOM fits, while deviation validation uses residuals from the search checkpoint. Combined risk is the gate-weighted terrain risk plus half the positive local gap between predicted and measured relative brightness (Supplementary Section S8). A secondary joint rule requires a calibrated lower-brightness reference ≥ 4.2, combined risk ≤0.15761, cross-seed SD ≤0.014576 and at least two substitutions from every visible variant. All thresholds and neighbour-rank sensitivity checks are reported in Supplementary Sections S7–S8.

### 2.9. Biophysical analysis of selected sequences

We analysed the native-ranked Top 30, their 25 distinct starting sequences and WT using mature-chromophore models based on PDB 2G3O chain C (Wilmann et al., 2006). All 56 systems underwent three independently initialized runs, each with the same preparation, 0.5-ns equilibration and 3-ns explicit-solvent production simulation at 300 K and 1 bar, using OpenMM, ff14SB and compatible anionic chromophore parameters (Eastman et al., 2024; Breyfogle et al., 2023). The first run supplies the primary comparisons; the other two assess repeatability. We compared each candidate with both WT and its actual start within each run. Readouts describe the largest non-interface hydrophobic surface patch, protein–phenolate hydrogen-bond occupancy and torsional fluctuations around the bonds linking the methine bridge to the imidazolinone (I) and phenolate (P) rings. Sequence rankings were fixed before these analyses; six examples were selected afterwards to illustrate distinct molecular changes. Surface-radius, terminal-segment, split-trajectory and additional-seed checks, together with full sequences and all candidate results, are given in Supplementary Sections S9 and S14 and Data 1, 3 and 4.

## 3. Results

### 3.1. GLOM reduces absolute error and enriches measured bright variants

GLOM achieved random-holdout RMSE 0.170 ±0.001 and MAE 0.083± 0.002 (n=3,144), and depth-holdout RMSE 0.247± 0.002 and MAE 0.131 ±0.002 (n=14,280). These were the lowest mean errors across the supervised configurations evaluated (Fig. 3; Supplementary Tables S2–S3). On the depth test, one-hot ridge had higher Spearman correlation (0.914 versus 0.870).

**Figure 3.**
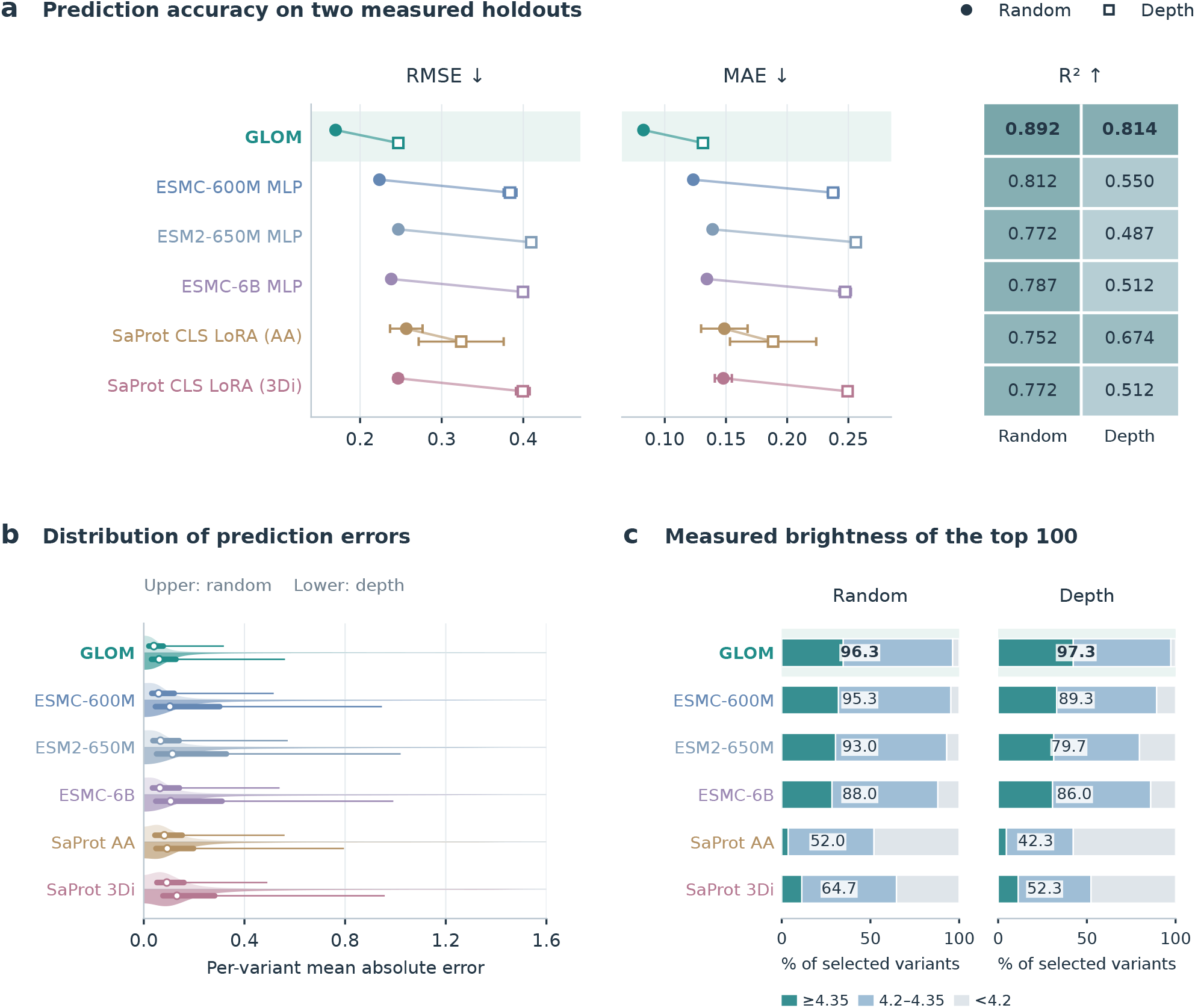
GLOM combines low prediction error with high measured screening yield. Six complete predictors use the same random/depth holdouts (n=3,144/14,280). **A**. RMSE and MAE: means and sample SDs over three training seeds; circles, random; squares, depth. Heatmap cells show mean *R*^2^. **B**. Per-variant absolute errors averaged across the three fits. Upper/lower densities show random/depth holdouts; circles, medians; thick lines, interquartile ranges; thin extensions, 95th percentiles. **C**. Measured brightness of each model’s prediction-ranked Top 100, averaged across three rankings. Bar segments show log_10_ fluorescence *≥* 4.35, [4.2, 4.35) and *<* 4.2; printed values are the percentage*≥* 4.2. These are measured test variants. The SaProt CLS controls use amino-acid-only or predicted-3Di input. All controls and batch sizes are in Supplementary Tables S2–S3 and S20–S22; architecture ablations are shown in Fig. S12. **Alt text:** Six predictors are compared on random and mutation-depth holdouts. GLOM has the lowest RMSE and MAE and the highest R-squared. Error distributions and screening bars complement these summaries. Its prediction-ranked Top 100 includes 96.3% and 97.3% measured bright variants, respectively, averaged across three training seeds.

**Figure 4.**
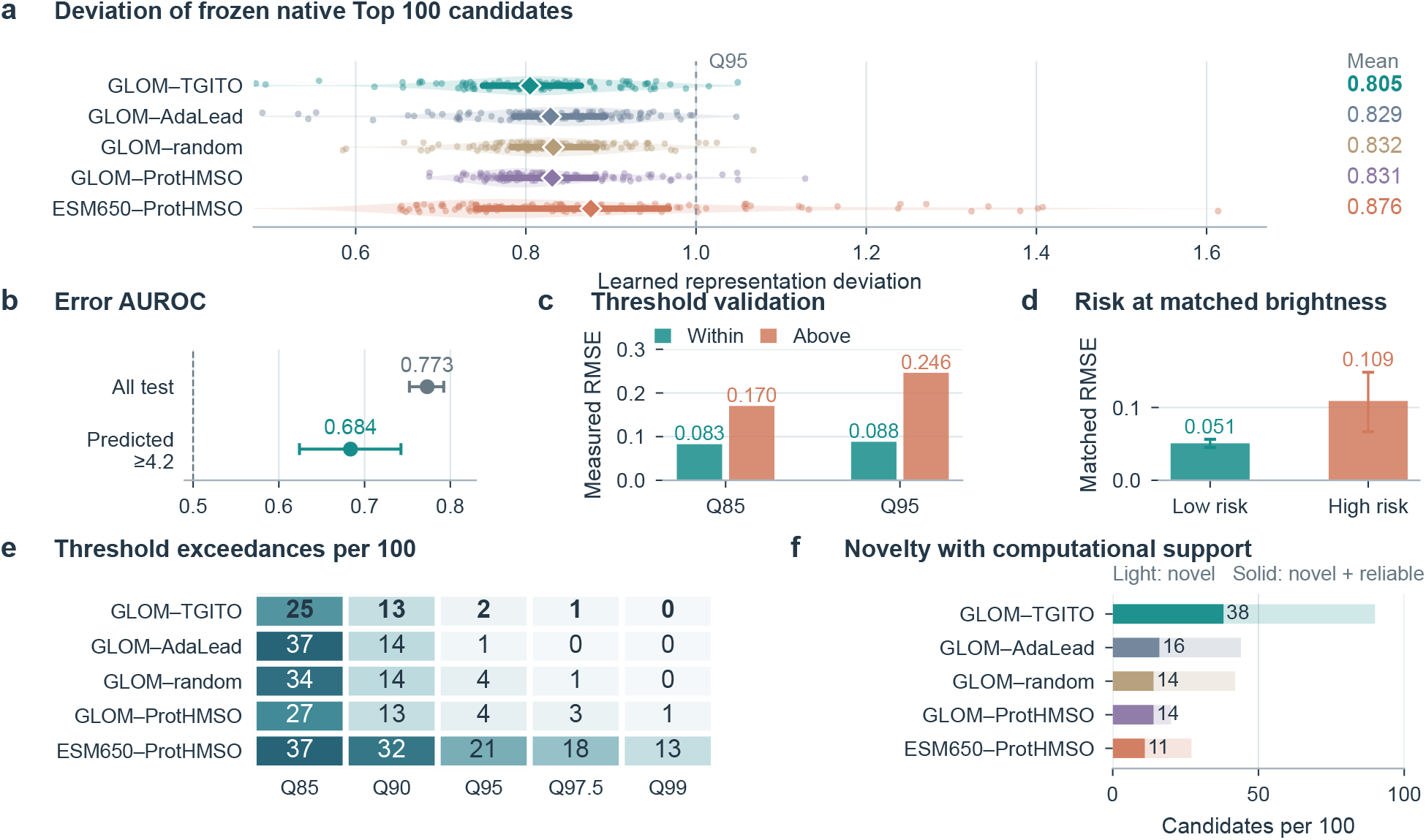
How unfamiliar are the optimized sequences to the predictor? Deviation measures distance from a candidate’s learned GLOM representation to the measured training neighbourhood, quantifying local model support; lower deviation indicates closer support. **A**. Frozen native Top 100 distributions: diamonds, means; thick lines, interquartile ranges; dots, sequences; dashed line, calibration Q95. **B–D**. Measured-label checks establish relevance to error: absolute-error AUROC (n=3,144/973 for all/high-prediction tests), RMSE within/above Q85 and Q95, and brightness- and distance-matched low/high-combined-risk RMSE (n=210/295). B,D show 95% bootstrap intervals. **E**. Exceedances per 100 at Q85–Q99. **F**. Candidates at least two substitutions from every visible sequence (light), also meeting the joint reliability rule (solid). Each selection comes from one campaign. Complete validations are in Supplementary Tables S6 and S15–S18. **Alt text:** GLOM–TGITO selects the batch with the lowest mean representation deviation. Measured-test analyses associate greater deviation and combined risk with larger errors. A grid compares Q85 to Q99 exceedances. Thirty-eight candidates meet the joint novelty-reliability rule, compared with 11 to 16 for four comparator pipelines.

Random- and depth-holdout *R*^2^ were 0.892 and 0.814. On the depth split, MAE was 44.8–48.8% lower than the frozen-encoder MLPs based on ESMC-600M, ESM2-650M and ESMC-6B. These absolute-error comparisons complement rank correlation (Fig. 3a). The per-variant error distributions show the same pattern: GLOM had median errors of 0.040 and 0.061 on the two holdouts (Fig. 3b).

The best SaProt-650M mean RMSEs were 0.247 ±0.005 (random, CLS with predicted 3Di) and 0.324 ±0.052 (depth, CLS with amino-acid input). GLOM reduced these RMSEs by 31.0% and 23.8%. Paired test-row bootstrap intervals for GLOM-minus-SaProt RMSE and MAE were below zero across four controls and both splits (Supplementary Table S21).

Averaged across three training seeds, the measured bright fractions (≥4.2) among GLOM-ranked Top 100 test variants were 96.3% and 97.3% on the random and depth splits, respectively, versus 52.0–64.7% and 42.3–61.7% for the SaProt controls. GLOM therefore enriched measured bright variants in both test splits (Fig. 3c; Supplementary Tables S14 and S22).

### 3.2. Complementary views support dynamic multiterrain climbs

The global, regional and key-site branches provide different descriptions of the same sequence. Global–regional and global– key CKA were 0.0023 and 0.0084, while regional–key CKA was 0.487. Matched ridge probes achieved RMSE 0.429, 0.305 and 0.293 on the individual branches and 0.192 on their concatenation. Thus the branches differ in geometry and jointly retain more linearly accessible fluorescence information (Supplementary Fig. S12c,d and Table S5). Architecture ablations are reported in Supplementary Section S6. TGITO uses these intermediate features and a fixed sequence projection to query measured neighbours in twelve views throughout optimization.

In the illustrated trajectory, six of twelve views contribute at each saved state and the active set changes four times as the query moves across the regional landscape (Fig. 2; Supplementary Fig. S10). Four discrete proposals are accepted over 70 steps. The final batch then combines predicted brightness with terrain support, potential, low risk and diversity.

### 3.3. Deviation and terrain risk distinguish measured prediction errors

On 3,144 measured test variants, deviation identified absolute prediction errors >0.1 (AUROC 0.773, 95% bootstrap interval 0.752–0.792). Among 973 variants predicted ≥4.2, AUROC was 0.684 for absolute error and 0.709 for overprediction. RMSE rose from 0.083 inside calibration Q85 to 0.170 outside, and from 0.088 inside Q95 to 0.246 outside (Fig. 4b,c; Supplementary Table S6).

Combined terrain risk also separated prediction errors when matched for predicted brightness and nearest-training Hamming distance: stratum-standardized RMSE was 0.0508 versus 0.1090 in low-versus high-risk groups (difference −0.0582; 95% conditional interval −0.0982 to −0.0166). The agreement between these two descriptors and measured residuals supports their use in candidate prioritization.

The calibration-based reliability rule accepted 80 measured test variants, of which 78 were bright (97.5%; Wilson interval 91.3–99.3%). Requiring at least two substitutions from every training sequence retained fourteen variants, all bright (78.5– 100% interval; Supplementary Table S16).

### 3.4. Integrated selection lowers deviation while retaining sequence novelty

Of 2,700 search tasks, 2,595 returned valid endpoints and yielded 2,594 distinct sequences; 2,593 were absent from the measured catalogue. We then compared how strongly each selected batch departed from the measured training neighbourhood in the common GLOM representation (Fig. 4a).

All five native Top 100 batches were absent from the measured catalogue. GLOM–TGITO’s mean deviation was 0.8048, compared with 0.8290, 0.8323, 0.8310 and 0.8763 for GLOM–AdaLead, GLOM–random, GLOM–ProtHMSO and ESM650–ProtHMSO. Its mean was 2.9–8.2% lower, and its within-batch 95th percentile was 0.952, compared with 0.971– 1.325. The lowest mean persisted across neighbour ranks 8/16/32/64 and native Top 6/30/100 batches (Supplementary Tables S7 and S13).

Native selection reduced GLOM–TGITO mean deviation from 0.9034 in its full endpoint pool to 0.8048 (10.91%), versus 0.03–3.77% in comparators. The ordering changed for unselected pools and raw-encoder distances (Supplementary Fig. S4).

At Q85, exceedance counts were 25/37/34/27/37 in the order above; at Q95 they were 2/1/4/4/21 (Fig. 4e). Supplementary Tables S9 and S18 report the full threshold grid, and Section S7 reports calibration-resampling sensitivity.

GLOM–TGITO selected Top 100 variants with terrain risk 0.118, support 0.317 and mean pairwise diversity 7.620 substitutions. Ninety were ≥ 2 mutations from every visible sequence; 38 satisfied the joint novelty–reliability rule compared with 11–16 for comparators. Joint novelty–reliability counts are shown in Fig. 4f; threshold comparisons are provided in Fig. 4e and Supplementary Tables S15 and S18.

Common cross-seed predicted brightness remained high at 4.391, compared with 4.400/4.401/4.414/4.317. Relative to AdaLead and GLOM–ProtHMSO, terrain risk decreased by 12.8%/6.2%, support increased by 32.2%/16.8% and diversity increased by 14.5%/25.2%. Reliability-only counts were 42/46/45/52/42; GLOM–ProtHMSO led predicted brightness and cross-seed stability. The integrated procedure therefore favoured sequence-distinct candidates with learned support.

### 3.5. Selected sequences alter surface exposure and chromophore dynamics

The Top 30 retain the G56–Y57–G58 chromophore-forming sequence while changing its surrounding scaffold. To examine the biological consequences of these designs, we compared the first mature-chromophore simulation of each candidate with WT and its starting sequence. Twenty candidates had a smaller non-interface hydrophobic surface patch than their starts, and eighteen were below WT (Fig. 5a). Mean patch area fell from 187.8 to 178.1 Å^2^, a 5.2% reduction. The decrease remained 4.5% when the 25 distinct starts were weighted equally. Twenty-five candidates had narrower I-axis fluctuations than WT. Nine combined a smaller patch with higher phenolate hydrogen-bond occupancy; eleven combined a smaller patch with narrower I-axis fluctuations relative to their starts (Fig. 5b; Supplementary Table S23).

**Figure 5.**
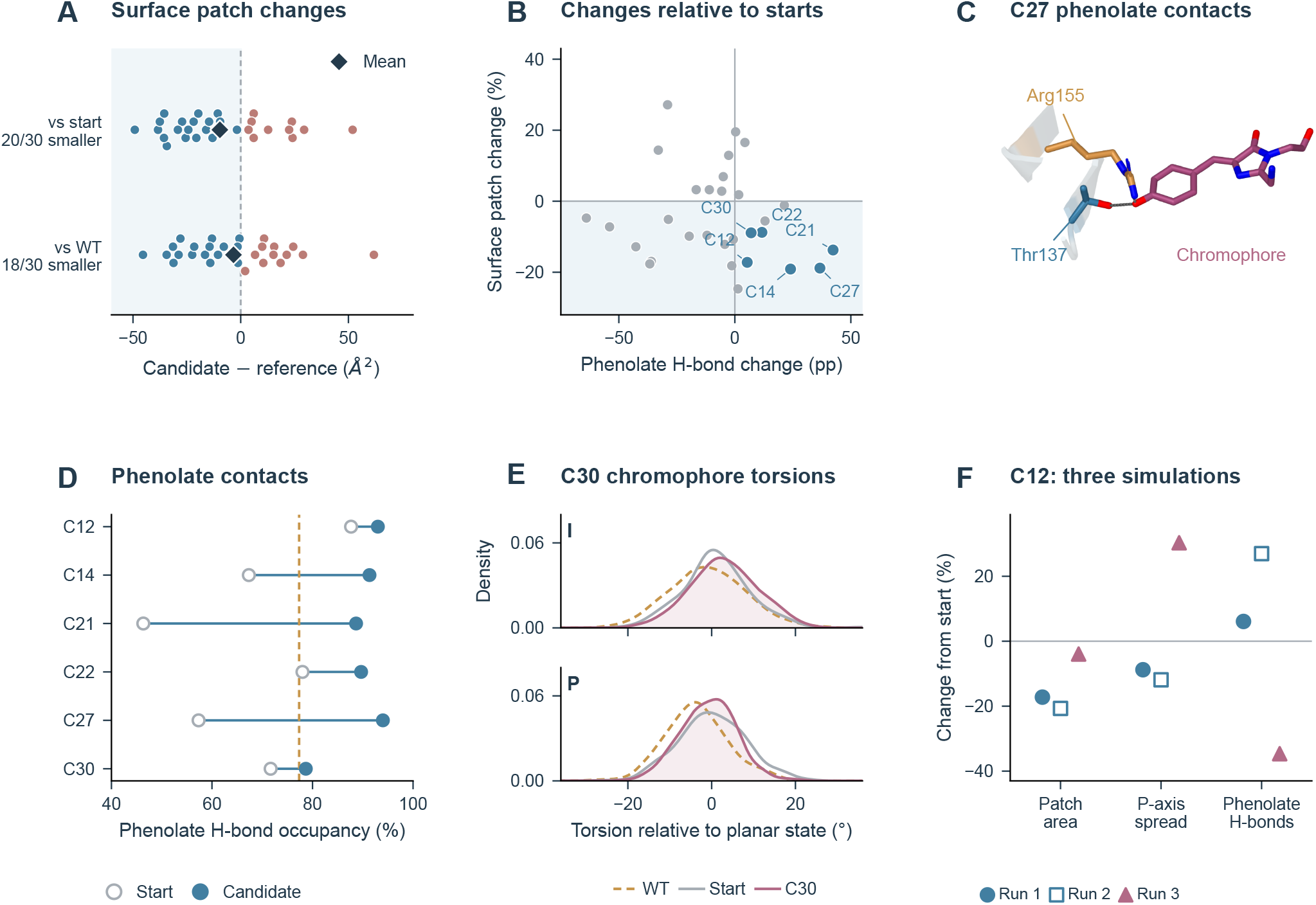
Biophysical changes in the selected Top 30 and representative sequences. **A**. Candidate-minus-reference differences in the largest non-interface hydrophobic surface patch, relative to the actual start or WT. Each dot is one candidate; vertical offsets separate overlapping points. Blue/red indicate decreases/increases; diamonds, means; dashed line, no change. The 30 candidate–start pairs contain 25 distinct starts. **B**. All candidates plotted by start-relative changes in patch area and phenolate hydrogen-bond occupancy; labelled points are the six examples. pp, percentage points. **C**. C27 conformation with simultaneous Thr137 and Arg155 contacts to the phenolate oxygen; dashed lines indicate the contacts. The conformation was selected from doubly contacted frames, while occupancies use all frames. **D**. Phenolate hydrogen-bond occupancy for the six examples; dashed line, WT. **E**. C30 chromophore torsion distributions alongside its start and WT. Angles are centred on the planar reference, 0° for I and 180° for P. **F**. C12 changes in three independently initialized simulations, expressed as percentages of the corresponding starting-sequence values. Panels A–E use the first 3-ns production trajectory per system; F shows all three runs. Surface patches use a 5 Å radius and include modelled residues 219–221 while excluding the reference dimer interface. Figures S13–S14, Tables S23–S25 and Supplementary Data 3–4 provide structures, alternative surface definitions and complete run-level comparisons. **Alt text:** Surface hydrophobic patches decrease in 20 of 30 candidates relative to their starts and in 18 relative to WT. Representative structures, contact occupancies and torsion distributions illustrate chromophore-environment changes. C27 shows Thr137 and Arg155 contacts. Three C12 simulations retain smaller patches but differ in hydrogen-bond and torsional changes.

C12, C14, C21, C22, C27 and C30 illustrate these different combinations, with their original Top 30 ranks retained (Supplementary Table S12 and Fig. S13). Surface changes were pronounced in C12, which adds E122K/A219D to its start, and C27, which adds F167V/V218E. Their patch areas fell by 17.2% and 18.8%, respectively. C14 and C27 also retained smaller patches when the modelled terminal residues were excluded. Across all three simulations, C12 and C14 retained smaller surface patches than their starts; C12’s phenolate-contact and P-axis responses varied between runs (Fig. 5f; Supplementary Table S25). Surface hydrophobicity is relevant to fluorescent-protein yield: reduced aggregation supported efficient maturation in Cycle3 GFP, and surface engineering improved reporter behaviour in the closely related TurboGFP (Fukuda et al., 2000; Evdokimov et al., 2006).

The chromophore environment changed through a different route. C21 and C22 increased phenolate hydrogen-bond occupancy from 46.3% to 88.7% and from 78.0% to 89.7%, respectively (Fig. 5d). In C27, simultaneous contacts with Thr137 and Arg155 occupied 50.0% of frames, compared with 9.3% in its start and 24.7% in WT (Fig. 5c). These residues contact the phenolate in copepod GFP structures (Bomati et al., 2014). A more persistent local hydrogen-bond network offers a plausible route to tuning the emissive chromophore environment, as shown in MD-guided engineering of the brighter YuzuFP variant (Ahmed et al., 2025).

C30 instead showed a clearer change in chromophore motion. Its I- and P-axis circular SDs were 7.90° and 6.53°, respectively: 11.5% and 11.4% below WT, and 4.2% and 17.9% below its start (Fig. 5e). Both decreases persisted in each trajectory half. Mean-square fluctuations of chromophore heavy atoms also fell by 29.6% relative to the start. Restriction of chromophore twisting is relevant because bond rotation provides a non-radiative decay route, whose energetics depend on the surrounding protein (Romei et al., 2020). Together with the surface effects, these contact and motion changes provide a molecular rationale for prioritizing distinct designs for reporter development.

## 4. Discussion

GLOM–TGITO seeks high-fluorescence candidates across multiple promising regions while using local evidence to assess their predictions. Its multiscale representation reduces fluorescence prediction error on both ppluGFP2 holdouts and supplies complementary views of the measured landscape. The complete discovery procedure then selects sequence-distinct candidates with lower mean learned-representation deviation than the four comparator pipelines. Measured-test analyses give this descriptor an empirical connection to prediction error.

The discovery comparison highlights the value of selecting sequence-distinct candidates with local model support. GLOM– TGITO retained more candidates meeting the joint novelty– reliability criterion while maintaining high predicted brightness. The Top 30 analysis adds a molecular view of the output. Surface-patch reductions recur across the selected sequences, while individual candidates differ in their effects on phenolate contacts and chromophore motion. These changes connect the search results to complementary aspects of fluorescent-reporter function.

This retrospective evaluation covers one fluorescent protein and one discovery campaign per method. Random-split variants can be closely related, and historical panel and policy selection included assay data overlapping the current tests. Forward-budget matching excludes TGITO’s additional backward computations. Repeated campaigns, nested panel and policy selection, and tests on additional proteins would extend this evaluation.

GLOM–TGITO combines high predicted activity, local experimental support and exploration across multiple regions. Its biophysical readouts further distinguish designs aimed at surface compatibility from those that alter the chromophore environment, linking sequence discovery to concrete molecular properties of fluorescent reporters.

## Author contributions

Yuchen Xiong: Conceptualization, Methodology, Software, Investigation, Formal analysis, Validation, Visualization, Writing—original draft. Yosef Kristian Tirta: Software, Investigation, Formal analysis, Writing— original draft. Shiyu Wang and Zhenquan Hu: Software. Wenjing Ma and Shuguang Yuan: Supervision, Resources, Writing— review & editing.

## Acknowledgements

The project received a Gold Award in the Protein Design Track of SynBio Challenges 2026. Research and computational resources were provided by Wenjing Ma, Shuguang Yuan and AlphaMol Science Ltd. OpenAI Codex assisted with language editing and figure preparation; further details are provided in Supplementary Section S15.

## Data and code availability

Code, model weights, fixed data partitions, training seeds and reproduction instructions are available in the GLOM–TGITO repository (release repro-v2026.10.28). The published ppluGFP2 assay is described in Gonzalez Somermeyer et al. (2022). Numerical comparison results, plotting inputs and protocol records accompany the Supplementary Information. Supplementary Data 1 provides the Top 30 sequences, their starting sequences and six representative designs; Data 2 documents historical sample overlap; Data 3 contains the biophysical protocol, first-run metrics, structural conformations and matched training-background contrasts. Data 4 provides all three simulation runs, with candidate-level changes relative to WT and actual starts.

