## Supplementary Information for "GLOM–TGITO: multiscale protein modelling and multiview landscape-guided sequence discovery"

**Abstract**

This supplement provides the complete GLOM representation and TGITO optimization specifications, measured-error validation, supervised and discovery comparators, and reproducibility details. Section S9 records the Top 30 sequence panel and the six illustrated representatives. Historical assay exposure during feature-panel selection and hyperparameter optimization is described in Sections S4 and S9. Section S6 and Fig. S12 contain the architecture controls, measured prioritization and branch diagnostics; Section S13 gives the four SaProt comparisons, and Section S14 reports the Top 30 biophysical comparisons against WT and the actual search starts. Discovery batches were fixed before the reliability analyses.

### S1. Reference, mutation features and GLOM architecture

Residue positions refer to the following 221-residue reference, without the initiating methionine; spaces are included only for readability:

```
PAMKIECRITGTLNGVEFELVGGGEGTPEQGRMTNKMKSTKGALTFSPYLLS
HVMGYGFYHFGTYPSGYENPFLHAINNGGYTNTRIEKYEDGGVLHVSFSYRYE
AGRVIGDFKVVGTGFPEDSVIFTDKIIRSNATVEHLHPMGDNVLVGSFARTFS
LRDGGYYSFVVDSHMHFKSAIHPSILQNGGPMFAFRRVEELHSNTELGIVEY
QHAFKTPIAFA
```

The amino-acid alphabet is ACDEFGHIKLMNPQRSTVWY. An input Q has dimensions 221×20 and contains either one-hot residues or relaxed probabilities.

At residue $i$, the mutation indicator is $1 - Q_i(r_i)$, where $r_i$ is the fixed reference residue. The other three channels sum $Q_i(a)$ over non-reference amino acids in the hydrophobic (A,V,I,L,M,F,W,P,G), polar (S,T,N,Q,C,Y) and charged (D,E,K,R,H) classes. All four channels are zero at an unchanged position in a hard sequence. They occupy columns 0, 7, 8 and 9 of the ten-channel input container. These reference-relative features describe the mutation itself; search constraints count substitutions relative to each starting sequence.

The fixed key positions are 24, 33, 35, 36, 52, 53, 55, 56, 57, 58, 59, 62, 84, 86, 100. The nine windows are 53–62, 55–64, 51–60, 207–216, 203–212, 77–86, 27–36, 165–174, 97–106. These historically consolidated panels were informed by mutation-indicator ridge analysis; their selection provenance and assay exposure are detailed in Section S9. Supplementary Fig. S1 maps them to the experimental reference structure. The three-dimensional coordinates provide structural context for the selected positions; model input features follow the sequence-derived path described below.

Frozen ESM2-150M supplies 640-dimensional residue states [1]. The Bridge projects them to 512 dimensions and applies three Transformer layers, each with eight heads, feed-forward width 2,048, pre-normalization and dropout 0.1. A 21-way head predicts probabilities over twenty Foldseek 3Di symbols and one padding token [2]. Their expectation under a learned 21×128 embedding table is concatenated with the original 640-dimensional ESM2 states and 512-dimensional Bridge states, then projected 1,280→1,024→1,280 to produce H. The output is a SaProt-aligned residue representation: Bridge pretraining combines masked 3Di cross-entropy with a 0.2-weighted masked cosine loss against SaProt residue embeddings [3]. SaProt targets were generated with westlake-repl/SaProt_650M_AF2; no smoothness regularization was used. The Bridge is frozen for the present GLOM fits and discovery campaign. Thus SaProt supplies representation-level supervision, while inference uses the continuous ESM2–Bridge pathway.

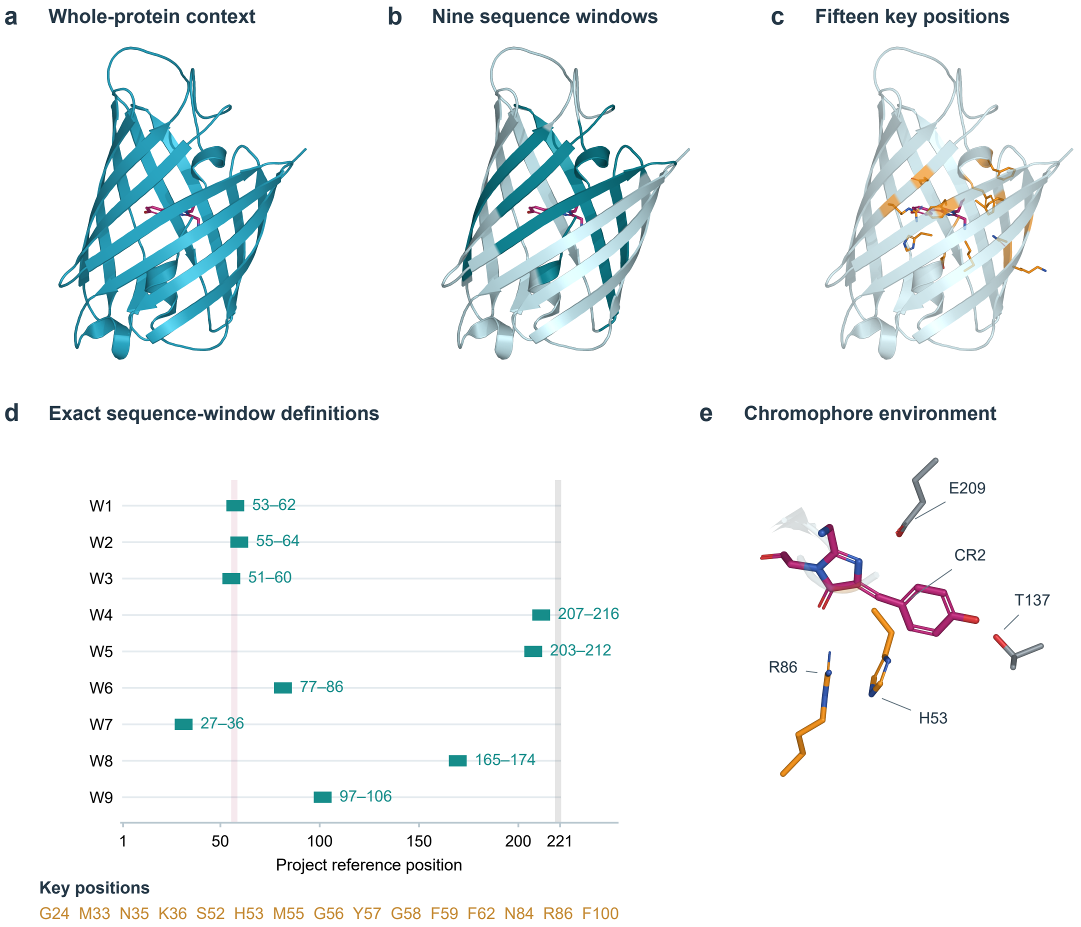


**Figure S1: Sequence windows and key positions in the reference structure**. a–c, Experimental chain C of PDB 2G3O (2.1 Å resolution) is rendered as a cartoon, with the mature CR2 chromophore in magenta. The observed chain is shown in a; the union of the nine sequence windows is highlighted in teal in b; the selected key positions are shown in amber in c. Overlapping windows remain distinct tokens in GLOM. d, Exact reference-coordinate ranges of the nine ten-residue windows. The magenta band marks project G56–Y57–G58; the grey band marks the unresolved terminal positions. e, The observed CR2 environment includes selected H53 and R86 sidechains in amber and neighbouring T137 and E209 in grey. Nitrogen and oxygen atoms use blue and red. Expanding the deposited CR2 precursor gives a sequence identical to an initiating methionine followed by the 221-residue project reference. Project G56, Y57 and G58 (precursor author G57, Y58 and G59) form one mature CR2 component at author residue 57. Chain C has coordinates for project positions 1–218; residues 219–221 are not modelled in this rendering. The panels display deposited geometry and the implemented selections, rather than additional structural inputs to the fluorescence predictor.

The global branch projects H through 512 and 256 dimensions. The four mutation channels have their own learned $4 \to 24 \to 24$ projection with GELU activations; concatenation with the residue features is followed by $280 \to 256$ fusion. Mean, maximum and attention pooling give 768 dimensions, compressed through 768 to the 256-dimensional global summary G. The regional branch has a separate H projection through 512 and 256 dimensions. For each window, it concatenates attention pools over all residues and over positions with mutation probability above 0.5; an empty mutation mask returns zero. The resulting 512-dimensional descriptor is compressed through 512 to 256 dimensions. A third H projection, $1280 \to 256 \to 256$, supplies fifteen key tokens. Thus the learned 24-dimensional mutation embedding enters the global branch directly, while the mutation indicator defines regional pooling; key tokens receive regional information through cross-attention.

Separate four-head attention modules let key tokens attend to regional tokens and regional tokens attend to key tokens. Both use the original token sets. Residual layer normalization, a shared 256→512→256 feed-forward network and a second normalization update each set. Mean, maximum and attention pools are compressed 768→768→256 into Rf and Kf. The concatenation z=[G;Rf;Kf] has width 768. The regression head uses widths 512,256,128,64,32,1, GELU activations and dropout 0.30,0.25,0.15 after its first three hidden activations. Global, regional and key projection dropout rates are 0.20,0.10,0.05; global compression, regional compression and the shared feed-forward network use 0.20,0.12,0.15. Attention pooling uses a learned scalar linear score and normalized masked softmax. The h32 feature is the linear output before its GELU activation. Main Fig. 1 summarizes the architecture; Supplementary Fig. S1 maps the selected positions and windows onto the reference structure.

#### S2. Multiview memory, support and dynamic gating

The twelve views comprise three global pools before compression (256 dimensions each), mean and maximum pools over the updated regional tokens (256 each), five window descriptors before compression (512 each), flattened key tokens before attention (3,840), and a fixed sequence projection (512). The selected window descriptors cover positions 165–174, 77–86, 97–106, 27–36 and 55–64. The sequence view projects 4,420 one-hot features through a Gaussian matrix with entry standard deviation $1/\sqrt{4420}$ and seed 13202; relaxed search applies the same matrix to flattened Q. This fixed projection is one search view. Familiarity evaluation instead uses learned z768, and neither z768 nor h32 is an active terrain source.

Each view contains the same 22,003 training observations. UMAP uses cosine distance, 30 neighbours, minimum distance 0.08 and deterministic source-specific seeds [4]. For each point, 48 layout neighbours excluding itself define measured and predicted brightness means and population standard deviations, with $10^{-6}$ added to each standard deviation. Scores $z_i^t$ and $z_i^p$ standardize the point's measured and predicted brightness against those neighbours. Fractions $h_{i,4.20}$ and $h_{i,4.35}$ count measured bright neighbours. The raw disagreement score is:

$$r_i = \max(z_i^p, 0)\max(-z_i^t, 0) + \max(z_i^p - z_i^t, 0) \tag{S1}$$

For each source, a standardized-input MLP classifier with hidden widths 48,24 predicts the training-memory label y≥4.35. Its thirteen inputs comprise two UMAP coordinates, the point prediction, predicted neighbour mean/standard deviation/z-score, measured neighbour mean/standard deviation/z-score, the two bright fractions, positive signed z-score gap and raw disagreement. It uses ReLU, regularization $10^{-4}$, initial learning rate $5\times10^{-4}$, at most 220 iterations and internal early stopping with a 15% validation fraction. Its probability p_i is a descriptor of an observed memory row, whose measured brightness is available; it is not applied to an unknown candidate's true brightness. Stored support and risk are:

$$S_i = \mathrm{clip}(0.65p_i + 0.25h_{i,4.35} + 0.10h_{i,4.20}, 0, 1.5) \tag{S2}$$

$$R_i = \mathrm{clip}\big(0.70r_i + 0.30(1 - p_i)\max(z_i^p, 0), 0, 4\big) \tag{S3}$$

Source quality is estimated separately on 3,143 calibration queries. Their support scores interpolate the 32 nearest normalized training features with weights proportional to exp(cosine similarity/0.15). Quality Q_s is clip(0.70×AUROC+0.30×precision@60,0.05,1.5), evaluated against calibration brightness ≥4.35. Calibration-query labels evaluate scores but do not enter their interpolated descriptors. AUPRC is recorded but does not contribute to quality.

Online inference retrieves 32 training neighbours by cosine similarity in each original feature space, using softmax temperature 0.15. Let $L_s^t$ and $L_s^p$ be the weighted positive parts of the stored measured and predicted z-scores. Let $B_s^t$ and $B_s^p$ be the corresponding weighted brightness values. The source potential is:

$$P_s = 0.7L_s^t + 0.3L_s^p + 0.35(0.7B_s^t + 0.3B_s^p) \tag{S4}$$

Interpolated support $S_s$ and risk $R_s$ determine which sources contribute. Authority $A_s$ is the detached value $0.7L_s^t{+}0.3L_s^p$, bounded below by $10^{-6}$. The quality exponent is 0.8665330602. Support, risk and gap coefficients are 0.5249775580, 0.7454479755 and 0.1654493334. The six highest-scoring sources are combined at gate temperature 0.2037978423:

$$g_s = A_s Q_s^{\gamma}(1 + \lambda_S S_s) - \lambda_R R_s - \lambda_D \max(L_s^p - L_s^t, 0) \tag{S5}$$

$$T(x) = \sum_{s\in\mathrm{Top6}(g)} \frac{\exp(g_s/\tau_g)}{\sum_{r\in\mathrm{Top6}(g)}\exp(g_r/\tau_g)} P_s(x) \tag{S6}$$

The live gap is the positive difference between the two interpolated positive-part summaries, not the interpolation of the stored signed gap. Source identities and weights are recomputed at every soft or hard forward evaluation. Top-k membership and regional mutation masks are hard selections; authority is detached. Within these selections, gradients flow through similarities, interpolation and the remaining gate terms. UMAP is not differentiated, and interpolated query coordinates serve diagnostics rather than a second online neighbour search.

#### S3. Peak initialization, constrained search and final selection

Each UMAP plane is divided into a 50×50 grid. A peak is the brightest point of an occupied cell and must exceed the maxima of its neighbouring occupied cells. Its region contains all observations in the surrounding 3×3 cells and must contain a measured sequence with brightness ≥4.2. At most 120 regions per source are retained. Preliminary scores weight source quality, fraction≥4.30, normalized maximum brightness, fraction≥4.35, inverse mean source risk, uniform source prior, global-top200 membership and normalized peak brightness by 0.22,0.18,0.16,0.12,0.12,0.10,0.06,0.04. Brightness normalization clips (y−4.2)/0.25 to [0,1]; inverse risk is 1/(1+R), with R averaged over bright training rows.

Union-find merges regions with row-set Jaccard overlap≥0.92 and differences in mean, 90th percentile and fraction≥4.35 no larger than 0.035,0.05,0.06. Group scores retain the first six terms, averaging member qualities, risks and priors. The last two terms become 0.06f(n)+0.04min(1,n/4), where n is the number of represented sources and f(1)=1, otherwise f(n)=min(0.6,0.1n). Thus the grouping score is not a monotone consensus reward. Source-balanced round-robin selection retains 900 of 941 consolidated groups from 943 retained peaks.

The summit is the brightest row with y≥4.2. The ridge maximizes normalized brightness plus 0.30d/(d90+$10^{-6}$), where d is distance to the summit and d90 its 90th percentile among bright rows; the summit is excluded when possible. The edge is farthest from the summit among rows meeting max(4.2,min(4.35,median bright-row brightness)). Distances use the highest-scoring member's source coordinates. The 900 regions each define three role-specific search tasks, giving 2,700 tasks from 1,308 distinct starting sequences. Reuse occurs both across regions and between roles within a region. Each task has a distinct random seed. Of the 900 regions, 620 have three distinct role sequences, 234 have two and 46 have one.

All positions are mutable. Starting-residue logits are 5, alternatives 0, with Gaussian noise of standard deviation 0.15. Q=softmax(logits/0.8) uses a fixed temperature. For the following count equation, p_i means seed-change probability 1−Q_i(s_i), b=19/[exp(5/0.8)+19], $\theta$=clip(b+0.08,0.05,0.75), and $\tau$=0.035. The correction subtracts the mutation probability introduced by softmax initialization:

$$c_i = \mathrm{clip}\left(\frac{f(p_i) - f(b)}{f(1) - f(b)}, 0, 1\right), \quad f(u) = \tau \log(1 + \exp((u - \theta)/\tau)) \tag{S7}$$

Let d_all sum every corrected site count, d8 sum the eight largest, $\bar{H}$ be mean categorical entropy over 221 positions and y0 the hard starting-sequence prediction. The relaxed objective is:

$$\begin{aligned} J = \widehat{y} + 0.0464303621T - 0.2224716472[1 - d_8]_+^2 \\ - 0.25[d_{\mathrm{all}} - 2]_+^2 + 0.001\bar{H} - 12[y_0 - 0.05 - \widehat{y}]_+^2 \end{aligned} \tag{S8}$$

Seventy ordinary gradient steps on amino-acid logits use learning rate 0.0388846264. A proposed continuous step is retained if relaxed brightness is at least y0−0.05 and the relaxed objective drops by no more than 0.01. Otherwise the previous state is restored. Every six steps, logits minus the learning-rate-scaled latest loss gradient rank alternatives. Up to four alternatives per position define valid singles; pairs combine different positions among the highest-ranked 64 singles. Up to 64 combined proposals are scored as hard sequences. Hard scores use integer mutation counts and zero entropy. The best proposal must respect the two-substitution seed-relative budget, brightness floor and current hard objective. Acceptance resets relaxed logits around that proposal, while the original seed remains the budget reference.

The endpoint maximizes hard objective among eligible recorded states, sampled every ten steps, after accepted proposals and at termination. It must differ from the start, satisfy the hard budget and be absent from visible training, validation and calibration sequences. Runs without an eligible endpoint are excluded from candidate ranking. Valid endpoints are assigned the native composite:

$$C = 0.35c_{\text{pred}} + 0.25c_{\text{support}} + 0.20c_{\text{terrain}} + 0.15c_{\text{lowrisk}} + 0.05c_{\text{entropy}} \tag{S9}$$

Here c_pred=clip((prediction−4.2)/0.25,0,1). Let N denote min–max normalization across all valid endpoint rows. Support, potential and gate entropy are normalized separately; c_lowrisk=1−N($\bar{R}$+0.5$\bar{\Delta}$), where bars denote gate-weighted averages and $\Delta_s = \max(L_s^p - L_s^t, 0)$ is the positive live gap. Constant inputs normalize to zero. Confidence is computed before exact sequence deduplication. Descending confidence, with a lexical sequence tie-break, defines greedy Top6/30/100 selection requiring prediction≥4.31 and pairwise Hamming distance≥2. Each batch uses the same ranked candidate pool and constraints, differing only in the stopping count: Top6 is the first six entries of Top30, which is the first thirty entries of Top100. Confidence serves as a heuristic ranking score for this diversity-constrained selection.

### S4. Historical tuning and runtime verification

Historical multivariate TPE with successive halving tuned ten parameters: terrain and exploration weights, learning rate, source count, gate temperature, support/risk/gap coefficients, quality exponent and proposal interval. Fidelities used four starts for 25 steps, eight for 45 and sixteen for 70. The archived study contains 39 records: three failed, 31 pruned and five complete. Four configurations were confirmed in three stochastic batches of 24 starts at 70 steps. Its objective was:

$$J_{\text{HPO}} = 0.50T_{\text{treasure}} + 0.20C_{\text{coverage}} + 0.20G_{\text{predicted}} + 0.10D_{\text{unique}} \tag{S10}$$

Treasure rewards proximity within two substitutions to measured development targets, weighting distances 0/1/2 by 1/0.7/0.4 and normalized target brightness by power 1.5; coverage counts target neighbourhoods reached. Predicted gain is clipped to a 0.25 brightness scale, and diversity is the unique-sequence fraction. These historical Hamming terms are not the present learned-representation familiarity endpoint. Historical targets and their relation to the current partitions are documented in Section S9 and Supplementary Data 2.

The parameters in Table S1 were fixed for the discovery campaign. ESM unmasked-input scaling was 0.88, with mutation channels as defined in Section S1. Relaxed mutation counts were used for gradient updates and discrete objectives for hard proposals. Search used float32, whereas historical tuning used bfloat16. Hard- and soft-input encoders agreed on five feature-consistency probes. Reconstructed predictions differed from cached predictions by RMSE 0.001003 and maximum absolute error 0.008092; the cache used float16 H features and bfloat16 GLOM operations.

**Table S1:** Historical trial-12 parameters and effective runtime values

| Parameter | Historical | Runtime |
|---|---|---|
| terrain_weight | 0.0464304 | 0.0464304 |
| seed_explore_weight | 0.222472 | 0.222472 |
| lr | 0.0388846 | 0.0388846 |
| terrain_topk | 6 | 6 |
| terrain_gate_temp | 0.203798 | 0.203798 |
| terrain_support_weight | 0.524978 | 0.524978 |
| terrain_risk_weight | 0.745448 | 0.745448 |
| terrain_pred_gap_weight | 0.165449 | 0.165449 |
| terrain_source_quality_power | 0.866533 | 0.866533 |
| proposal_every | 6 | 6 |

All ten parameters use the confirmed historical values. Historical policy selection used labels overlapping the current test catalogue (Section S9). The accompanying parameter CSV gives full numerical precision and configuration fields. Search uses float32 (amp_ dtype=off), whereas historical tuning used bfloat16; arithmetic precision was not an HPO dimension.

A preliminary implementation check comprised 36 contexts spanning twelve sources and three roles, two stochastic repeats and three configurations: 216 trajectories, plus 24 baseline checks. The checks covered parameter routing, finite objectives and gradients, six normalized active gate weights, changing weights, feasibility and hard constraints. The prespecified acceptance criteria were at least 70% feasible endpoints and changing weights in at least 50% of primary trajectories. Test brightness and comparative performance were excluded from these criteria. The main comparison comprises one discovery campaign per method.

### S5. Baseline adaptations and reproduction

All four baselines each use a shared set of 2,700 distinct highest-measured training starting sequences in a fixed shuffled order. This set differs from the multipeak initialization of GLOM–TGITO. They retain predictor-based endpoint ranking and the same final brightness and pairwise-diversity constraints. Official FLEXS AdaLead uses one round, sequence batches of 32, evaluation batches of 16, mutation parameter $\mu$=1 (per-site rate 1/221), threshold 0.05 and no recombination [5]. Its mutation generator rejects candidates outside the two-substitution radius or already visible in training, validation or calibration. Random search selects one or two positions with equal probability, samples non-anchor residues uniformly and excludes visible sequences; it is not uniform over the union of all radius-one and radius-two sequences.

The ProtHMSO comparison implements its joint-mask top-three proposal rule within this GFP radius and budget [6]. One or two random positions are masked jointly in the starting sequence; top-three joint alternatives form candidates. The GLOM and ESM2-650M variants (GLOM–ProtHMSO and ESM650–ProtHMSO) share this label-free proposal bank and rank it using their respective trained predictors. This adapts the original proposal rule to the GFP task, the project's trained predictors and endpoint selection. Each baseline receives the corresponding TGITO context's forward-sequence budget. Each method totals 1,476,063 counted fitness forwards; TGITO also performs 189,000 backward steps. Masked-language-model proposal costs are recorded separately.

Reproduction entry points are relative to the experiment root: prepare_data.py fixes partitions; position_features.py reconstructs mutation channels; train_models.py fits predictors; build_train_terrain.py builds memory; discovery_v3.py runs the discovery campaign and records its configuration and outputs; discovery_v3_baselines.py runs comparisons. Pipeline model and optimizer definitions reside under repro_project/pipeline, with encoder_contract.py, optimizer_consistency.py and fast_proposals.py supplying the runtime adapters. PROTOCOL.json, EFFECTIVE_CONFIG.json and completion records preserve parameters, source/input hashes, counts and output identities. The post-hoc score_embedding_familiarity.py uses the same search GLOM checkpoint to extract z768, with float32 features and float64 distance accumulation. Two other training seeds provide separate brightness rescoring. Neither deviation analysis nor external evidence changes frozen candidate ranks or imputes fluorescence.

### S6 Predictive results and additional controls

Tables S2 and S3 report supervised prediction metrics; Table S4 reports sequence-only adaptation controls. Together, the comparisons comprise 78 predictor evaluations, including six GLOM fits and 24 SaProt fits. Supervised neural models use the same three seeds on fixed partitions; sample SD measures variation across fits. Classical controls have one fit. All configurations are evaluated against the same test sequences and labels within each split.

GLOM branches and the regression head were initialized from scratch, with ESM2 and the Bridge frozen. Training labels were standardized using the training mean and population SD. Huber loss ($\delta = 1$) was minimized with AdamW (learning rate $3 \times 10^{-5}$, weight decay $10^{-4}$), batch size 64, BF16 autocasting and gradient-norm clipping at 1. A cosine schedule decayed the learning rate to 5% of its initial value over a 120-epoch cap. Validation Spearman selected checkpoints; training stopped after eighteen epochs without improvement. The architectural controls used the same training procedure.

Frozen ESMC-600M, ESM2-650M and ESMC-6B baselines use mean residue embeddings, excluding special tokens. The fitted MLP has hidden widths 512 and 256, train-fitted standardization, Huber loss and AdamW with learning rate $10^{-3}$, weight decay 0.01 and batch size 256. Training stops at 200 epochs or after twenty epochs without improvement. The amino-acid one-hot ridge control flattens 221 positions with twenty residue identities into 4,420 features. Ridge strengths are selected from 0.1, 1, 10, 100 and 1000. Hamming kNN selects k from 1, 4, 8, 16 and 32 with deterministic sequence-hash tie ordering; both protocols selected k=32. ESMC provenance follows the official release [7]. SaProt's frozen-MLP control uses the same head-fitting procedure on native SaProt embeddings; its three adapter controls and their input representations are described in Section S13.

ESM2-150M LoRA used rank 16, alpha 32, dropout 0.05 and query/value targets [8]. There were 1,394,433 trainable parameters. Training used microbatch 32 and accumulation 2, AdamW at $10^{-4}$, weight decay 0.01, a 25-epoch cap and patience 6. The static EVOLVEpro-style control was a 100-tree random forest on frozen ESM2-650M embeddings; it is not a reproduction of the full active-learning procedure or its original backbone setting [9].

Sequence-only ProteinTTT adapted ESM2-650M on the assay reference for 30 SGD steps, with learning rate $4\times10^{-5}$, batch 4, accumulation 4 and mask ratio 0.15; MSA use was disabled [10]. Three adaptation seeds reset the model state. An affine training-label map converted marginal scores to the fluorescence scale for error reporting. This was one-reference adaptation, not supervised or per-candidate adaptation. Wild-type marginal Spearman changed from 0.242860 without adaptation to 0.243075 ± 0.000010.

Linear probes used the five frozen representations listed in Table S5. The geometry diagnostics used fifteen intermediates on 1,000 test sequences and linear CKA [11]. The five probes include the supervised 32-dimensional joint bottleneck. Retrained architectural controls assess the contribution of the local organization.

Architectural controls test whether the same inputs suffice without GLOM's local organization. Keeping only the global branch increased random-split RMSE from 0.170 to 0.178. Removing regional–key cross-attention gave RMSE 0.175 and MAE 0.085; the full predictor reduced mean MAE by 3.3%, with the same direction in all three seeds. A matched-input Transformer reached RMSE 0.174 and MAE 0.086 (Table S2; Fig. S12a). The corresponding fixed-seed paired intervals included zero.

The branches describe different geometries. Linear CKA was 0.0023 between global and regional summaries and 0.0084 between global and key-site summaries; regional–key CKA was 0.4871 (Fig. S12c). Low CKA indicates dissimilar representations. Concatenation lowered matched ridge-probe RMSE to 0.192 from 0.293–0.429 for individual branches (Fig. S12d), supporting complementary predictive information. Figure S12b shows measured Top 100 prioritization; Tables S14 and S22 give all batch sizes.

**Table S2:** Complete supervised prediction results: random split

| **Method** | **Seeds** | **RMSE** | **MAE** | $\mathbf{R^2}$ | **Spearman** $\rho$ | **Pearson r** |
|---|---|---|---|---|---|---|
| GLOM | 3 | $0.170 \pm 0.001$ | $0.083 \pm 0.002$ | $0.892 \pm 0.001$ | $0.886 \pm 0.008$ | $0.946 \pm 0.000$ |
| GLOM: global only | 3 | $0.178 \pm 0.005$ | $0.087 \pm 0.004$ | $0.882 \pm 0.007$ | $0.872 \pm 0.012$ | $0.942 \pm 0.002$ |
| GLOM without regional–key cross-attention | 3 | $0.175 \pm 0.001$ | $0.085 \pm 0.001$ | $0.885 \pm 0.002$ | $0.888 \pm 0.000$ | $0.944 \pm 0.001$ |
| Matched-input Transformer | 3 | $0.174 \pm 0.001$ | $0.086 \pm 0.001$ | $0.886 \pm 0.001$ | $0.888 \pm 0.002$ | $0.942 \pm 0.001$ |
| One-hot Ridge | 1 | 0.221 | 0.129 | 0.816 | 0.863 | 0.904 |
| Hamming kNN | 1 | 0.385 | 0.238 | 0.443 | 0.715 | 0.792 |
| ESM2-150M + Ridge | 1 | 0.337 | 0.249 | 0.574 | 0.630 | 0.757 |
| ESM2-150M + LoRA | 3 | $0.180 \pm 0.005$ | $0.094 \pm 0.003$ | $0.879 \pm 0.006$ | $0.833 \pm 0.010$ | $0.938 \pm 0.003$ |
| ESMC-600M + MLP | 3 | $0.224 \pm 0.002$ | $0.123 \pm 0.002$ | $0.812 \pm 0.003$ | $0.831 \pm 0.006$ | $0.904 \pm 0.000$ |
| ESM2-650M + MLP | 3 | $0.247 \pm 0.002$ | $0.139 \pm 0.001$ | $0.772 \pm 0.003$ | $0.781 \pm 0.006$ | $0.881 \pm 0.001$ |
| ESMC-6B + MLP | 3 | $0.238 \pm 0.002$ | $0.134 \pm 0.002$ | $0.787 \pm 0.003$ | $0.805 \pm 0.007$ | $0.891 \pm 0.002$ |
| ESMC-600M + Ridge | 1 | 0.273 | 0.190 | 0.721 | 0.725 | 0.849 |
| ESM2-650M + Ridge | 1 | 0.297 | 0.210 | 0.669 | 0.697 | 0.818 |
| ESMC-6B + Ridge | 1 | 0.287 | 0.204 | 0.691 | 0.707 | 0.831 |
| ESM2-650M + RF | 1 | 0.409 | 0.312 | 0.372 | 0.508 | 0.622 |
| SaProt-650M, frozen MLP | 3 | $0.416 \pm 0.009$ | $0.272 \pm 0.010$ | $0.350 \pm 0.028$ | $0.495 \pm 0.029$ | $0.606 \pm 0.022$ |
| SaProt-650M, mean-pool LoRA | 3 | $0.272 \pm 0.019$ | $0.142 \pm 0.011$ | $0.722 \pm 0.040$ | $0.684 \pm 0.016$ | $0.863 \pm 0.016$ |
| SaProt-650M, CLS LoRA (AA) | 3 | $0.257 \pm 0.020$ | $0.149 \pm 0.019$ | $0.752 \pm 0.038$ | $0.760 \pm 0.020$ | $0.900 \pm 0.006$ |
| SaProt-650M, CLS LoRA (3Di) | 3 | $0.247 \pm 0.005$ | $0.148 \pm 0.007$ | $0.772 \pm 0.009$ | $0.740 \pm 0.010$ | $0.907 \pm 0.005$ |

Values are mean ± sample SD for three training seeds (20261013/14/15) on one fixed split; single-fit Ridge, kNN and RF have no SD. Random test n=3,144; depth test n=14,280 (training/validation/calibration depth $\leq 2$, test depth $\geq 3$). Supervised fitting uses training labels and validation model selection; historical panel selection is described in Section S9. RF is an EVOLVEpro-style static estimator adaptation, not the original full active-learning pipeline.

**Table S3:** Complete supervised prediction results: depth split

| Method | Seeds | RMSE | MAE | $R^2$ | Spearman $\rho$ | Pearson r |
|---|---|---|---|---|---|---|
| GLOM | 3 | $0.247 \pm 0.002$ | $0.131 \pm 0.002$ | $0.814 \pm 0.003$ | $0.870 \pm 0.004$ | $0.912 \pm 0.002$ |
| One-hot Ridge | 1 | 0.305 | 0.158 | 0.715 | 0.914 | 0.874 |
| Hamming kNN | 1 | 0.501 | 0.333 | 0.231 | 0.763 | 0.775 |
| ESMC-600M + MLP | 3 | $0.384 \pm 0.008$ | $0.238 \pm 0.004$ | $0.550 \pm 0.019$ | $0.783 \pm 0.005$ | $0.805 \pm 0.006$ |
| ESM2-650M + MLP | 3 | $0.410 \pm 0.004$ | $0.256 \pm 0.002$ | $0.487 \pm 0.009$ | $0.733 \pm 0.004$ | $0.765 \pm 0.003$ |
| ESMC-6B + MLP | 3 | $0.400 \pm 0.006$ | $0.247 \pm 0.005$ | $0.512 \pm 0.014$ | $0.762 \pm 0.010$ | $0.782 \pm 0.005$ |
| ESMC-6B + Ridge | 1 | 0.394 | 0.282 | 0.525 | 0.718 | 0.771 |
| SaProt-650M, frozen MLP | 3 | $0.556 \pm 0.003$ | $0.374 \pm 0.003$ | $0.054 \pm 0.011$ | $0.433 \pm 0.011$ | $0.487 \pm 0.013$ |
| SaProt-650M, mean-pool LoRA | 3 | $0.427 \pm 0.015$ | $0.250 \pm 0.010$ | $0.443 \pm 0.039$ | $0.646 \pm 0.010$ | $0.735 \pm 0.015$ |
| SaProt-650M, CLS LoRA (AA) | 3 | $0.324 \pm 0.052$ | $0.189 \pm 0.035$ | $0.674 \pm 0.100$ | $0.791 \pm 0.028$ | $0.856 \pm 0.031$ |
| SaProt-650M, CLS LoRA (3Di) | 3 | $0.399 \pm 0.008$ | $0.249 \pm 0.002$ | $0.512 \pm 0.021$ | $0.687 \pm 0.028$ | $0.791 \pm 0.032$ |

Values are mean ± sample SD for three training seeds (20261013/14/15) on one fixed split; single-fit Ridge, kNN and RF have no SD. Random test n=3,144; depth test n=14,280 (training/validation/calibration depth $\leq 2$, test depth $\geq 3$). Supervised fitting uses training labels and validation model selection; historical panel selection is described in Section S9. RF is an EVOLVEpro-style static estimator adaptation, not the original full active-learning pipeline.

**Table S4:** Sequence-only reference adaptation, separately evaluated

| Method / score | Seeds | RMSE | MAE | $R^2$ | Spearman $\rho$ | Pearson r |
|---|---|---|---|---|---|---|
| Unadapted ESM2: WT | 1 | 0.499 | 0.400 | 0.065 | 0.243 | 0.255 |
| Unadapted ESM2: masked | 1 | 0.516 | 0.424 | -0.000 | 0.020 | 0.023 |
| ProteinTTT: WT | 3 | $0.499 \pm 0.000$ | $0.400 \pm 0.000$ | $0.065 \pm 0.000$ | $0.243 \pm 0.000$ | $0.256 \pm 0.000$ |
| ProteinTTT: masked | 3 | $0.516 \pm 0.000$ | $0.424 \pm 0.000$ | $0.000 \pm 0.000$ | $0.021 \pm 0.000$ | $0.023 \pm 0.000$ |

Random test n=3,144. TTT adapts the reference sequence without fluorescence labels; regression errors use a training-label affine brightness map. This differs from the supervised learning setting. Three-decimal SDs rounded to 0.000 need not be exactly zero.

**Table S5:** Matched linear probes of learned GLOM representations

| Representation | Ridge $\alpha$ | RMSE | MAE | $R^2$ | Spearman $\rho$ | Pearson r |
|---|---|---|---|---|---|---|
| Global | 100 | 0.429 | 0.315 | 0.310 | 0.523 | 0.557 |
| Region | 1000 | 0.305 | 0.188 | 0.651 | 0.638 | 0.807 |
| Key-site | 1000 | 0.293 | 0.187 | 0.678 | 0.651 | 0.824 |
| Joint bottleneck | 1 | 0.167 | 0.083 | 0.896 | 0.878 | 0.947 |
| Concatenated branches | 1000 | 0.192 | 0.117 | 0.862 | 0.841 | 0.929 |

Fixed GLOM seed 20261013; random test n=3,144. Ridge $\alpha$ selected on validation Spearman. G/Rf/Kf have 256 dimensions each; concatenation 768; supervised joint bottleneck 32. The five probes use frozen features; retrained architectural controls are reported in Table S2.

### S7 Learned deviation and its statistical interpretation

For the search checkpoint, the three 256-dimensional branch summaries are concatenated and normalized. The reference bank contains only training sequences. Euclidean neighbour distance in this normalized representation is a monotone transformation of cosine similarity. The kth-neighbour distance describes local support in the learned assay-related representation.

$$D_k(x) = \frac{\|\bar{z}(x) - \bar{z}(x_{(k)})\|_2}{Q_{0.95}\{d_k(u) : u \in \mathcal{C},\ \widehat{y}(u) \geq 4.2\}}, \qquad \bar{z} = \frac{z}{\|z\|_2} \tag{S11}$$

The high-prediction reference contains 950 calibration variants selected by the fixed search prediction threshold 4.2, and its k=32 Q95 is 0.5837557862799769. All GLOM-driven Top 100 candidates meet that prediction threshold, whereas 92 of the ESM650–ProtHMSO candidates do. The same reference supplies a common geometric scale for all methods. The full calibration Q95, 0.6986382698225647, is a secondary normalization. A common rescaling preserves mean-distance ordering.

Deviation was analysed after candidate identities and rankings had been fixed. Mean deviation and Q85–Q99 comparisons were chosen for emphasis after inspecting the results; the full Q50–Q99 grid, medians and upper tails are also reported. Candidate distributions can cross, so a lower mean need not imply fewer exceedances at every threshold.

AUROC validation used the search checkpoint's residuals on measured test sequences. Across the full test, embedding distance had Spearman correlation 0.3552 with absolute error; the corresponding correlation was 0.1892 in the high-prediction subset. Q85 divided the latter into 815 inside and 158 outside observations, with large-error frequencies of approximately 6.5% and 20.3%. Table S6 gives error ratios and sample counts at all main thresholds. Q95 exceedance in measured high-prediction test variants was 49/973, or 5.04%, with Wilson interval 3.83–6.60%.

Neighbour-rank sensitivity uses k=8, 16, 32 and 64, each normalized by its own calibration Q95. GLOM–TGITO has the lowest selected-batch mean at every k. Its native confidence/diversity selection reduces mean deviation from 0.9034 in the full endpoint pool to 0.8048 in the Top 100. Other unselected pools have smaller means, and raw ESM2-150M distance gives a different ranking. These comparisons locate the observed advantage in the complete selection procedure and its task-related representation.

Measured-error AUROC was 0.773 (95% bootstrap interval 0.752–0.792) for all 3,144 test variants. Among the 973 variants predicted to have brightness $\geq 4.2$, AUROC was 0.684 (0.624–0.742) for absolute error and 0.709 (0.632–0.780) for overprediction, using an error threshold of 0.1. Calibration resampling gave Q95 exceedance-count intervals of 1–3, 1–6, 1–5, 4–6 and 18–24 for the fixed Top 100 batches of GLOM–TGITO, GLOM–AdaLead, GLOM–random, GLOM–ProtHMSO and ESM650–ProtHMSO. These intervals quantify calibration-boundary uncertainty for the fixed batches. Full resampling results are provided in Supplementary Data.

**Table S6:** Measured-error validation in the high-prediction test subset

| Q | Raw cutoff | Outside n | RMSE in | RMSE out | RMSE ratio | Large-error RR |
|---|---|---|---|---|---|---|
| Q85 | 0.507199 | 158 | 0.083 | 0.170 | 2.063 | 3.114 |
| Q90 | 0.539004 | 100 | 0.082 | 0.206 | 2.509 | 3.049 |
| Q95 | 0.583756 | 49 | 0.088 | 0.246 | 2.795 | 3.405 |
| Q97.5 | 0.611476 | 28 | 0.088 | 0.321 | 3.658 | 5.017 |
| Q99 | 0.648382 | 17 | 0.090 | 0.379 | 4.228 | 6.659 |

Thresholds fitted on 950 calibration sequences predicted $\geq$4.2 and evaluated on 973 test sequences predicted $\geq$4.2 by frozen GLOM seed 20261013. Outside means distance > cutoff. RMSE ratio = outside RMSE / inside RMSE. Large-error RR = Pr(|prediction − measured brightness| >0.1 | outside) / corresponding inside probability. At Q85 the large-error RR is 3.114.

**Table S7:** Neighborhood-size sensitivity of mean normalized distance

| Method | k=8 | k=16 | k=32 | k=64 |
|---|---|---|---|---|
| GLOM–TGITO | 0.773 | 0.784 | 0.805 | 0.813 |
| GLOM–AdaLead | 0.815 | 0.822 | 0.829 | 0.828 |
| GLOM–Random | 0.821 | 0.827 | 0.832 | 0.830 |
| GLOM–ProtHMSO | 0.802 | 0.816 | 0.831 | 0.835 |
| ESM650–ProtHMSO | 0.830 | 0.852 | 0.876 | 0.885 |

Native Top100 n=100 per method. Each column uses its own k-specific Q95 cutoff from predicted-high calibration sequences; denominators therefore differ across columns. Lower distance means greater proximity in the learned GLOM representation.

**Table S8:** All unique outputs versus native Top100 selection

| Method | Pool n | Pool mean D | Top100 D | D reduction | Pool >Q95 | Top100 >Q95 |
|---|---|---|---|---|---|---|
| GLOM–TGITO | 2594 | 0.903 | 0.805 | 10.9% | 628/2594 | 2/100 |
| GLOM–AdaLead | 2700 | 0.862 | 0.829 | 3.8% | 379/2700 | 1/100 |
| GLOM–Random | 2700 | 0.855 | 0.832 | 2.7% | 389/2700 | 4/100 |
| GLOM–ProtHMSO | 2694 | 0.857 | 0.831 | 3.1% | 375/2694 | 4/100 |
| ESM650–ProtHMSO | 2697 | 0.877 | 0.876 | 0.0% | 634/2697 | 21/100 |

D uses k32 GLOM distance / high-prediction calibration Q95. Reduction = (pool mean D − Top100 mean D) / pool mean D. Pools contain unique final outputs, and Top100 batches follow each method's native ranking and diversity constraints.

**Table S9:** Complete threshold grid: top100, high-prediction reference

| Q | Raw cutoff | TGITO | AdaLead | Random | GLOM–PH | ESM650–PH |
|---|---|---|---|---|---|---|
| Q50 | 0.3897 | 92/100 | 94/100 | 97/100 | 100/100 | 94/100 |
| Q60 | 0.4242 | 85/100 | 85/100 | 91/100 | 95/100 | 80/100 |
| Q70 | 0.4594 | 65/100 | 74/100 | 74/100 | 67/100 | 57/100 |
| Q75 | 0.4715 | 53/100 | 65/100 | 60/100 | 53/100 | 54/100 |
| Q80 | 0.4902 | 36/100 | 50/100 | 45/100 | 34/100 | 46/100 |
| Q85 | 0.5072 | 25/100 | 37/100 | 34/100 | 27/100 | 37/100 |
| Q90 | 0.5390 | 13/100 | 14/100 | 14/100 | 13/100 | 32/100 |
| Q95 | 0.5838 | 2/100 | 1/100 | 4/100 | 4/100 | 21/100 |
| Q97.5 | 0.6115 | 1/100 | 0/100 | 1/100 | 3/100 | 18/100 |
| Q99 | 0.6484 | 0/100 | 0/100 | 0/100 | 1/100 | 13/100 |

Cells are outside-count / batch or pool size; all native selections remain fixed. PH = ProtHMSO; TGITO, AdaLead and Random use GLOM. High-prediction reference n=950; all-calibration n=3,143. Distances use k=32. Quantiles were examined retrospectively. Within each method, Top6, Top30 and Top100 use the same candidate pool, ranking and constraints and form nested ordered prefixes.

### S8 Brightness risk and joint acceptance

Candidate brightness is rescored by averaging GLOM seeds 20261014 and 20261015, neither of which drove search. On the measured random test set, this evaluator has RMSE 0.164905, MAE 0.079100, $R^2$ 0.898059 and Spearman 0.896165. The common ESM2-650M evaluator reaches 0.242610, 0.134407, 0.779353 and 0.794454, respectively. Its scores provide a second predictive perspective, with lower accuracy on this assay. The two GLOM fits share architecture and training data, so their agreement measures fitting stability. Section S13 reports the SaProt prediction benchmarks.

Per-sequence cross-seed dispersion uses population standard deviation (ddof=0), distinct from the sample standard deviation across training runs. We define the optimism gap as the GLOM seed-20261013 prediction minus the mean of seeds 20261014 and 20261015. Seed 20261013 drives the four GLOM-based searches; for ESM-guided candidates, this gap is a common GLOM diagnostic rather than a comparison with the ESM search score. Table S11 reports all three predictions, the two-seed mean, mean per-sequence dispersion and mean gap.

Terrain risk R and the non-negative model–measurement gap Δ are combined as R+0.5Δ for the matched reliability analysis. Six common predicted-brightness/nearest-training-sequence strata contained low- and high-risk groups of 210 and 295 measured variants, representing 951 variants in the standardized target population. Mean predictions were 4.2419 and 4.2401. Standardized RMSE was 0.0508 versus 0.1090, with difference −0.0582 (conditional bootstrap interval −0.0982 to −0.0166). The false-bright fraction differed by −8.70 percentage points (−16.07 to −0.98).

The MaxWEnt control holds the trained mean head fixed and fits diagonal noise scales on 568,321 regression-head weights, selecting the scale using validation MSE no more than two standard errors above its baseline. The retained setting used 300 steps and scale 0.001666; 64 weight draws supplied predictive dispersion. This head-only adaptation of weight-entropy maximization [12] evaluates weight dispersion as an error diagnostic. It was useful on the full measured test but weaker than representation distance in the high-prediction regime.

The empirical lower-brightness reference is L90(x)=m(x)−q r(x), where m averages the two GLOM seeds unused in search and r=R+0.5Δ. Calibration strata are defined by m≥4.2 versus <4.2 and nearest-training Hamming distance 1 versus ≥2. Within each stratum, q is the order statistic of (m−y)/r at rank ceil(0.9(n+1)), where n is the stratum size. The same q is applied to test and candidate rows in that stratum. The risk median and dispersion Q75 use 942 calibration variants with cross-seed mean ≥4.2; this is a different prediction-based reference from the 950 variants used for embedding deviation. The full per-stratum measured coverage, widths and quantiles are in empirical_bound_test_validation.csv. Source data retain all 27 measured joint-rule settings.

The exploratory joint rule requires an empirical lower-brightness reference ≥4.2, combined risk ≤0.1576107955, cross-seed SD ≤0.0145756602 and nearest-visible-sequence Hamming distance ≥2. The risk and dispersion thresholds are the high-prediction calibration median and Q75. The resulting counts are 38, 16, 14, 14 and 11 for the five Top 100 batches. This secondary analysis reports the full threshold grid. The empirical lower reference uses calibration residuals scaled by terrain-derived risk; it is not a distribution-free conformal guarantee for optimized designs [13].

Source data additionally report fixed-backbone ProteinMPNN scores [14] and preservation of the canonical chromophore segment as structural compatibility checks. Between-method differences in ProteinMPNN scores were small.

**Table S10:** Common-evaluator accuracy on measured test sequences

| Evaluator | Scope | n | RMSE | MAE | $R^2$ | Spearman $\rho$ |
|---|---|---|---|---|---|---|
| GLOM cross-seed | All test | 3144 | 0.165 | 0.079 | 0.898 | 0.896 |
| GLOM cross-seed | Measured ≥4.2 | 850 | 0.067 | 0.039 | -0.811 | 0.596 |
| GLOM cross-seed | Predicted ≥4.2 | 966 | 0.096 | 0.046 | 0.141 | 0.601 |
| GLOM cross-seed | Nearest train: 1 | 1802 | 0.121 | 0.057 | 0.928 | 0.878 |
| GLOM cross-seed | Nearest train: ≥2 | 1342 | 0.210 | 0.109 | 0.863 | 0.898 |
| ESM2-650M | All test | 3144 | 0.243 | 0.134 | 0.779 | 0.794 |
| ESM2-650M | Measured ≥4.2 | 850 | 0.131 | 0.079 | -5.838 | 0.373 |
| ESM2-650M | Predicted ≥4.2 | 616 | 0.121 | 0.057 | 0.108 | 0.526 |
| ESM2-650M | Nearest train: 1 | 1802 | 0.167 | 0.088 | 0.863 | 0.786 |
| ESM2-650M | Nearest train: ≥2 | 1342 | 0.317 | 0.197 | 0.687 | 0.774 |

Evaluators average seeds 20261014/15, which did not drive the search. Predicted-high subsets use each evaluator's own ≥4.2 predictions and differ in membership. Nearest-training categories use Hamming solely as a sequence-distance stratum. Restricted-range $R^2$ can be negative.

**Table S11:** GLOM predictions, cross-seed dispersion and optimism gaps on native selections

| Method | Top n | Seed 13 | Seed 14 | Seed 15 | Cross mean | Mean seed SD | Mean gap |
|---|---|---|---|---|---|---|---|
| GLOM–TGITO | 6 | 4.461 | 4.427 | 4.424 | 4.426 | 0.020 | 0.035 |
| GLOM–TGITO | 30 | 4.448 | 4.401 | 4.412 | 4.407 | 0.022 | 0.041 |
| GLOM–TGITO | 100 | 4.434 | 4.385 | 4.397 | 4.391 | 0.019 | 0.043 |
| GLOM–AdaLead | 6 | 4.485 | 4.437 | 4.414 | 4.426 | 0.020 | 0.059 |
| GLOM–AdaLead | 30 | 4.464 | 4.419 | 4.411 | 4.415 | 0.017 | 0.049 |
| GLOM–AdaLead | 100 | 4.443 | 4.399 | 4.401 | 4.400 | 0.017 | 0.043 |
| GLOM–Random | 6 | 4.493 | 4.422 | 4.421 | 4.421 | 0.019 | 0.072 |
| GLOM–Random | 30 | 4.470 | 4.412 | 4.424 | 4.418 | 0.019 | 0.052 |
| GLOM–Random | 100 | 4.449 | 4.396 | 4.406 | 4.401 | 0.018 | 0.048 |
| GLOM–ProtHMSO | 6 | 4.501 | 4.410 | 4.427 | 4.419 | 0.011 | 0.082 |
| GLOM–ProtHMSO | 30 | 4.477 | 4.411 | 4.433 | 4.422 | 0.016 | 0.055 |
| GLOM–ProtHMSO | 100 | 4.453 | 4.403 | 4.425 | 4.414 | 0.016 | 0.040 |
| ESM650–ProtHMSO | 6 | 4.288 | 4.259 | 4.291 | 4.275 | 0.021 | 0.013 |
| ESM650–ProtHMSO | 30 | 4.294 | 4.286 | 4.308 | 4.297 | 0.032 | -0.003 |
| ESM650–ProtHMSO | 100 | 4.304 | 4.306 | 4.328 | 4.317 | 0.021 | -0.012 |

Seeds 13/14/15 abbreviate 20261013/20261014/20261015. Predictions and gaps are batch means on the log-fluorescence scale. Cross mean averages seeds 14 and 15; gap is seed 13 minus that mean. Seed 13 is the search predictor for GLOM-based methods and a common GLOM evaluator for ESM650–ProtHMSO. Mean seed SD averages per-sequence population SD between seeds 14 and 15, not uncertainty in a batch mean. Each method has one campaign; the three batch sizes are nested selections.

### S9 Historical design provenance and selected candidate sequences

The local panels predate the present splits and were informed by ridge analysis of the original assay. The final fifteen sites occur in the saved top-thirty ranking; the later consolidation rule is not fully documented. Supplementary Data 2 provides the sample reconstructed from the archived selection script and input data, together with source hashes. This reconstruction is not a contemporaneous execution record.

Historical policy tuning used measured target brightness and neighbourhood coverage (Section S4). The present campaign uses these parameters with newly fitted GLOM models and training-only memories. Although predictor partitions are sequence-disjoint, historical panel and policy selection used assay data overlapping the current tests; the evaluation is therefore retrospective. Supplementary Data 2 documents the overlap. Nested panel and policy selection, full-chain multiterrain ablation and cross-protein evaluation are needed to test generalization.

The formal discovery campaign's native-ranked Top 30 were fixed before biophysical evaluation. IDs C01–C30 follow that confidence ranking; Supplementary Data 1 gives every complete sequence, its actual start and both sets of substitutions. C12, C14, C21, C22, C27 and C30 were selected after the trajectory analysis to illustrate surface, contact-network and torsional changes. These examples are not a newly ranked Top 6. Table S12 records their sequence identities and search-model predictions; Section S14 reports the complete cohort.

**Table S12:** Six representative sequences from the native Top 30

| ID | Substitutions relative to WT | Changes from actual start | Predicted $\log_{10}$ F | Confidence |
|---|---|---|---|---|
| C12 | E105D; E122K; L199H; A219D | E122K; A219D | 4.4351 | 0.7505 |
| C14 | Q30R; K36N; G81S; D129G; G206E | K36N; D129G | 4.4338 | 0.7461 |
| C21 | M3I; T27S; Y49F; G206E | M3I; G206E | 4.4652 | 0.7369 |
| C22 | V168L; L199H | V126I; V168L | 4.4142 | 0.7363 |
| C27 | E70V; V95I; F167V; L199H; I218E | F167V; V218E | 4.5048 | 0.7313 |
| C30 | K36E; E122A; L149M; G206E | E122A; G206E | 4.4275 | 0.7252 |

IDs retain the original native confidence ranks. Predictions refer to the final discrete sequences. C22 contains a reversion to the WT residue at position 126; C27 changes V218 in its start to E218, whereas the WT residue is I218. Complete Top 30 and start sequences are supplied in Data 1.

**Table S12:** Representative Top 30 sequences: full amino-acid sequences (continued)

**C12** E105D; E122K; L199H; A219D

```
PAMKIECRITGTLNGVEFELVGGGEGTPEQGRMTNKMKSTKGALTFSPYL
LSHVMGYGFYHFGTYPSGYENPFLHAINNGGYTNTRIEKYEDGGVLHVSF
SYRYDAGRVIGDFKVVGTGFPKDSVIFTDKIIRSNATVEHLHPMGDNVLV
GSFARTFSLRDGGYYSFVVDSHMHFKSAIHPSILQNGGPMFAFRRVEEHH
SNTELGIVEYQHAFKTPIDFA
```

**C14** Q30R; K36N; G81S; D129G; G206E

```
PAMKIECRITGTLNGVEFELVGGGEGTPERGRMTNNMKSTKGALTFSPYL
LSHVMGYGFYHFGTYPSGYENPFLHAINNGSYTNTRIEKYEDGGVLHVSF
SYRYEAGRVIGDFKVVGTGFPEDSVIFTGKIIRSNATVEHLHPMGDNVLV
GSFARTFSLRDGGYYSFVVDSHMHFKSAIHPSILQNGGPMFAFRRVEELH
SNTELEIVEYQHAFKTPIAFA
```

**C21** M3I; T27S; Y49F; G206E

```
PAIKIECRITGTLNGVEFELVGGGEGSPEQGRMTNKMKSTKGALTFSPFL
LSHVMGYGFYHFGTYPSGYENPFLHAINNGGYTNTRIEKYEDGGVLHVSF
SYRYEAGRVIGDFKVVGTGFPEDSVIFTDKIIRSNATVEHLHPMGDNVLV
GSFARTFSLRDGGYYSFVVDSHMHFKSAIHPSILQNGGPMFAFRRVEELH
SNTELEIVEYQHAFKTPIAFA
```

All sequences use the 221-residue project reference and retain the G56–Y57–G58 chromophore-forming motif. The CSV and FASTA records in Supplementary Data 1 contain the same sequences.

**Table S12:** Representative Top 30 sequences: full amino-acid sequences (continued)

**C22** V168L; L199H

```
PAMKIECRITGTLNGVEFELVGGGEGTPEQGRMTNKMKSTKGALTFSPYL
LSHVMGYGFYHFGTYPSGYENPFLHAINNGGYTNTRIEKYEDGGVLHVSF
SYRYEAGRVIGDFKVVGTGFPEDSVIFTDKIIRSNATVEHLHPMGDNVLV
GSFARTFSLRDGGYYSFLVDSHMHFKSAIHPSILQNGGPMFAFRRVEEHH
SNTELGIVEYQHAFKTPIAFA
```

**C27** E70V; V95I; F167V; L199H; I218E

```
PAMKIECRITGTLNGVEFELVGGGEGTPEQGRMTNKMKSTKGALTFSPYL
LSHVMGYGFYHFGTYPSGYVNPFLHAINNGGYTNTRIEKYEDGGILHVSF
SYRYEAGRVIGDFKVVGTGFPEDSVIFTDKIIRSNATVEHLHPMGDNVLV
GSFARTFSLRDGGYYSVVVDSHMHFKSAIHPSILQNGGPMFAFRRVEEHH
SNTELGIVEYQHAFKTPEAFA
```

**C30** K36E; E122A; L149M; G206E

```
PAMKIECRITGTLNGVEFELVGGGEGTPEQGRMTNEMKSTKGALTFSPYL
LSHVMGYGFYHFGTYPSGYENPFLHAINNGGYTNTRIEKYEDGGVLHVSF
SYRYEAGRVIGDFKVVGTGFPADSVIFTDKIIRSNATVEHLHPMGDNVMV
GSFARTFSLRDGGYYSFVVDSHMHFKSAIHPSILQNGGPMFAFRRVEELH
SNTELEIVEYQHAFKTPIAFA
```

All sequences use the 221-residue project reference and retain the G56–Y57–G58 chromophore-forming motif. The CSV and FASTA records in Supplementary Data 1 contain the same sequences.

### S10 Visual provenance and reproducibility

Main Fig. 2 shows a regional view of one recorded climb; Supplementary Fig. S3 compares measured and predicted surfaces from the 22,003-sequence training memory. The surfaces use 35-neighbour Gaussian averaging, linear interpolation on 220 × 220 grids and Gaussian smoothing with $\sigma$=2 cells; cells outside the data convex hull are masked. Colours follow the empirical distribution of training fluorescence, with red beginning at its median, 4.1303; elevation remains linear in $\log_{10}$ fluorescence. These surfaces summarize the representation geometry. Online search evaluates its potential in the original high-dimensional features.

The example is context 0054, selected by native Top 6 rank 1 before visual inspection. Its twelve saved states span steps 0–70, with four active-source-set changes and four accepted discrete proposals. In each view, neighbour-weighted UMAP coordinates locate the relaxed queries, and path heights follow that view's display surface. The corresponding hard sequence is the amino-acid argmax. Main Fig. 2A marks the initial state and accepted-proposal checkpoints at steps 0, 6, 12, 18 and 24; Fig. 2B shows the joint objective evaluated on the hard sequence at all twelve checkpoints. The sequence reached at step 24 persists through step 70. Lines connect the recorded states. Supplementary Fig. S10 provides the other views, all saved states, predictions and gate weights.

Raw coordinates, measurements, predictions, states and display grids accompany the figures. Supplementary Figs S6–S8 report endpoint source weights, computational details and additional threshold and measured-error analyses, respectively.

Main Fig. 5 uses the frozen Top 30 biophysical summaries. The structural image is an actual C27 production conformation selected from frames with simultaneous Thr137/Arg155 phenolate contacts. Supplementary Fig. S13 uses, for each representative, the conformation closest to the aligned time-average of its C-alpha coordinates. Frame indices and selection criteria are recorded in Supplementary Data 3.

The LaTeX source archive contains manuscript sources and figures; Supplementary Data contain numerical plotting inputs and protocol snapshots. SHA256 records connect these inputs to the frozen experiment reports. Training and search scripts, model weights and large feature arrays are retained in the experiment archive. Tables S19–S22 report the SaProt settings, individual fits, paired error intervals and measured-protein prioritization results.

### S11 Additional evidence for prediction and complete-framework discovery

Tables S13–S17 report batch-size sensitivity, measured-protein prioritization, joint-rule counts and matched-error validation. Screening metrics cover all 78 predictor evaluations. The 405 discovery joint-rule combinations are evaluated on the fixed candidate batches using their per-sequence scores.

**Table S13:** Complete-pipeline evidence at three native batch sizes

| Method | Batch n | Mean D | Predicted brightness | Terrain risk | Support | Diversity | Joint pass |
|---|---|---|---|---|---|---|---|
| GLOM–TGITO | 6 | 0.7792 | 4.426 | 0.0983 | 0.3824 | 4.87 | 2 |
| GLOM–AdaLead | 6 | 0.8347 | 4.426 | 0.1246 | 0.2744 | 6.13 | 0 |
| GLOM–random | 6 | 0.8356 | 4.421 | 0.1203 | 0.3024 | 7.93 | 1 |
| GLOM–ProtHMSO | 6 | 0.8567 | 4.419 | 0.1327 | 0.2576 | 5.27 | 0 |
| ESM650–ProtHMSO | 6 | 1.1314 | 4.275 | 0.1570 | 0.1836 | 7.00 | 1 |
| GLOM–TGITO | 30 | 0.7815 | 4.407 | 0.1074 | 0.3544 | 7.06 | 7 |
| GLOM–AdaLead | 30 | 0.8566 | 4.415 | 0.1286 | 0.2606 | 6.60 | 3 |
| GLOM–random | 30 | 0.8275 | 4.418 | 0.1300 | 0.2545 | 6.43 | 3 |
| GLOM–ProtHMSO | 30 | 0.8209 | 4.422 | 0.1218 | 0.2876 | 5.92 | 5 |
| ESM650–ProtHMSO | 30 | 0.9343 | 4.297 | 0.1375 | 0.2070 | 6.61 | 2 |
| GLOM–TGITO | 100 | 0.8048 | 4.391 | 0.1182 | 0.3168 | 7.62 | 38 |
| GLOM–AdaLead | 100 | 0.8290 | 4.400 | 0.1356 | 0.2397 | 6.66 | 16 |
| GLOM–random | 100 | 0.8323 | 4.401 | 0.1312 | 0.2441 | 6.72 | 14 |
| GLOM–ProtHMSO | 100 | 0.8310 | 4.414 | 0.1260 | 0.2711 | 6.09 | 14 |
| ESM650–ProtHMSO | 100 | 0.8763 | 4.317 | 0.1353 | 0.2148 | 6.38 | 11 |

Each row describes the frozen native batch, without reranking. D uses the same k32 GLOM representation and high-prediction calibration Q95 denominator. Brightness averages two unused GLOM seeds. Diversity is mean pairwise Hamming distance; it is not the embedding familiarity score. Joint pass uses the fixed $\geq$2 substitutions, lower brightness $\geq$4.2, median-risk and Q75-dispersion rule. The three batch sizes are nested selections from one campaign per method.

For each fitted predictor, stable descending prediction order defines the Top 6/30/100 measured test variants. We summarize measured brightness, fractions with brightness at or above 4.2 and 4.35, and enrichment for the measured top decile. Enrichment divides the selected top-decile fraction by its prevalence in the full test set. Table S14 compares SaProt with the other predictors; Table S22 provides all SaProt batch sizes with seed variation. These retrospective tests use measured test variants, not generated candidates.

**Table S14:** Retrospective prioritization of measured test proteins

| Split | Predictor | Top6 mean | Top30 mean | Top100 mean | Top100 ≥4.2 | Top100 ≥4.35 | Top10% enrichment |
|---|---|---|---|---|---|---|---|
| Random | GLOM | 4.379 | 4.340 | 4.324 | 96.3% | 34.7% | 8.62 |
| Random | GLOM global branch only | 4.364 | 4.352 | 4.314 | 95.7% | 34.0% | 7.79 |
| Random | GLOM without regional–key cross-attention | 4.357 | 4.350 | 4.319 | 96.3% | 32.0% | 8.72 |
| Random | Matched-input Transformer | 4.369 | 4.355 | 4.318 | 96.7% | 35.0% | 8.35 |
| Random | One-hot ridge | 4.037 | 4.230 | 4.283 | 89.0% | 33.0% | 7.39 |
| Random | ESM2-150M + LoRA | 4.347 | 4.324 | 4.299 | 92.3% | 26.3% | 7.12 |
| Random | ESMC-600M + MLP | 4.366 | 4.347 | 4.309 | 95.3% | 32.0% | 8.12 |
| Random | ESM2-650M + MLP | 4.375 | 4.328 | 4.291 | 93.0% | 30.3% | 7.39 |
| Random | ESMC-6B + MLP | 4.345 | 4.313 | 4.283 | 88.0% | 28.3% | 7.29 |
| Random | Hamming kNN | 4.378 | 4.288 | 4.262 | 88.0% | 26.0% | 6.99 |
| Random | ESM2-150M + Ridge | 4.041 | 4.094 | 4.152 | 44.0% | 7.0% | 2.00 |
| Random | ESM2-650M + RF | 4.316 | 4.233 | 4.203 | 62.0% | 5.0% | 2.69 |
| Random | ESMC-600M + Ridge | 4.171 | 4.131 | 4.187 | 57.0% | 4.0% | 2.30 |
| Random | ESM2-650M + Ridge | 4.111 | 4.121 | 4.174 | 52.0% | 6.0% | 2.60 |
| Random | ESMC-6B + Ridge | 4.221 | 4.229 | 4.207 | 56.0% | 10.0% | 3.29 |
| Random | SaProt-650M, frozen MLP | 4.225 | 4.199 | 4.179 | 58.3% | 7.7% | 3.39 |

Sequences are ordered by each fitted model's test prediction, with stable ties; brightness summaries use the measured labels of that held-out test catalogue. Neural entries average three training seeds, and classical predictors have one fit. Enrichment is the fraction of selected sequences in the measured top decile divided by its prevalence in the full test set. These are predictor-screening results, distinct from newly generated proteins. Seed-level source data include all marginal-score/ ProteinTTT controls. No discovery confidence/diversity filter is applied.

**Table S14:** Retrospective prioritization of measured test proteins (continued)

| Split | Predictor | Top6 mean | Top30 mean | Top100 mean | Top100 ≥4.2 | Top100 ≥4.35 | Top10% enrichment |
|---|---|---|---|---|---|---|---|
| Random | SaProt-650M, mean-pool LoRA | 4.285 | 4.237 | 4.227 | 64.7% | 10.0% | 3.29 |
| Random | SaProt-650M, CLS LoRA (AA) | 4.210 | 4.202 | 4.204 | 52.0% | 3.7% | 1.63 |
| Random | SaProt-650M, CLS LoRA (3Di) | 4.290 | 4.276 | 4.223 | 64.7% | 11.3% | 3.39 |
| Depth | GLOM | 4.348 | 4.334 | 4.331 | 97.3% | 42.3% | 9.27 |
| Depth | One-hot ridge | 4.297 | 4.338 | 4.340 | 95.0% | 54.0% | 9.00 |
| Depth | ESMC-600M + MLP | 4.299 | 4.307 | 4.283 | 89.3% | 33.0% | 8.03 |
| Depth | ESM2-650M + MLP | 4.314 | 4.287 | 4.273 | 79.7% | 31.3% | 7.13 |
| Depth | ESMC-6B + MLP | 4.323 | 4.292 | 4.281 | 86.0% | 30.7% | 7.57 |
| Depth | Hamming kNN | 4.353 | 4.247 | 4.279 | 89.0% | 31.0% | 8.20 |
| Depth | ESMC-6B + Ridge | 3.882 | 3.979 | 4.050 | 34.0% | 8.0% | 2.00 |
| Depth | SaProt-650M, frozen MLP | 3.951 | 4.055 | 4.044 | 48.0% | 11.7% | 3.87 |
| Depth | SaProt-650M, mean-pool LoRA | 4.245 | 4.232 | 4.171 | 61.7% | 12.7% | 4.40 |
| Depth | SaProt-650M, CLS LoRA (AA) | 4.218 | 4.187 | 4.180 | 42.3% | 4.7% | 2.33 |
| Depth | SaProt-650M, CLS LoRA (3Di) | 4.218 | 4.179 | 4.181 | 52.3% | 11.3% | 3.63 |

Sequences are ordered by each fitted model's test prediction, with stable ties; brightness summaries use the measured labels of that held-out test catalogue. Neural entries average three training seeds, and classical predictors have one fit. Enrichment is the fraction of selected sequences in the measured top decile divided by its prevalence in the full test set. These are predictor-screening results, distinct from newly generated proteins. Seed-level source data include all marginal-score/ ProteinTTT controls. No discovery confidence/diversity filter is applied.

The joint criterion combines sequence novelty with the existing calibration-based reliability rule. In the Top 100, novelty-only counts at distance ≥2 are 90/44/42/20/27; reliability-only counts are 42/46/45/52/42; their intersections are 38/16/14/14/11 in the order GLOM–TGITO, GLOM–AdaLead, GLOM–random, GLOM–ProtHMSO and ESM650–ProtHMSO. The joint criterion therefore identifies more sequence-novel candidates meeting the specified reliability thresholds.

**Table S15:** Joint novelty–reliability counts across the complete brightness–risk grid

| Lower brightness | Risk cutoff | GLOM–TGITO | GLOM–AdaLead | GLOM–random | GLOM–ProtHMSO | ESM650–ProtHMSO |
|---|---|---|---|---|---|---|
| 4.2 | Q25 | 35 | 15 | 11 | 13 | 11 |
| 4.2 | Q50 | 38 | 16 | 14 | 14 | 11 |
| 4.2 | Q75 | 39 | 20 | 16 | 14 | 11 |
| 4.3 | Q25 | 25 | 10 | 3 | 9 | 3 |
| 4.3 | Q50 | 26 | 10 | 3 | 10 | 3 |
| 4.3 | Q75 | 26 | 12 | 4 | 10 | 3 |
| 4.35 | Q25 | 3 | 2 | 1 | 5 | 0 |
| 4.35 | Q50 | 3 | 2 | 1 | 5 | 0 |
| 4.35 | Q75 | 3 | 2 | 1 | 5 | 0 |

Native Top100, at least two substitutions from every visible sequence; cross-seed SD cutoff is the high-prediction calibration Q75 (0.01457566). Lower brightness is the terrain-risk-scaled empirical 90% lower reference. Risk cutoffs Q25/Q50/Q75 are fixed from calibration data. This complete 3×3 grid includes the stringent 4.35 setting, where GLOM–ProtHMSO retains more candidates. The source archive additionally contains all 405 combinations of method, batch size, novelty distance, brightness and risk. Novelty distance is a sequence-discovery criterion, not a familiarity metric.

**Table S16:** Measured-test validation of the joint reliability rule

| Min. distance to train | Risk cutoff | Accepted n | Measured ≥4.2 | Fraction | Wilson 95% interval |
|---|---|---|---|---|---|
| 1 | Q25 | 56 | 55 | 98.2% | 90.6–99.7% |
| 1 | Q50 | 80 | 78 | 97.5% | 91.3–99.3% |
| 1 | Q75 | 101 | 99 | 98.0% | 93.1–99.5% |
| 2 | Q25 | 10 | 10 | 100.0% | 72.2–100.0% |
| 2 | Q50 | 14 | 14 | 100.0% | 78.5–100.0% |
| 2 | Q75 | 16 | 16 | 100.0% | 80.6–100.0% |

Lower-brightness reference ≥4.2 and the same Q75 dispersion cutoff are used for every row. Measured cases use nearest-training distance, whereas generated-candidate novelty uses training+validation+calibration; these reference sets differ. The fixed Q50 rule accepts 80 measured variants (78 bright), or 14 when requiring distance ≥2 (all 14 bright). Intervals describe the measured test selections. The source CSV retains all 27 settings, including zero-acceptance cases.

**Table S17:** Measured errors at matched predicted brightness and sequence-distance strata

| Risk group | Observed n | Mean prediction | RMSE | RMSE 95% interval | False-bright fraction |
|---|---|---|---|---|---|
| Low | 210 | 4.2419 | 0.0508 | 0.0451–0.0564 | 27.0% |
| High | 295 | 4.2401 | 0.1090 | 0.0668–0.1484 | 35.7% |

Six shared strata of predicted brightness and nearest-training Hamming distance are standardized to a target population of 951 measured variants. Low/high risk groups contain 210/295 observed variants. Combined risk is gate-weighted terrain risk plus 0.5 times the gate-weighted positive relative-brightness gap defined in Section S2. The false-bright fraction uses measured brightness <4.2. Low-minus-high RMSE difference is −0.0582 (95% conditional stratified-bootstrap interval −0.0982 to −0.0166); the false-bright difference is −8.70 percentage points (−16.07 to −0.98). The 2,000 resamples quantify conditional measured-data uncertainty.

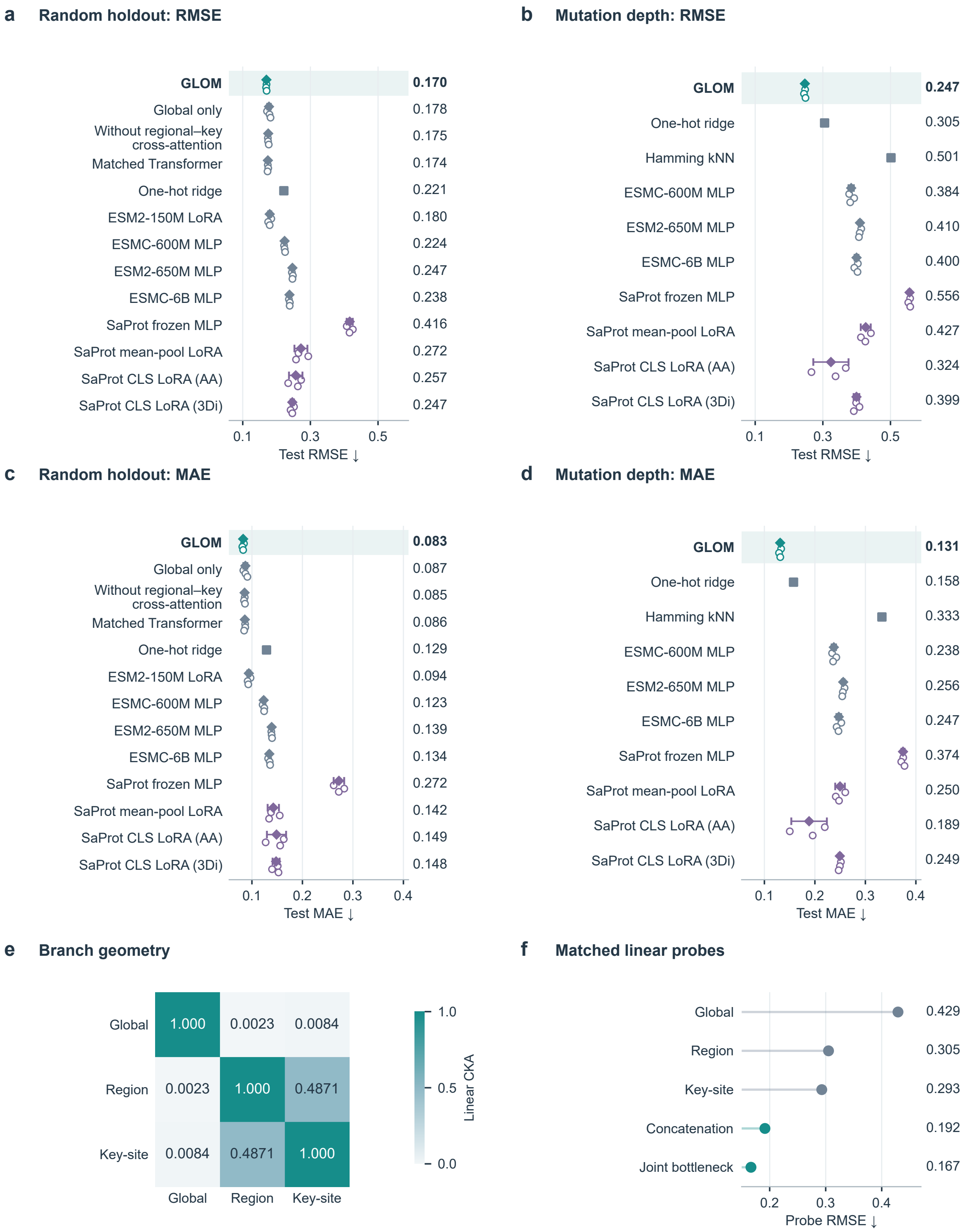


**Figure S2: Predictive accuracy and representation controls**. (a,b) Test RMSE and (c,d) test MAE on random holdout and mutation-depth extrapolation, respectively. Each metric uses the same test sequences and fitted models. Open circles are individual training seeds, offset vertically for visibility; diamonds and intervals show their mean and sample standard deviation. Squares denote one classical fit. Printed values are means. (e) Linear centered kernel alignment (CKA) of the global, regional and key-site branches on 1,000 test sequences. (f) Test RMSE of matched ridge probes on the three branches, their 768-dimensional concatenation and the supervised 32-dimensional joint bottleneck. Panels e,f use GLOM seed 20261013. The bottleneck follows supervised fusion. The four SaProt configurations use the 650M base; CLS indicates the official-recipe adaptation. Historically assay-informed local panels remain fixed, as described in the Methods.

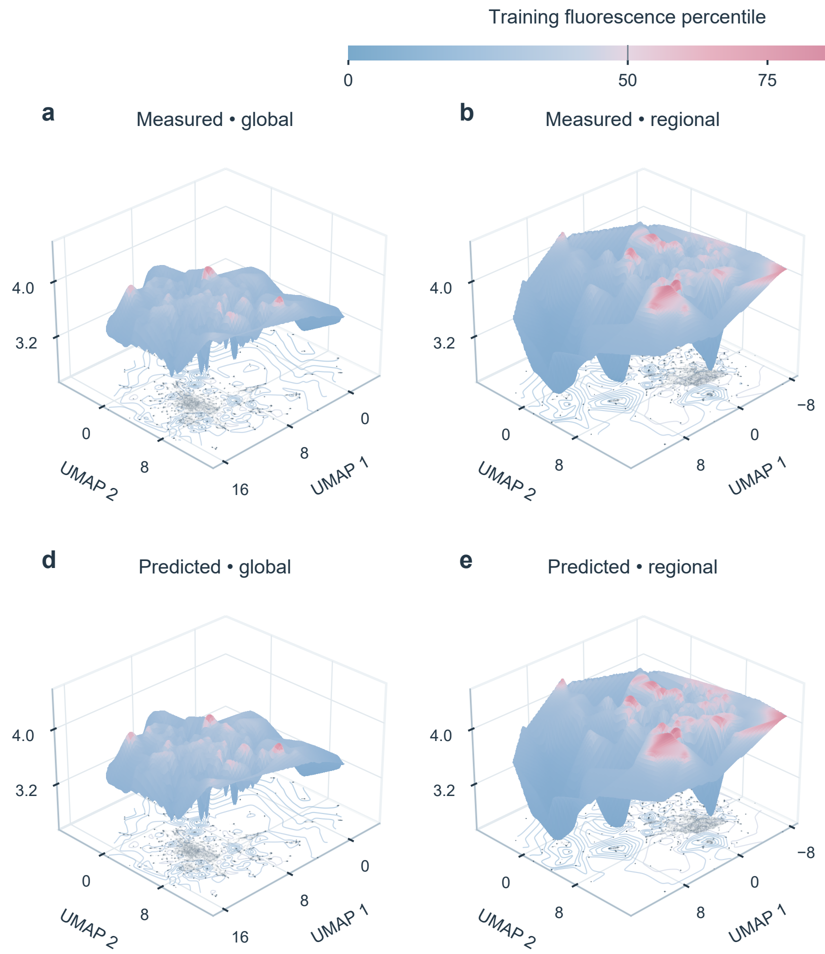


**Figure S3: Measured and predicted landscapes of the formal training memory**. a–c, measured log10 fluorescence; d–f, model-predicted log10 fluorescence, shown at the same 22,003 original training-only memory coordinates. Columns show global max pooling, regional mean pooling and flattened key-site representations. Every measured and predicted panel shares the same numeric height and color scale. The display uses 35-neighbor Gaussian-weighted averaging of actual point values, linear interpolation on a 220×220 grid and Gaussian smoothing with sigma 2 grid cells; cells outside the observed convex hull are masked. Faint base-plane points preserve all original saved UMAP coordinates. Surfaces interpolate the training observations within the convex hull. Exact unsmoothed observations, predictions, row IDs and display grids are archived with the figure. The planes display representation geometry; online potentials are evaluated in the original feature spaces. All fluorescence surfaces use the same blue-to-red training-percentile scale. Red begins at the training median (4.1303); elevation remains linear in $\log_{10}$ fluorescence.

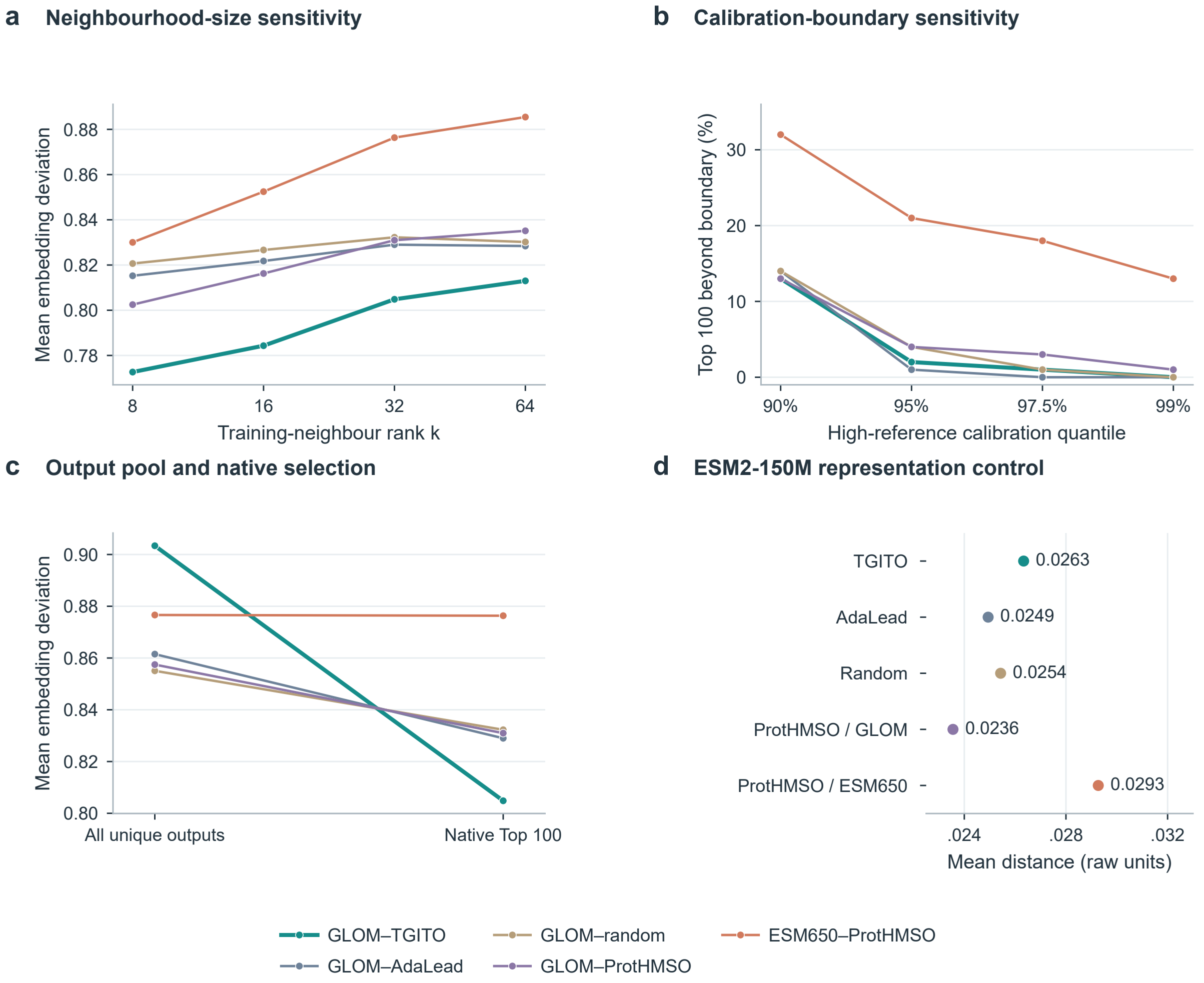


**Figure S4: Neighbourhood sensitivity and selection controls**. (a) Mean embedding deviation at neighbour ranks 8, 16, 32 and 64. (b) Native Top 100 proportions beyond four secondary calibration boundaries at k=32. (c) Mean deviation in the full unique output pool and native Top 100. (d) Mean k=32 distance in the frozen ESM2-150M representation, without calibration scaling. Distance is Euclidean distance to the kth training neighbour after per-vector L2 normalization; deviation divides that distance by the specified calibration quantile. Panels a–c use the calibration subset with predicted log-fluorescence $\geq$4.2; a and c use its q95 distance for scaling. Lines join aggregate results at the indicated settings.

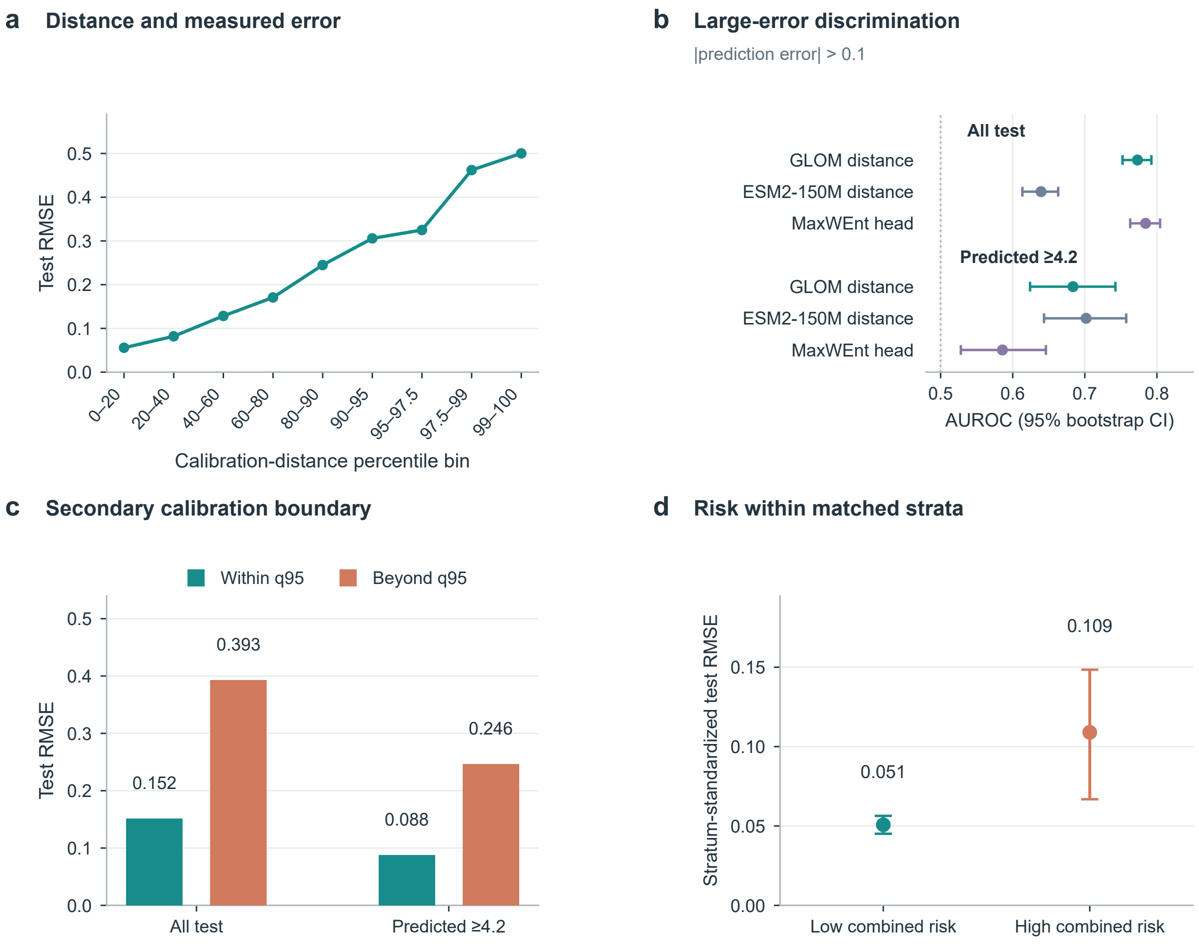


**Figure S5: Additional measured-error checks**. (a) GLOM test RMSE in calibration-distance percentile bins, with 644, 620, 636, 615, 330, 156, 88, 27 and 28 variants from left to right. (b) AUROC for absolute prediction errors greater than 0.1, with 95% bootstrap confidence intervals, on all measured test sequences (n=3,144) and the predicted-high subset (n=973). (c) Test RMSE within and beyond the q95 secondary boundary at k=32; group sizes are 3,001/143 for all test variants and 924/49 for the predicted-high subset. (d) Stratum-standardized test RMSE in low- and high-combined-risk groups (n=210/295), with bootstrap intervals. Combined risk is $\bar{R} + 0.5\bar{\Delta}$ (Section S3); mean predictions are 4.242 and 4.240 for the low- and high-risk groups, respectively. Distances are kth-neighbour Euclidean distances after L2 normalization. Panels a–c use prediction errors from the search checkpoint. Panel d uses cross-seed prediction errors and conditional stratified bootstrap intervals; its residuals are a separate analysis. Errors are on the $\log_{10}$ fluorescence scale.

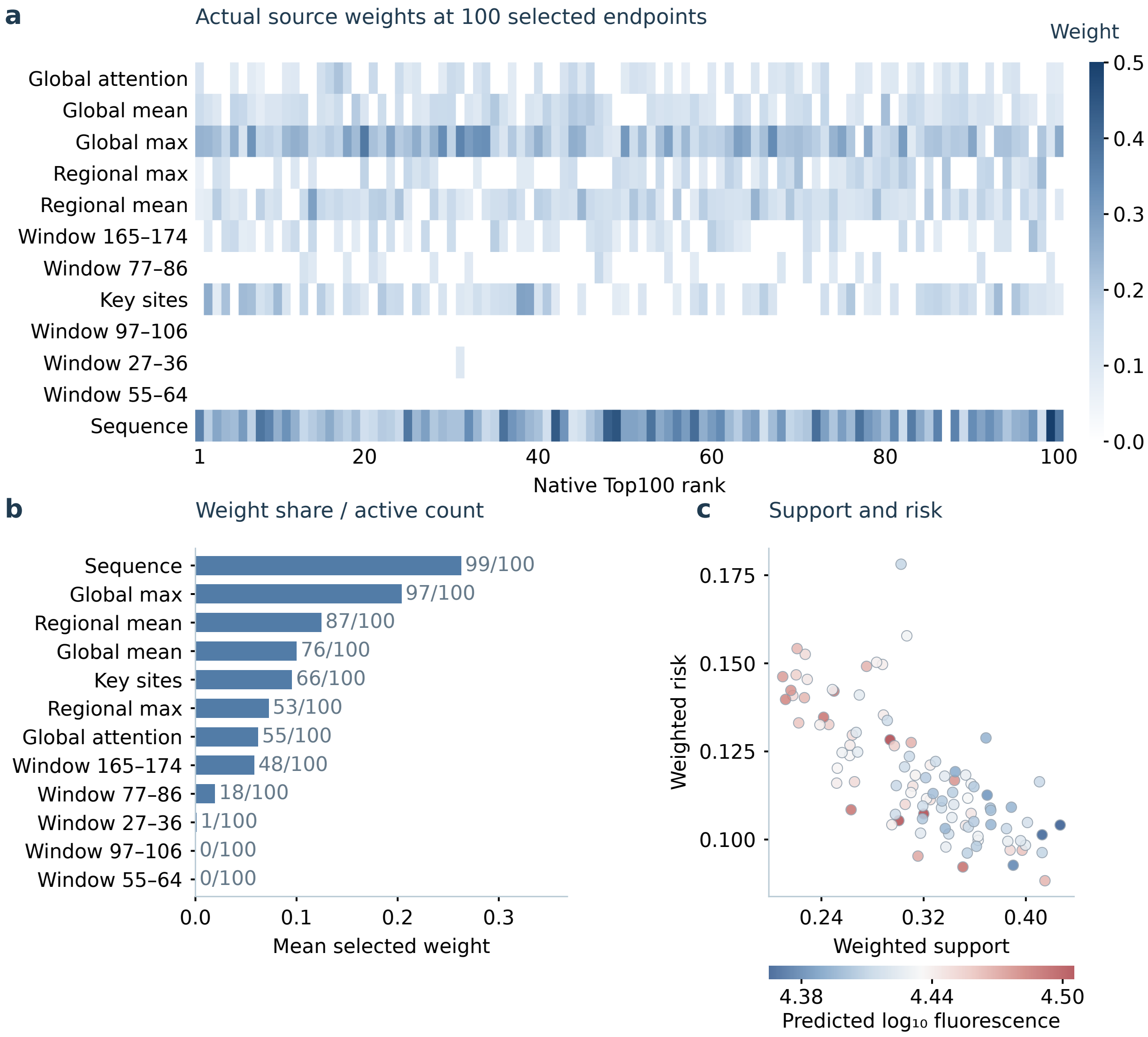


**Figure S6: Source contributions at the selected endpoints**. Source selection at the 100 frozen native endpoints of the GLOM–TGITO discovery campaign. a, each column corresponds to a candidate in native rank order, with the twelve actually recorded source weights; exactly six are nonzero and all twelve sum to one. b, mean source-weight share, with labels giving the number of endpoints at which the source was active. c, saved weighted support and risk, colored by the search-model prediction.

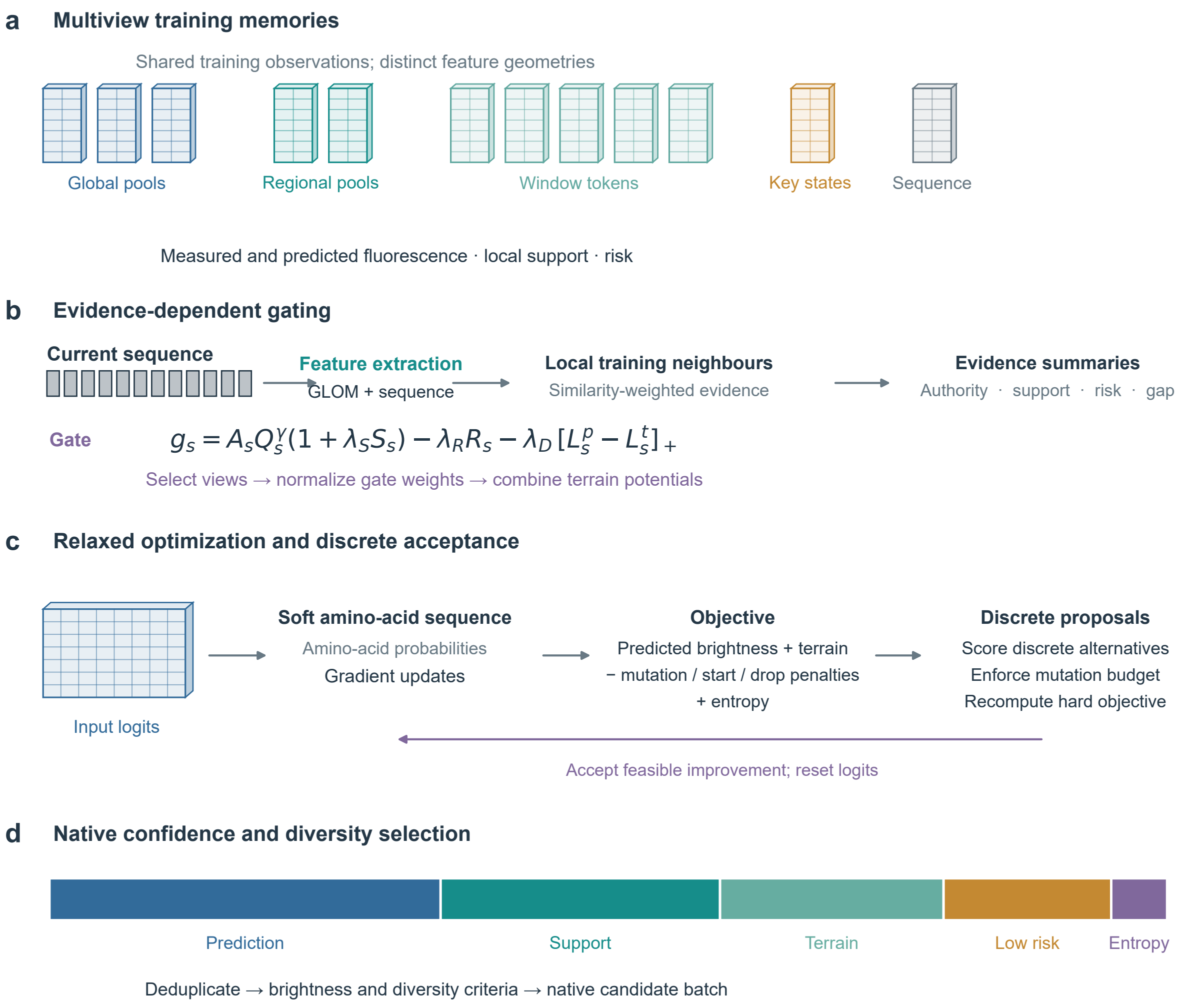


**Figure S7: Computational path from training memories to native selection.** a, Twelve source memories share the same 22,003 measured training sequences. Three global pools, two post-attention regional pools, five pre-compression window descriptors, flattened pre-attention key states and a fixed projected sequence representation provide their geometries. Tensor glyphs are schematic. b, Online candidate features retrieve 32 cosine neighbours per source. Weighted local evidence, calibrated source quality and prediction–measurement disagreement determine a six-of-twelve gate. Gradients follow the original feature-space queries. c, Amino-acid-logit updates alternate with hard substitution proposals; accepted feasible proposals reset the relaxed input. The mutation budget is relative to each starting sequence. d, Native confidence combines prediction, support, terrain, low risk and gate entropy; segment widths represent their score weights. Brightness and greedy diversity constraints are applied after deduplication. Deviation is evaluated after native candidate identities and rankings are fixed. Full equations, confidence weights and numerical settings are given in Supplementary Methods S1–S4 and Table S1.

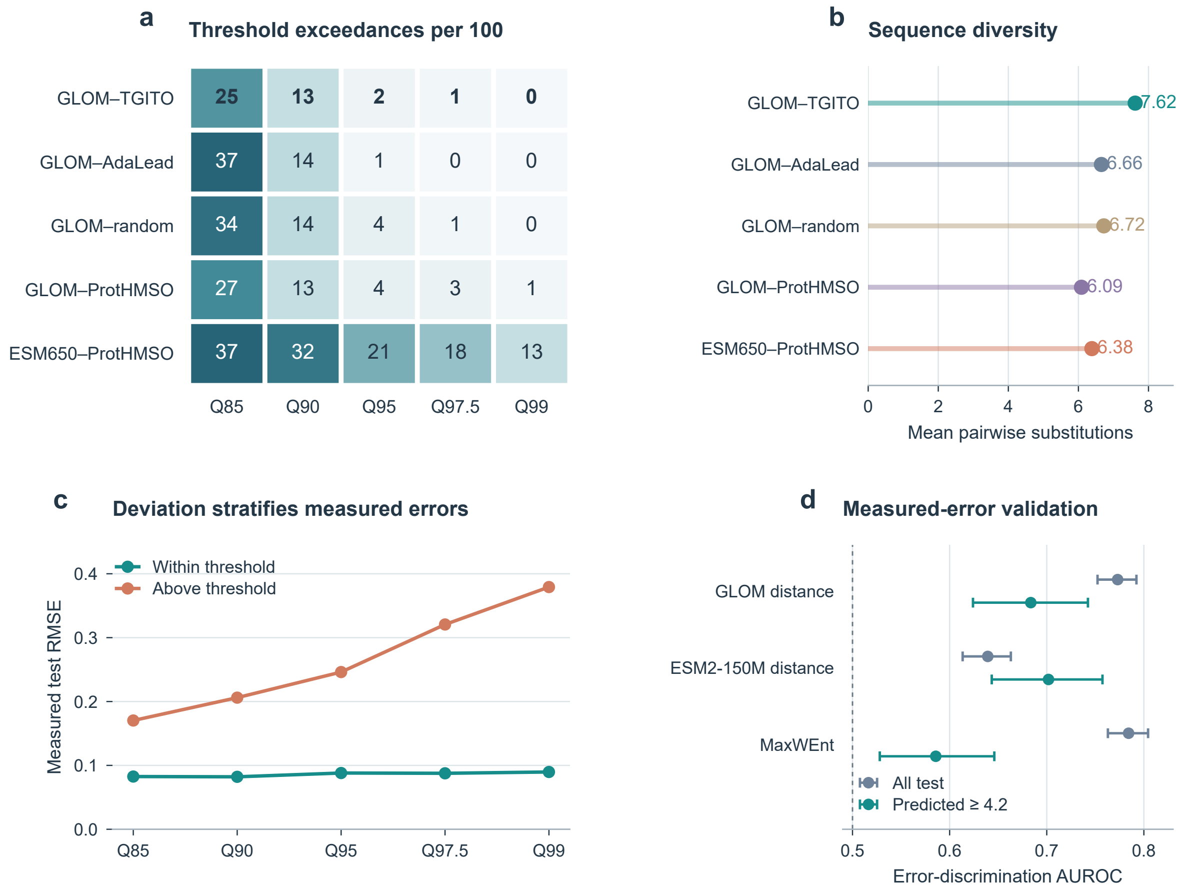


**Figure S8: Threshold comparisons and measured-error validation for discovery assessment**. a, Numbers of native Top100 candidates above Q85, Q90, Q95, Q97.5 and Q99, with thresholds fixed on calibration variants. b, Mean pairwise Hamming distance, used here to describe sequence diversity. c, Prediction RMSE below and above the same deviation thresholds among 973 measured test variants with predicted fluorescence $\geq$4.2. Above-threshold group sizes are 158, 100, 49, 28 and 17, respectively (Table S6). d, AUROC for absolute prediction error $>$0.1 on all 3,144 test variants and on the high-prediction subset. Intervals are 95% bootstrap intervals from 1,000 resamples. GLOM representation distance is compared with raw ESM2-150M distance and a MaxWEnt regression-head pilot. Panels c and d use measured-test errors from the search checkpoint to assess the diagnostic value of deviation; candidate brightness rescoring uses the two other GLOM seeds.

### S12 Complete threshold comparison and detailed architecture

Table S18 reports Q85–Q99 exceedances, sequence novelty and joint acceptance in the native Top 100 batches. Table S9 provides the wider Q50–Q99 grid; Tables S7, S8 and S13 report neighbourhood-size, complete-pool and selected-batch sensitivity. Main Fig. 1 summarizes the architecture; Section S1 specifies layer dimensions and selected sites; Fig. S9 summarizes the separation of training, search and assessment. Fig. S10 shows the complete recorded trajectory with relaxed predictions, hard-sequence predictions and accepted proposals.

**Table S18:** Threshold exceedance and joint novelty–reliability in native Top100 batches

| Method | Q85 | Q90 | Q95 | Q97.5 | Q99 | Novel ≥2 | Joint pass |
|---|---|---|---|---|---|---|---|
| GLOM–TGITO | 25% | 13% | 2% | 1% | 0% | 90 | 38 |
| GLOM–AdaLead | 37% | 14% | 1% | 0% | 0% | 44 | 16 |
| GLOM–Random | 34% | 14% | 4% | 1% | 0% | 42 | 14 |
| GLOM–ProtHMSO | 27% | 13% | 4% | 3% | 1% | 20 | 14 |
| ESM650–ProtHMSO | 37% | 32% | 21% | 18% | 13% | 27 | 11 |

Each batch contains 100 sequences. Q columns count candidates beyond high-prediction calibration distance quantiles. Novel ≥2 counts sequences at least two substitutions from every visible training/validation/calibration sequence. Joint pass additionally requires a risk-scaled empirical lower-brightness reference ≥4.2, combined risk ≤0.15761 and cross-seed SD ≤0.014576. The full rule and threshold grid appear in Supplementary Tables S15–S16. Counts describe the frozen batches; rankings were not changed.

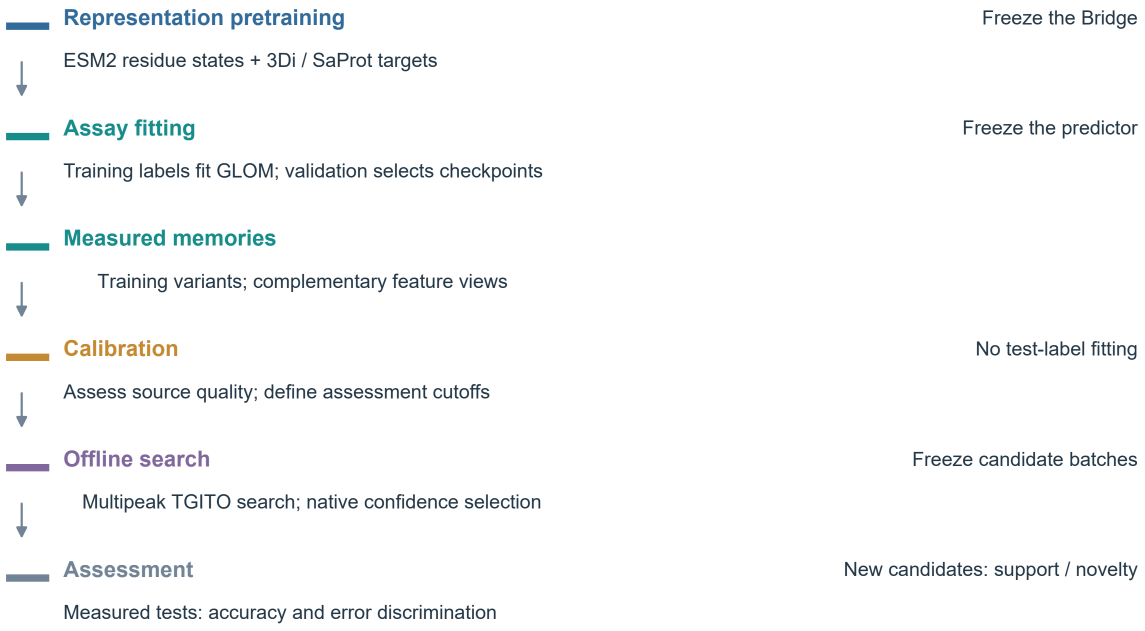


**Figure S9: Data roles across the integrated workflow**. Bridge pretraining supplies a frozen representation. Assay training fits the GLOM head and branches and populates measured memories; validation selects fitted checkpoints. Calibration assesses source quality and fixes assessment thresholds. Search changes sequence inputs while all fitted models and measured memories remain fixed. Native selection precedes post-hoc deviation analysis. Measured tests assess regression, screening and error discrimination. Historical feature-panel and HPO provenance is described in Section S9.

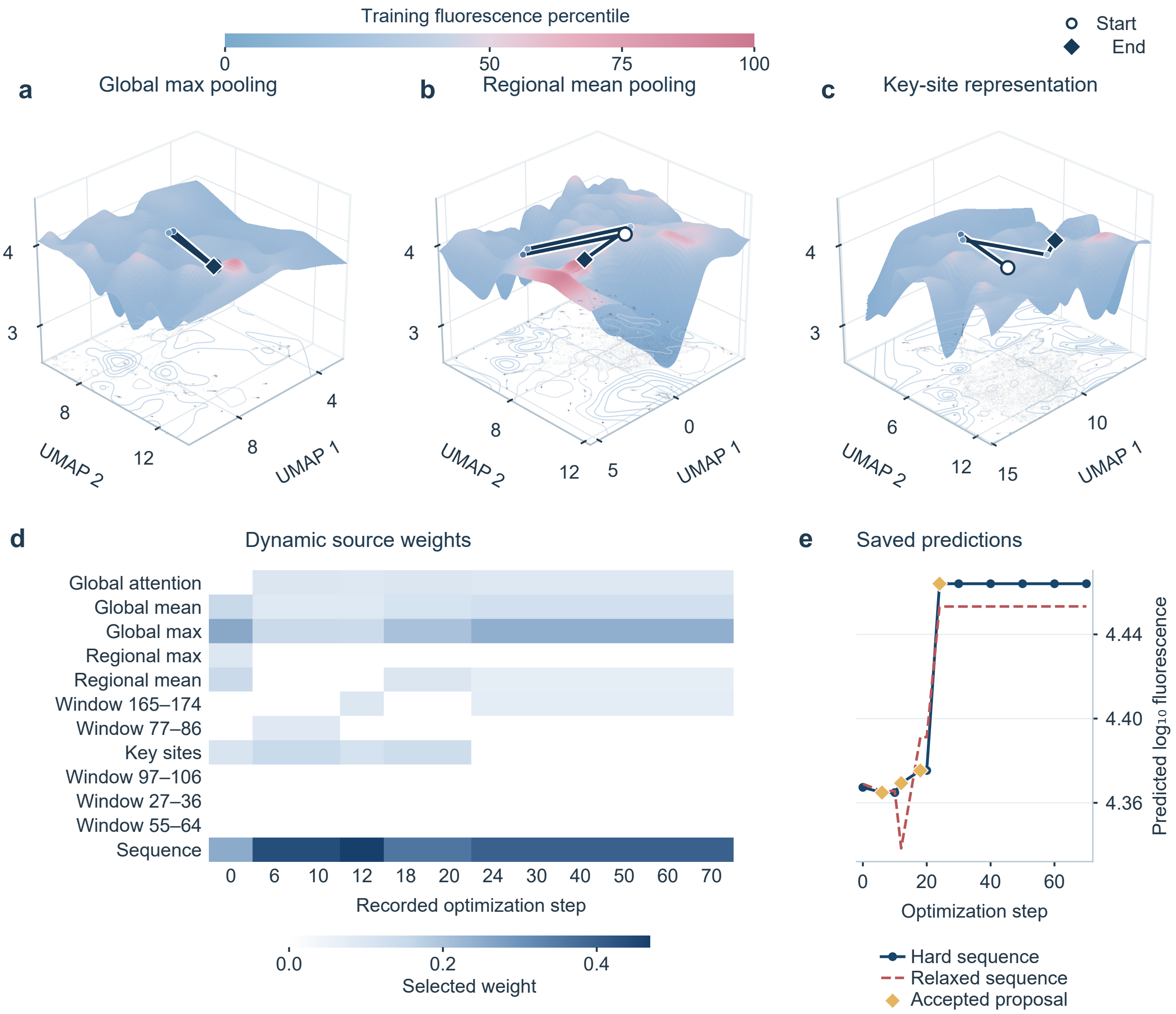


**Figure S10: Complete recorded multiterrain trajectory and hard-sequence acceptance**. a–c, Measured fluorescence landscapes from 22,003 training sequences in global, regional and key-site representations. Blue to red denotes increasing fluorescence percentile relative to the training labels, with red beginning at their median (4.1303); elevation remains linear in $\log_{10}$ fluorescence. The same recorded relaxed-query trajectory is shown in all three views; it was chosen by the highest-ranked native Top6 candidate (context 0054). Open circles mark initialization and diamonds mark step 70. Path heights follow the displayed surfaces. d, Source weights at twelve saved states. Six of twelve sources are active at each state, with four changes in the active set between saved states. Columns represent saved states rather than equal time intervals. e, Hard- and relaxed-sequence predictions, with four accepted discrete proposals. Across the campaign, 900 regions and three roles give 2,700 tasks, with 1,308 distinct starting sequences. Online gradients use the original feature spaces; UMAP surfaces and paths visualize their geometry. Matched measured and predicted surfaces are shown in Supplementary Fig. S3, endpoint source contributions in Supplementary Fig. S6 and computational details in Supplementary Fig. S7.

### S13 SaProt controls trained on the project partitions

#### Model, inputs and training

All four controls use the native 33-layer SaProt-650M encoder with 1,280-dimensional states and its 446-token joint vocabulary [3]. We start from the general pretrained `westlake-repl/SaProt_650M_AF2` weights. No externally fluorescence-fine-tuned task head or adapter is loaded. The two input modes are amino-acid-only (AA), with every structural position masked by `#`, and amino acid plus predicted 3Di (3Di). The latter uses argmax tokens from the same frozen ESM2-150M/Bridge used in this project; its padding state maps to `#`. Each input has 221 residue tokens plus beginning/end tokens.

The frozen-MLP control averages residue states excluding special tokens and fits the same standardized two-layer head as the other frozen encoders. The mean-pool LoRA control updates query/value adapters with rank 16, scale 32 and dropout 0.05, and fits a LayerNorm–256–GELU–dropout–1 head. Both use train-standardized Huber loss and select checkpoints by validation Spearman. The latter uses AdamW at $10^{-4}$, weight decay 0.01 and effective batch size 64, with a 25-epoch cap and patience six.

The two CLS controls adapt the SaProtHub fluorescence recipe [15]. A CLS regression head applies dropout, a $1280 \rightarrow 1280$ linear layer, tanh, dropout and a scalar output. Rank-eight LoRA (scale 16, zero adapter dropout) targets query, key, value, intermediate dense and output dense layers; the classifier is trained as well. Each fit updates 7,723,521 of 660,607,383 parameters. AdamW uses learning rate $10^{-4}$, betas (0.9,0.98), weight decay 0.01 and gradient clipping at 1. A microbatch of sixteen with four accumulation steps gives an effective batch of 64. The learning rate follows a cosine schedule to 5% of its initial value. Raw-label MSE is minimized, and validation MSE selects the checkpoint. The AA and 3Di runs differ only in structural input.

The official model card specifies fifty epochs and mixed FP16. Our adaptation uses a 25-epoch cap, patience six and BF16 with FP32 master weights, matching the local adapter budget. It also uses the project partitions and seeds. Table S19 summarizes the four settings. All runs reserve calibration and test labels from fitting and checkpoint selection, and all three seeds are reported without test-based selection of an input mode.

#### Results and paired comparisons

Tables S2–S3 include all four SaProt configurations alongside the other predictors. Main Fig. 3 compares the complete GLOM predictor with representative baselines; Fig. S11 shows every SaProt control. For the main error distributions, each test variant contributes its mean absolute residual across three fitted seeds, rather than the error of an ensemble prediction. Non-negative reflected kernel densities display these values; medians, quartiles and 95th percentiles are computed directly from the per-variant values. Table S20 gives the individual fits. The lowest SaProt mean RMSE is 0.246532 on random holdout (CLS, 3Di) and 0.323892 on depth holdout (CLS, AA), compared with 0.169986 and 0.246827 for GLOM. The corresponding reductions are 31.0% and 23.8%. SaProt benefits from adaptation relative to its frozen-MLP control, but the relative merits of AA and predicted 3Di differ between splits.

For each split and baseline, we align prediction rows and labels to GLOM, match the three training seeds, and compute GLOM-minus-baseline RMSE and MAE. Each of 2,000 bootstrap replicates resamples test rows with replacement, uses the same rows for both models and all seeds, computes the metric difference per seed, and averages the three differences. The 2.5th and 97.5th percentiles form the intervals in Table S21. Every RMSE and MAE interval lies below zero. These intervals describe test-row uncertainty conditional on the fitted models. Across-seed SD in Tables S2–S3 reports fitting variation.

Table S14 compares measured-test prioritization, and Table S22 reports measured means, medians, minima, bright fractions and top-decile enrichment for Top 6/30/100 selections. Averaged across three training seeds, GLOM's Top 100 measured bright fractions are 96.3% and 97.3%, versus 52.0–64.7% and 42.3–61.7% across the SaProt controls. The best SaProt mean brightness does not always come from the configuration with the lowest global RMSE, which illustrates why screening and regression are both reported. Supplementary Fig. S11 shows RMSE, MAE, $R^2$ and Spearman for every SaProt fit.

#### Reproducibility and scope

The two protocol snapshots pin the general pretrained model, official implementation and Bridge checkpoint by revision or SHA256. They also record data hashes, token mappings, optimizer settings and checkpoint-selection rules. All models were evaluated on identical sequence identities and labels: 3,144 random-test and

14,280 depth-test variants. This comparison comprises 24 SaProt fits and six GLOM fits across the two splits. Prediction files, per-run configurations, histories and source scripts are archived with the experiment.

These controls evaluate the fluorescence-prediction stage of GLOM–TGITO using SaProt representations on the same assay partitions. Discovery candidates and their native ranks were fixed before these supervised comparisons.

**Table S19:** SaProt-650M configurations on the fixed project partitions

| Configuration | Input | Pooling / head | Adapter | Loss / validation criterion | Epoch cap / patience |
|---|---|---|---|---|---|
| Frozen MLP | 3Di | Mean; 512/256 MLP | Encoder frozen | Standardized Huber; Spearman | 200 / 20 |
| Mean-pool LoRA | 3Di | Mean; LN/256/1 | r16; Q,V | Standardized Huber; Spearman | 25 / 6 |
| CLS LoRA (AA) | AA | CLS; 1280/1 | r8; Q,K,V,dense | Raw MSE; MSE | 25 / 6 |
| CLS LoRA (3Di) | 3Di | CLS; 1280/1 | r8; Q,K,V,dense | Raw MSE; MSE | 25 / 6 |

Each configuration has three seeds on each split. AA masks structural tokens; 3Di uses frozen Bridge predictions. Q/K/V denote attention query/key/value projections. The two CLS rows use the official-recipe adaptation; all task heads and adapters are trained on the project training data.

**Table S20:** Individual GLOM and SaProt fits

| Split | Configuration | Seed | RMSE | MAE | $R^2$ | $\rho$ | Pearson |
|---|---|---|---|---|---|---|---|
| Random | GLOM | 20261013 | 0.1703 | 0.0842 | 0.8913 | 0.8771 | 0.9457 |
| Random | GLOM | 20261014 | 0.1691 | 0.0810 | 0.8929 | 0.8922 | 0.9464 |
| Random | GLOM | 20261015 | 0.1706 | 0.0826 | 0.8909 | 0.8872 | 0.9462 |
| Random | frozen MLP | 20261013 | 0.4075 | 0.2619 | 0.3776 | 0.5261 | 0.6269 |
| Random | frozen MLP | 20261014 | 0.4255 | 0.2828 | 0.3212 | 0.4693 | 0.5836 |
| Random | frozen MLP | 20261015 | 0.4157 | 0.2720 | 0.3521 | 0.4901 | 0.6074 |
| Random | mean-pool LoRA | 20261013 | 0.2648 | 0.1379 | 0.7372 | 0.6962 | 0.8677 |
| Random | mean-pool LoRA | 20261014 | 0.2938 | 0.1550 | 0.6765 | 0.6652 | 0.8461 |
| Random | mean-pool LoRA | 20261015 | 0.2575 | 0.1340 | 0.7514 | 0.6904 | 0.8762 |
| Random | CLS LoRA (AA) | 20261013 | 0.2727 | 0.1630 | 0.7213 | 0.7831 | 0.8944 |
| Random | CLS LoRA (AA) | 20261014 | 0.2344 | 0.1271 | 0.7941 | 0.7520 | 0.9068 |
| Random | CLS LoRA (AA) | 20261015 | 0.2629 | 0.1559 | 0.7409 | 0.7449 | 0.8982 |
| Random | CLS LoRA (3Di) | 20261013 | 0.2515 | 0.1511 | 0.7628 | 0.7477 | 0.9067 |
| Random | CLS LoRA (3Di) | 20261014 | 0.2414 | 0.1399 | 0.7815 | 0.7287 | 0.9025 |
| Random | CLS LoRA (3Di) | 20261015 | 0.2466 | 0.1526 | 0.7720 | 0.7447 | 0.9120 |

GLOM entries are the six frozen reference fits. All SaProt rows use the 650M base. Errors are on the original log-fluorescence scale; no post-hoc test calibration is applied.

**Table S20:** Individual GLOM and SaProt fits (continued)

| Split | Configuration | Seed | RMSE | MAE | $R^2$ | $\rho$ | Pearson |
|---|---|---|---|---|---|---|---|
| Depth | GLOM | 20261013 | 0.2475 | 0.1332 | 0.8127 | 0.8695 | 0.9114 |
| Depth | GLOM | 20261014 | 0.2442 | 0.1290 | 0.8176 | 0.8744 | 0.9140 |
| Depth | GLOM | 20261015 | 0.2487 | 0.1315 | 0.8109 | 0.8661 | 0.9106 |
| Depth | frozen MLP | 20261013 | 0.5581 | 0.3750 | 0.0477 | 0.4378 | 0.4919 |
| Depth | frozen MLP | 20261014 | 0.5526 | 0.3710 | 0.0664 | 0.4411 | 0.4970 |
| Depth | frozen MLP | 20261015 | 0.5583 | 0.3775 | 0.0468 | 0.4210 | 0.4733 |
| Depth | mean-pool LoRA | 20261013 | 0.4421 | 0.2602 | 0.4025 | 0.6360 | 0.7193 |
| Depth | mean-pool LoRA | 20261014 | 0.4126 | 0.2414 | 0.4795 | 0.6566 | 0.7486 |
| Depth | mean-pool LoRA | 20261015 | 0.4257 | 0.2478 | 0.4460 | 0.6447 | 0.7363 |
| Depth | CLS LoRA (AA) | 20261013 | 0.3678 | 0.2196 | 0.5865 | 0.7692 | 0.8302 |
| Depth | CLS LoRA (AA) | 20261014 | 0.2663 | 0.1503 | 0.7831 | 0.8227 | 0.8906 |
| Depth | CLS LoRA (AA) | 20261015 | 0.3376 | 0.1957 | 0.6516 | 0.7825 | 0.8464 |
| Depth | CLS LoRA (3Di) | 20261013 | 0.3984 | 0.2515 | 0.5147 | 0.6995 | 0.8077 |
| Depth | CLS LoRA (3Di) | 20261014 | 0.4083 | 0.2496 | 0.4902 | 0.6549 | 0.7535 |
| Depth | CLS LoRA (3Di) | 20261015 | 0.3916 | 0.2470 | 0.5312 | 0.7059 | 0.8107 |

GLOM entries are the six frozen reference fits. All SaProt rows use the 650M base. Errors are on the original log-fluorescence scale; no post-hoc test calibration is applied.

**Table S21:** Paired prediction-error differences: GLOM minus SaProt

| Split | SaProt configuration | Metric | Difference | 95% interval |
|---|---|---|---|---|
| Random | frozen MLP | RMSE | -0.2462 | [-0.2632, -0.2300] |
| Random | frozen MLP | MAE | -0.1896 | [-0.1997, -0.1797] |
| Random | mean-pool LoRA | RMSE | -0.1020 | [-0.1148, -0.0881] |
| Random | mean-pool LoRA | MAE | -0.0597 | [-0.0654, -0.0541] |
| Random | CLS LoRA (AA) | RMSE | -0.0867 | [-0.0980, -0.0761] |
| Random | CLS LoRA (AA) | MAE | -0.0661 | [-0.0710, -0.0613] |
| Random | CLS LoRA (3Di) | RMSE | -0.0765 | [-0.0874, -0.0654] |
| Random | CLS LoRA (3Di) | MAE | -0.0653 | [-0.0698, -0.0609] |
| Depth | frozen MLP | RMSE | -0.3095 | [-0.3177, -0.3008] |
| Depth | frozen MLP | MAE | -0.2433 | [-0.2496, -0.2367] |
| Depth | mean-pool LoRA | RMSE | -0.1800 | [-0.1870, -0.1728] |
| Depth | mean-pool LoRA | MAE | -0.1186 | [-0.1229, -0.1142] |
| Depth | CLS LoRA (AA) | RMSE | -0.0771 | [-0.0820, -0.0721] |
| Depth | CLS LoRA (AA) | MAE | -0.0573 | [-0.0600, -0.0549] |
| Depth | CLS LoRA (3Di) | RMSE | -0.1526 | [-0.1588, -0.1461] |
| Depth | CLS LoRA (3Di) | MAE | -0.1182 | [-0.1220, -0.1144] |

Negative differences favour GLOM. Intervals use 2,000 paired test-row bootstrap resamples and average the three fixed-seed metric differences. They condition on the fitted models and are not intervals across independent training campaigns.

**Table S22:** Measured-protein prioritization for GLOM and all SaProt configurations

| Split | Configuration | k | Mean | Median | Minimum | $\geq 4.2$ (%) | $\geq 4.35$ (%) | Enrichment |
|---|---|---|---|---|---|---|---|---|
| Random | GLOM | 6 | 4.379±0.011 | 4.383±0.006 | 4.332±0.033 | 100.0±0.0 | 88.9±9.6 | 9.981±0.000 |
| Random | GLOM | 30 | 4.340±0.002 | 4.344±0.001 | 4.203±0.022 | 97.8±1.9 | 44.4±1.9 | 9.426±0.192 |
| Random | GLOM | 100 | 4.324±0.002 | 4.331±0.002 | 4.144±0.042 | 96.3±1.2 | 34.7±3.2 | 8.617±0.115 |
| Random | frozen MLP | 6 | 4.225±0.065 | 4.266±0.019 | 3.983±0.279 | 61.1±9.6 | 22.2±9.6 | 4.990±0.000 |
| Random | frozen MLP | 30 | 4.199±0.022 | 4.236±0.019 | 3.186±0.688 | 64.4±1.9 | 8.9±3.8 | 4.325±0.880 |
| Random | frozen MLP | 100 | 4.179±0.038 | 4.214±0.010 | 2.775±0.000 | 58.3±6.5 | 7.7±0.6 | 3.394±0.780 |
| Random | mean-pool LoRA | 6 | 4.285±0.025 | 4.302±0.022 | 4.162±0.039 | 77.8±9.6 | 16.7±16.7 | 7.208±0.960 |
| Random | mean-pool LoRA | 30 | 4.237±0.046 | 4.242±0.014 | 3.677±0.661 | 74.4±12.6 | 13.3±5.8 | 4.436±0.384 |
| Random | mean-pool LoRA | 100 | 4.227±0.014 | 4.225±0.010 | 3.644±0.629 | 64.7±7.6 | 10.0±1.7 | 3.294±0.173 |
| Random | CLS LoRA (AA) | 6 | 4.210±0.005 | 4.210±0.009 | 4.137±0.021 | 55.6±9.6 | 0.0±0.0 | 2.218±0.960 |
| Random | CLS LoRA (AA) | 30 | 4.202±0.007 | 4.204±0.008 | 4.043±0.042 | 51.1±3.8 | 4.4±1.9 | 1.442±0.192 |
| Random | CLS LoRA (AA) | 100 | 4.204±0.013 | 4.204±0.011 | 3.964±0.126 | 52.0±6.6 | 3.7±1.5 | 1.630±0.472 |
| Random | CLS LoRA (3Di) | 6 | 4.290±0.033 | 4.293±0.048 | 4.182±0.049 | 83.3±16.7 | 22.2±9.6 | 5.545±2.541 |
| Random | CLS LoRA (3Di) | 30 | 4.276±0.030 | 4.267±0.028 | 4.110±0.074 | 80.0±15.3 | 25.6±5.1 | 5.434±1.168 |
| Random | CLS LoRA (3Di) | 100 | 4.223±0.025 | 4.230±0.003 | 3.649±0.661 | 64.7±1.5 | 11.3±1.2 | 3.394±0.346 |

Mean ± sample SD across three fits. Each row first summarizes the measured labels of a prediction-ranked batch, then averages that statistic across seeds. Minimum denotes the mean of the three within-batch minima. Enrichment is relative to the measured top-decile prevalence in the corresponding test set. All batches rank measured test sequences, with no terrain/confidence filter.

**Table S22:** Measured-protein prioritization for GLOM and all SaProt configurations (continued)

| Split | Configuration | k | Mean | Median | Minimum | $\geq 4.2$ (%) | $\geq 4.35$ (%) | Enrichment |
|---|---|---|---|---|---|---|---|---|
| Depth | GLOM | 6 | 4.348±0.020 | 4.354±0.019 | 4.307±0.037 | 100.0±0.0 | 55.6±19.2 | 10.000±0.000 |
| Depth | GLOM | 30 | 4.334±0.018 | 4.337±0.025 | 4.220±0.028 | 98.9±1.9 | 43.3±11.5 | 9.333±0.667 |
| Depth | GLOM | 100 | 4.331±0.005 | 4.341±0.004 | 4.076±0.159 | 97.3±2.1 | 42.3±1.5 | 9.267±0.153 |
| Depth | frozen MLP | 6 | 3.951±0.077 | 4.235±0.059 | 2.783±0.021 | 55.6±9.6 | 16.7±16.7 | 5.000±1.667 |
| Depth | frozen MLP | 30 | 4.055±0.114 | 4.234±0.045 | 2.771±0.000 | 57.8±8.4 | 18.9±1.9 | 5.000±0.882 |
| Depth | frozen MLP | 100 | 4.044±0.034 | 4.194±0.033 | 2.771±0.000 | 48.0±6.1 | 11.7±2.1 | 3.867±0.709 |
| Depth | mean-pool LoRA | 6 | 4.245±0.047 | 4.291±0.016 | 3.930±0.194 | 77.8±9.6 | 27.8±25.5 | 6.111±0.962 |
| Depth | mean-pool LoRA | 30 | 4.232±0.017 | 4.254±0.015 | 3.743±0.130 | 71.1±5.1 | 16.7±0.0 | 5.000±0.333 |
| Depth | mean-pool LoRA | 100 | 4.171±0.006 | 4.227±0.016 | 2.825±0.036 | 61.7±7.6 | 12.7±2.1 | 4.400±0.361 |
| Depth | CLS LoRA (AA) | 6 | 4.218±0.071 | 4.227±0.076 | 4.099±0.053 | 55.6±25.5 | 16.7±28.9 | 3.889±3.469 |
| Depth | CLS LoRA (AA) | 30 | 4.187±0.017 | 4.186±0.024 | 3.953±0.169 | 44.4±13.5 | 3.3±5.8 | 2.444±0.770 |
| Depth | CLS LoRA (AA) | 100 | 4.180±0.019 | 4.186±0.022 | 3.578±0.359 | 42.3±8.5 | 4.7±0.6 | 2.333±0.551 |
| Depth | CLS LoRA (3Di) | 6 | 4.218±0.092 | 4.207±0.082 | 4.089±0.133 | 61.1±41.9 | 16.7±16.7 | 4.444±3.469 |
| Depth | CLS LoRA (3Di) | 30 | 4.179±0.046 | 4.197±0.037 | 3.421±0.560 | 53.3±11.5 | 14.4±7.7 | 3.778±0.962 |
| Depth | CLS LoRA (3Di) | 100 | 4.181±0.015 | 4.204±0.006 | 2.936±0.183 | 52.3±3.1 | 11.3±2.5 | 3.633±0.404 |

Mean ± sample SD across three fits. Each row first summarizes the measured labels of a prediction-ranked batch, then averages that statistic across seeds. Minimum denotes the mean of the three within-batch minima. Enrichment is relative to the measured top-decile prevalence in the corresponding test set. All batches rank measured test sequences, with no terrain/confidence filter.

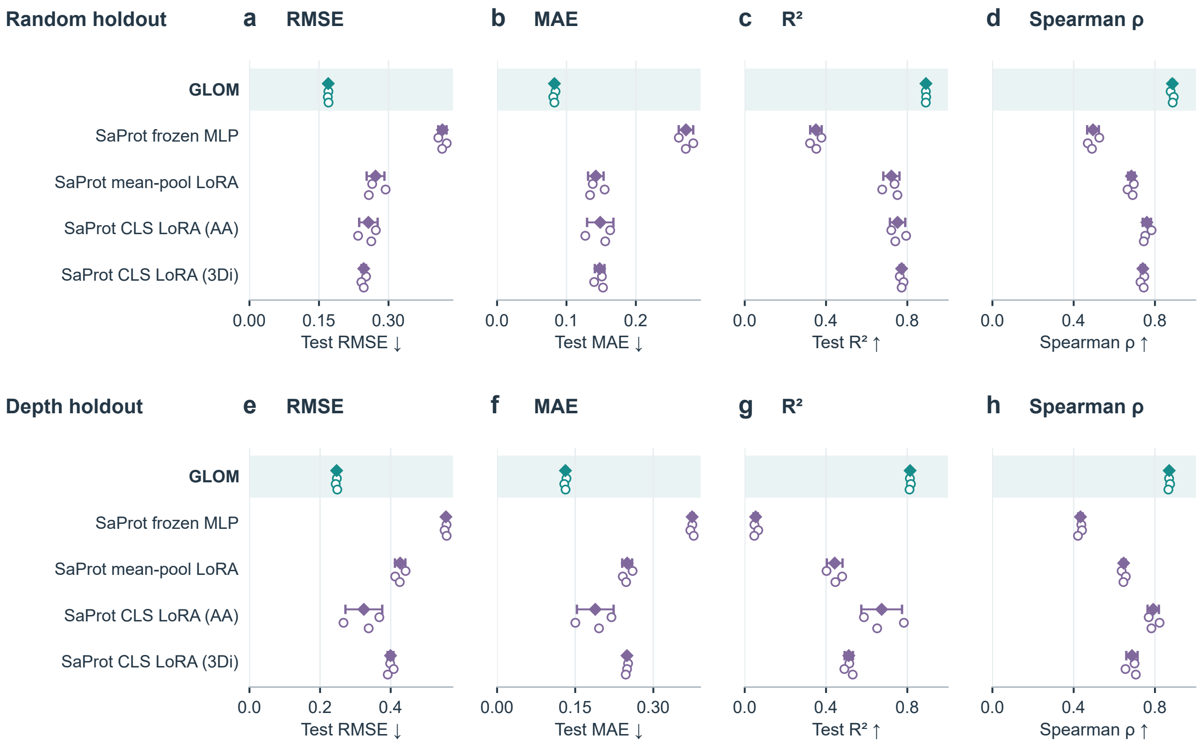


**Figure S11: SaProt prediction controls across both holdouts**. Rows show random and mutation-depth tests; columns show RMSE, MAE, $R^2$ and Spearman correlation. The same five configurations appear in each panel. Open circles indicate the three fitted seeds, with diamonds and intervals for their mean and sample SD. All SaProt controls start from the general pretrained 650M model and use the project partitions. CLS controls adapt the official fluorescence recipe. The four metrics report different aspects of prediction; the full numerical results are in Tables S2–S3 and S20.

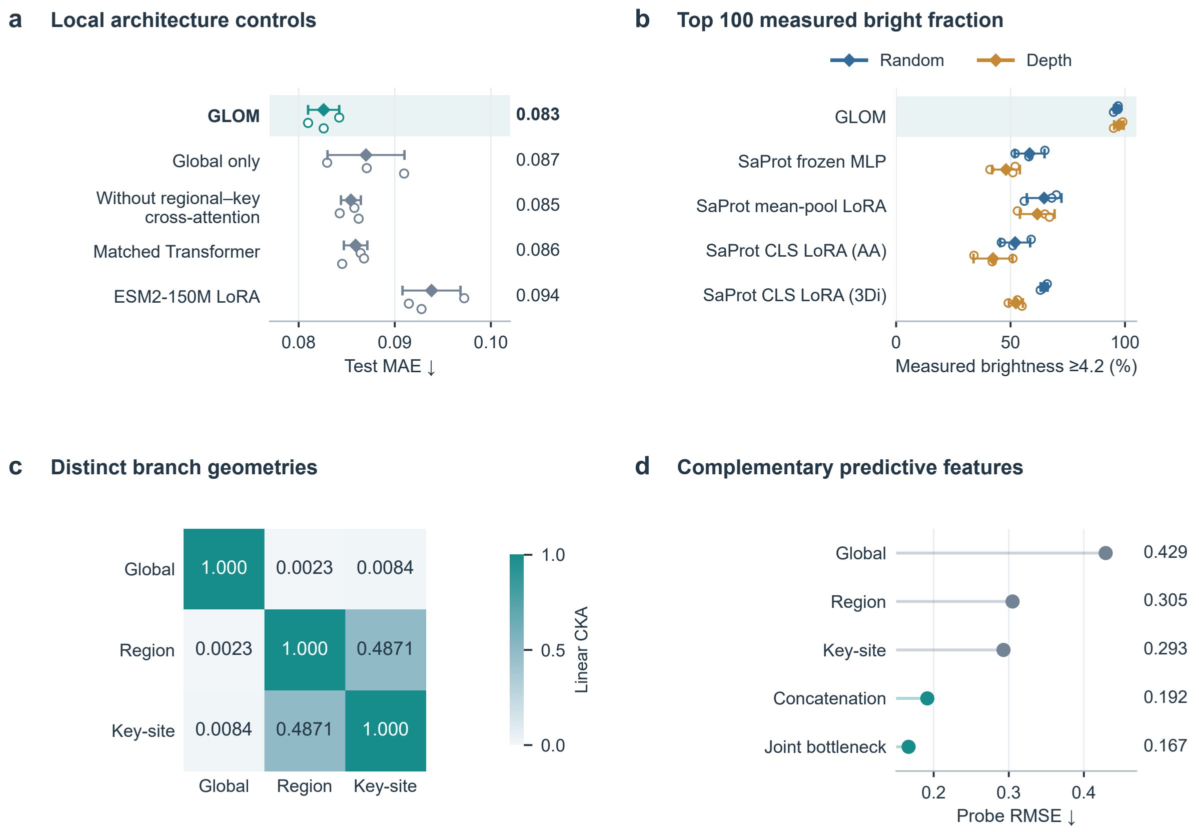


**Figure S12: Architectural controls, measured prioritization and complementary representations. A.** Random-holdout MAE for retrained architecture controls and ESM2-150M LoRA. **B.** Measured brightness $\geq 4.2$ among prediction-ranked Top 100 test variants. Open circles show training seeds; diamonds and bars show means and sample SDs. **C.** Linear CKA on 1,000 test sequences. **D.** Matched ridge-probe RMSE on 3,144 test sequences. C,D use GLOM seed 20261013. Low CKA denotes different representation geometries. SaProt controls use the 650M base.

### S14. Biophysical analysis of the native Top 30

#### S14.1 Candidate identities, structure preparation and simulations

The analysis includes all thirty sequences selected by the formal campaign's native confidence/diversity procedure, their twenty-five distinct starts and WT. Selection was frozen before the molecular analysis. Each candidate is compared with its own start, so substitutions introduced during search are distinguished from those inherited from the starting sequence. The six illustrated examples were chosen afterwards for their surface, contact or torsional patterns and retain their original ranking IDs (Table S12).

All systems were built from PDB 2G3O chain C, retaining the experimental heavy-atom geometry of the mature chromophore [16]. Project position $p$ corresponds to author position $p+1$ outside the chromophore; G56–Y57–G58 form the CR2 residue. PDBFixer supplied missing side chains and the unresolved terminal residues 219–221. The same repair and relaxation procedure was applied to WT, starts and candidates. The simulations describe the mature anionic chromophore, with a formal charge of $-1$. Asp/Glu and Lys/Arg used their standard charged states, and histidine used the neutral HIE form. Protonation states were fixed.

OpenMM simulations used ff14SB with compatible chromophore parameters from Breyfogle et al. [17], OPC water, a truncated-octahedral solvent box with 10 Å padding and 0.15 M NaCl plus neutralizing ions [18]. A Langevin-middle integrator maintained 300 K with a friction coefficient of 1 $ps^{-1}$ and a 2-fs timestep; bonds involving hydrogen were constrained. A Monte Carlo barostat maintained 1 bar, and particle-mesh Ewald electrostatics used a 1-nm real-space cutoff. After minimization, each system underwent 50 ps of strong and 50 ps of weaker heavy-atom positional restraints, followed by 400 ps of unrestrained equilibration. Production trajectories were unrestrained for 3 ns, with coordinates saved every 10 ps. Three runs with different initial-velocity seeds were completed for all 56 systems (168 production trajectories). A common seed was used across systems within each run. The first run supplies the primary comparison, and runs 2 and 3 assess repeatability for all 30 candidate/start/WT triplets (Table S25 and Supplementary Data 4).

#### S14.2 Molecular descriptors and paired comparisons

Trajectories were aligned to WT C-alpha positions 3–215. Hydrogen bonds require donor–acceptor distance $\leq 3.5$ Å and donor–hydrogen–acceptor angle $\geq 150°$. Phenolate occupancy is the fraction of frames with at least one protein hydrogen bond to the chromophore phenolate oxygen. Thr137 and Arg155 occupancies are also recorded individually and jointly. Chromophore torsions use N2–CA2–CB2–CG2 (I) and CA2–CB2–CG2–CD1 (P); circular SD avoids the discontinuity at $\pm 180°$. Heavy-atom mean-square fluctuations describe motion around the aligned trajectory-average coordinates.

Solvent-accessible surface areas use a 1.4 Å probe and 960 sampling points per heavy atom, evaluated every 50 ps. Hydrophobic surface atoms are the side-chain carbon/sulfur atoms of A, V, I, L, M, F, W, Y, P and C. For each exposed hydrophobic atom, the patch score sums the accessible area of hydrophobic atoms within a 5 Å radius; the largest score in each frame is then averaged over the trajectory. The principal score excludes residues buried by at least 5 Å$^2$ at the reference C/E dimer interface. This separates exposed surface patches from the native oligomer interface. It includes the modelled terminal segment; a second calculation is restricted to resolved positions 1–218. Four- and six-angstrom radii test sensitivity to the patch definition (Fig. S14).

For every metric $m$, start-relative and WT-relative differences are $m_{\text{candidate}} - m_{\text{start}}$ and $m_{\text{candidate}} - m_{\text{WT}}$. Relative changes divide by the corresponding reference value. The 5.2% cohort patch decrease in the main text is the relative difference between the two cohort means; it is not the mean of individual percentage changes. In the start-balanced summary, candidate values are first averaged within each of the 25 start groups, and those groups receive equal weight. Trajectory halves and independent initial-velocity seeds test the persistence of an effect. Saved frames are time-correlated observations, not independent replicates; candidate counts and changes are descriptive. These ground-state simulations assess surface and chromophore environments and do not model chromophore formation or excited-state emission.

#### S14.3 Cohort changes and representative molecular routes

In the first run, twenty of thirty candidates have smaller surface patches than their actual starts, and eighteen have smaller patches than WT (Table S23). The mean is 178.1 Å$^2$ for candidates, 187.8 Å$^2$ for their paired starts and 181.4 Å$^2$ for WT. Radius-4 and radius-6 calculations also give lower cohort mean patch areas, by 3.4% and 3.6%. Fourteen candidates retain the direction in both trajectory halves, and eleven retain it at all three radii. Excluding modelled residues 219–221 gives almost equal cohort means (141.15 versus 141.06 Å$^2$), indicating that terminal exposure contributes to the aggregate difference. C14 and C27 still show smaller patches in this restricted calculation, by 13.0% and 10.1%, respectively. Nine candidates combine a smaller primary patch with increased phenolate hydrogen bonding; eleven combine a smaller patch with reduced I-axis fluctuations. Twenty-five have lower I-axis SD than WT, and sixteen have lower I-axis SD than their starts. The remaining readouts show distinct responses across candidates (Fig. S14; Supplementary Data 3).

**Surface chemistry: C12 and C14.** C12 introduces E122K/A219D into an E105D/L199H background. Its largest non-interface patch decreases from 199.1 to 164.9, from 221.6 to 175.7 and from 188.9 to 181.4 Å$^2$ in runs 1–3, respectively. P-axis fluctuations decrease and phenolate occupancy increases in runs 1 and 2, but both directions reverse in run 3 (Main Fig. 5f). C14 introduces K36N/D129G into a Q30R/G81S/G206E background. Its patch decreases in all three runs, by 19.1%, 8.9% and 10.7%, including the first-run calculation restricted to resolved residues. Phenolate occupancy increases in runs 1 and 3 and decreases in run 2. The surface change is therefore the more consistent feature of C14. Reducing exposed hydrophobicity can affect aggregation and fluorescent-protein yield, making these surface changes relevant to reporter development. Cycle3 GFP and TurboGFP provide experimental precedents for that surface route [19, 20].

**Phenolate contacts: C21, C22 and C27.** C21 adds M3I/G206E, C22 combines V168L with a reversion from V126 to the WT I126, and C27 adds F167V/V218E to its actual start. Their first-run phenolate occupancies rise by 42.3, 11.7 and 36.7 percentage points, respectively. C27 increases simultaneous Thr137/Arg155 contacts from 9.3% to 50.0%, with 24.7% in WT. The equivalent Thr138/Arg156 pair contacts the phenolate in copGFP crystal structures [21]; these are positions 137/155 in the project sequence. The six maps in Fig. S13 locate both inherited and newly introduced changes. C27 also replaces a hydrophobic terminal residue with glutamate: I218E relative to WT, or V218E relative to its start. This changes terminal charge in the same region targeted during TurboGFP surface engineering [20]. The coupled surface and pocket effects make C27 an example of a design acting in more than one structural context.

More persistent phenolate contacts can change the environment of the emitting chromophore. In YuzuFP, MD-guided modification of the phenolate hydrogen-bond network accompanied increased brightness [22]. The YuzuFP results provide a precedent for engineering the local phenolate interaction network. C21 and C22 show increased phenolate occupancy with different responses in the two torsional coordinates, distinguishing changes in the contact network from changes in chromophore motion.

**Chromophore motion: C30.** C30 introduces E122A/G206E into a K36E/L149M background. Its I/P circular SDs are 7.90/6.53°, compared with 8.25/7.96° in its start and 8.93/7.38° in WT. Both coordinates decrease in each trajectory half. Chromophore heavy-atom mean-square fluctuation decreases from 0.217 to 0.153 Å$^2$ relative to the start (29.6%), versus 0.177 Å$^2$ for WT. These changes locate a plausible route through restricted chromophore motion. Experimental work links competition between fluorescence and non-radiative isomerization to the steric and electrostatic environment of the I and P bonds [23].

**Additional ionic contacts.** Search can also change contacts across barrel regions. C01 introduces G23K/R32S and forms a K23–E209 contact in every saved frame of runs 1 and 2, with mean closest charged-atom distances of 2.683 and 2.682 Å. C09 introduces A106E/G206E and shows K36–E206 contact occupancies of 53.3% and 51.3% in runs 1 and 2; C24 also forms this contact in its first run. These contacts are defined by charged N–O distance $\leq 4$ Å. The complete contact analysis accompanies the supplementary data. Ionic networks have been implicated in improved superfolder-GFP scaffold properties [24]. The new contacts identify another way in which the designed substitutions reorganize interactions across the barrel.

#### S14.4 Existing sequence evidence and reporter applications

We also searched the 22,003 training sequences for pairs identical outside each substitution present in the Top 30 or introduced from its start. Fifty-five of 79 distinct replacements have at least one exact-background contrast. L199H has positive fluorescence differences in 53/54 backgrounds (mean +0.140 log units), and G206E in 27/29 (+0.181). L199H occurs in nineteen candidates and is inherited in all nineteen; G206E occurs in ten candidates and is introduced during search in six. K36N, introduced in C14, has ten positive contrasts out of ten (+0.119). These matched training-library contrasts help interpret the effects of individual substitutions across sequence backgrounds. Supplementary Data 3 reports all contrasts and identifies substitutions lacking matched pairs.

The biophysical results suggest different priorities for reporter development. The surface route is relevant to soluble protein abundance and aggregation-sensitive fusion tags, whereas phenolate contacts and torsional restriction concern the mature chromophore environment. C12 emphasizes a surface change retained across three runs, C27 couples terminal chemistry with phenolate contacts, and C30 emphasizes restricted chromophore motion. The selected sequences thus offer different molecular starting points for reporter engineering.

Solubility-Weighted Index scores average the published residue weights, while SPURS predicts stability changes from standard-residue backbone inputs [25, 26]. The SPURS multi-mutant checkpoint was evaluated with three decoder-order seeds; these assess inference variability rather than separate model fits. These auxiliary predictions are supplied in Supplementary Data 3 and kept separate from the mature-chromophore simulations and fluorescence predictions.

**Table S23:** Complete Top 30 biophysical changes

| ID | Actual start | Patch vs start (%) | Patch vs WT (%) | H-bond vs start (pp) | I SD vs start (%) | P SD vs start (%) |
|---|---|---|---|---|---|---|
| C01 | S0054 | +3.2 | +6.0 | -11.0 | +2.4 | +0.3 |
| C02 | S2175 | -16.9 | +3.7 | -36.0 | -0.4 | +1.1 |
| C03 | S0441 | -5.6 | -0.3 | +13.0 | +3.8 | +21.6 |
| C04 | S1398 | -7.2 | -7.3 | -54.0 | -6.1 | +5.4 |
| C05 | S0069 | -12.8 | -4.3 | -42.7 | +2.2 | -12.8 |
| C06 | S0148 | +3.2 | +5.8 | -16.7 | -0.4 | +3.0 |
| C07 | S0367 | +2.8 | +1.1 | -5.7 | -5.4 | +15.6 |
| C08 | S0054 | +12.9 | +15.9 | -2.7 | -2.0 | +6.6 |
| C09 | S0938 | +6.9 | +8.6 | -5.0 | -1.3 | +6.1 |
| C10 | S1362 | -9.6 | +13.4 | -12.0 | +4.3 | +10.3 |
| C11 | S0069 | +1.8 | +11.8 | +1.7 | +10.5 | -4.4 |
| C12 | S0069 | -17.2 | -9.1 | +5.3 | +13.7 | -8.8 |
| C13 | S0297 | -1.2 | -19.0 | +21.3 | +11.2 | +4.7 |
| C14 | S0960 | -19.1 | -12.4 | +24.0 | -6.9 | +6.4 |
| C15 | S0069 | -24.7 | -17.3 | +1.3 | +4.7 | +4.0 |

One 3-ns production trajectory per system. Patch refers to the largest 5-angstrom non-interface hydrophobic patch. Hydrogen-bond changes are percentage points; other changes are percentages relative to the named reference. Negative patch and torsion changes indicate smaller areas or fluctuations. Absolute values, all auxiliary descriptors and both reference comparisons are in Supplementary Data 3.

**Table S23:** Complete Top 30 biophysical changes (continued)

| ID | Actual start | Patch vs start (%) | Patch vs WT (%) | H-bond vs start (pp) | I SD vs start (%) | P SD vs start (%) |
|---|---|---|---|---|---|---|
| C16 | S0607 | +16.5 | -7.8 | +4.3 | +0.0 | -0.4 |
| C17 | S1143 | -18.2 | -11.9 | -1.3 | -3.0 | +3.4 |
| C18 | S1593 | +27.2 | +34.1 | -29.0 | +6.9 | +8.5 |
| C19 | S1266 | -4.8 | +5.5 | -64.0 | +9.7 | +21.6 |
| C20 | S1058 | -12.2 | +8.4 | -4.3 | -6.8 | +6.0 |
| C21 | S1337 | -13.7 | -15.9 | +42.3 | +20.1 | +12.1 |
| C22 | S0662 | -8.8 | -8.0 | +11.7 | -4.0 | +5.5 |
| C23 | S0477 | -17.7 | -25.0 | -36.7 | -3.5 | +5.1 |
| C24 | S2575 | +14.4 | -1.5 | -33.0 | -2.3 | -6.1 |
| C25 | S2454 | -5.1 | +10.2 | -28.7 | -11.5 | +0.5 |
| C26 | S0441 | -10.8 | -5.8 | -0.7 | +15.4 | +8.8 |
| C27 | S0810 | -18.8 | -15.4 | +36.7 | -2.1 | -1.2 |
| C28 | S1224 | -9.9 | -0.8 | -19.7 | -0.8 | -8.8 |
| C29 | S2244 | +19.5 | -0.7 | +0.3 | +2.1 | -8.2 |
| C30 | S0238 | -8.9 | -17.2 | +7.0 | -4.2 | -17.9 |

One 3-ns production trajectory per system. Patch refers to the largest 5-angstrom non-interface hydrophobic patch. Hydrogen-bond changes are percentage points; other changes are percentages relative to the named reference. Negative patch and torsion changes indicate smaller areas or fluctuations. Absolute values, all auxiliary descriptors and both reference comparisons are in Supplementary Data 3.

**Table S24:** Absolute biophysical values for the six representative designs

| ID | Patch area (Å$^2$) | Phenolate H-bonds (%) | I circular SD (°) | P circular SD (°) |
|---|---|---|---|---|
| C12 | 199.1 / 164.9 | 87.7 / 93.0 | 8.05 / 9.15 | 7.75 / 7.07 |
| C14 | 196.4 / 158.9 | 67.3 / 91.3 | 8.70 / 8.10 | 6.73 / 7.17 |
| C21 | 176.9 / 152.6 | 46.3 / 88.7 | 7.44 / 8.94 | 6.81 / 7.63 |
| C22 | 183.0 / 167.0 | 78.0 / 89.7 | 8.60 / 8.26 | 6.92 / 7.30 |
| C27 | 189.1 / 153.4 | 57.3 / 94.0 | 9.05 / 8.86 | 7.35 / 7.26 |
| C30 | 164.9 / 150.2 | 71.7 / 78.7 | 8.25 / 7.90 | 7.96 / 6.53 |
| WT | 181.4 | 77.3 | 8.93 | 7.38 |

Each paired cell reports actual start / candidate, using the first production trajectory. WT is a common reference. Complete sequence identities and actual search changes are in Table S12.

**Table S25:** Three-run comparisons for all candidate/start/WT triplets

| ID | Patch Δ (Å$^2$) | H-bond Δ (pp) | P SD Δ (degrees) |
|---|---|---|---|
| C01 | +6.02 / +20.68 / +40.47 | -11.00 / +8.67 / -2.67 | +0.02 / +0.35 / -0.11 |
| C02 | -38.38 / -10.79 / -41.23 | -36.00 / +11.00 / +9.67 | +0.08 / -0.38 / +0.05 |
| C03 | -10.65 / +19.56 / -19.87 | +13.00 / +13.67 / +18.67 | +1.50 / -0.20 / -0.91 |
| C04 | -13.09 / -5.76 / -38.85 | -54.00 / +40.67 / +31.33 | +0.38 / -0.40 / -1.59 |
| C05 | -25.57 / -35.14 / +6.41 | -42.67 / +26.67 / -25.67 | -1.00 / -0.18 / +0.52 |
| C06 | +6.04 / -19.48 / +15.41 | -16.67 / -27.33 / -36.00 | +0.22 / -0.62 / +0.88 |
| C07 | +5.04 / +53.51 / +20.07 | -5.67 / -17.67 / +55.67 | +1.09 / +0.43 / +1.17 |
| C08 | +24.05 / +45.01 / +52.92 | -2.67 / +9.33 / -10.33 | +0.50 / +1.25 / -0.17 |
| C09 | +12.73 / -11.75 / -12.03 | -5.00 / -9.33 / +10.33 | +0.44 / -0.20 / +0.38 |
| C10 | -21.85 / -14.83 / +5.23 | -12.00 / -16.00 / -7.67 | +0.74 / -0.41 / -0.71 |
| C11 | +3.62 / -15.56 / -0.04 | +1.67 / +22.00 / +5.00 | -0.34 / -0.52 / -0.28 |
| C12 | -34.29 / -45.85 / -7.48 | +5.33 / +16.00 / -29.33 | -0.68 / -0.90 / +2.17 |
| C13 | -1.73 / +1.88 / -27.74 | +21.33 / +9.67 / +5.67 | +0.33 / +0.54 / -0.91 |
| C14 | -37.53 / -17.07 / -24.69 | +24.00 / -11.00 / +5.33 | +0.43 / -0.58 / +0.73 |
| C15 | -49.16 / -78.83 / -48.84 | +1.33 / -4.00 / -22.33 | +0.31 / -0.56 / -0.03 |
| C16 | +23.74 / -7.86 / -18.38 | +4.33 / +16.67 / +3.33 | -0.03 / -0.98 / +0.28 |
| C17 | -35.50 / -35.54 / -63.24 | -1.33 / -18.67 / -28.33 | +0.25 / +0.40 / +1.97 |
| C18 | +51.95 / -8.56 / +36.58 | -29.00 / -0.33 / -4.33 | +0.58 / +0.80 / +0.00 |
| C19 | -9.55 / -0.20 / +25.85 | -64.00 / +17.33 / -5.67 | +1.43 / -0.44 / +0.14 |
| C20 | -27.21 / +20.04 / -41.61 | -4.33 / +9.67 / +24.33 | +0.43 / -0.22 / -0.18 |
| C21 | -24.32 / +39.19 / +7.30 | +42.33 / +2.33 / +26.33 | +0.83 / +0.53 / -0.30 |
| C22 | -16.05 / +26.10 / +12.15 | +11.67 / -8.00 / -14.67 | +0.38 / -0.08 / +0.29 |
| C23 | -29.16 / -0.74 / -7.94 | -36.67 / -48.33 / +30.33 | +0.38 / +0.04 / +0.98 |
| C24 | +22.47 / -35.13 / -31.89 | -33.00 / -17.67 / -3.67 | -0.45 / -0.16 / +0.54 |
| C25 | -10.80 / -0.54 / -13.97 | -28.67 / -5.00 / +4.33 | +0.04 / +0.37 / +1.01 |
| C26 | -20.61 / +51.96 / -1.46 | -0.67 / +46.33 / -1.00 | +0.61 / -0.72 / -0.56 |
| C27 | -35.62 / +13.01 / -48.33 | +36.67 / -11.67 / +22.67 | -0.08 / +0.46 / -0.76 |
| C28 | -19.67 / +3.29 / -53.30 | -19.67 / +32.00 / +9.67 | -0.69 / +0.10 / +1.12 |
| C29 | +29.43 / +28.39 / -22.93 | +0.33 / +3.33 / -29.00 | -0.61 / -0.13 / -0.27 |
| C30 | -14.68 / +38.84 / +33.12 | +7.00 / +3.00 / +7.00 | -1.43 / -0.08 / +0.12 |

Each cell reports runs 1 / 2 / 3. Values are candidate minus actual start: square angstroms for patch area, percentage points (pp) for hydrogen-bond occupancy, and degrees for P-axis circular SD. Each run uses a different initial-velocity seed and 3-ns production. Supplementary Data 4 provides absolute values, all descriptors, both reference comparisons and summaries across runs.

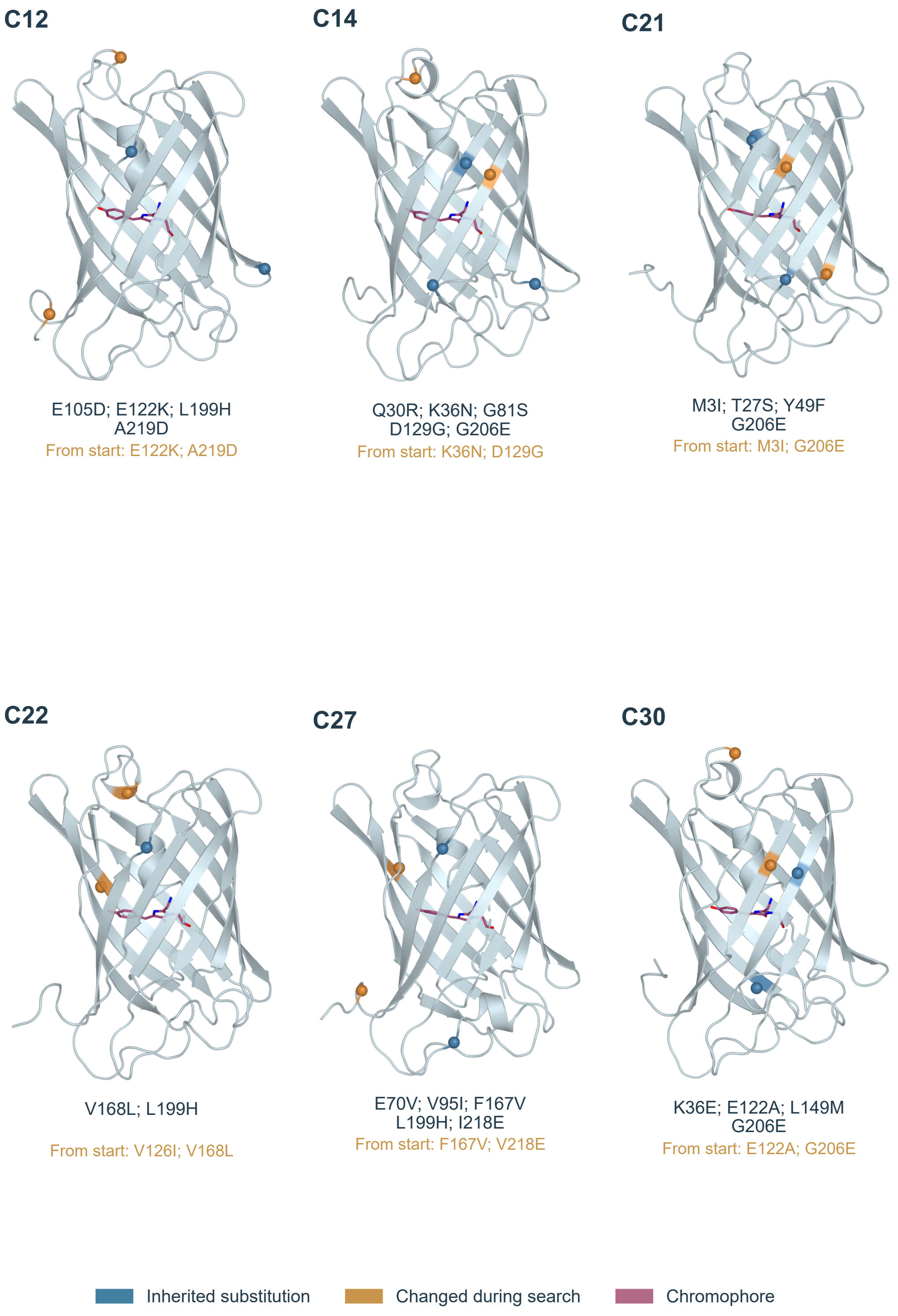


**Figure S13: Representative Top 30 sequences in their simulated mature folds**. Each image is an actual production conformation selected by proximity to the aligned time-average C-alpha structure. All use the same viewing direction. Blue marks inherited substitutions, amber marks positions changed from the actual start, and magenta marks the mature chromophore. Full WT-relative substitutions are listed below each structure, with the search-introduced changes shown separately. Positions 219–221 are modelled; C22's amber position 126 is a reversion to the WT residue. Each structure was obtained by candidate-specific relaxation from the experimental reference.

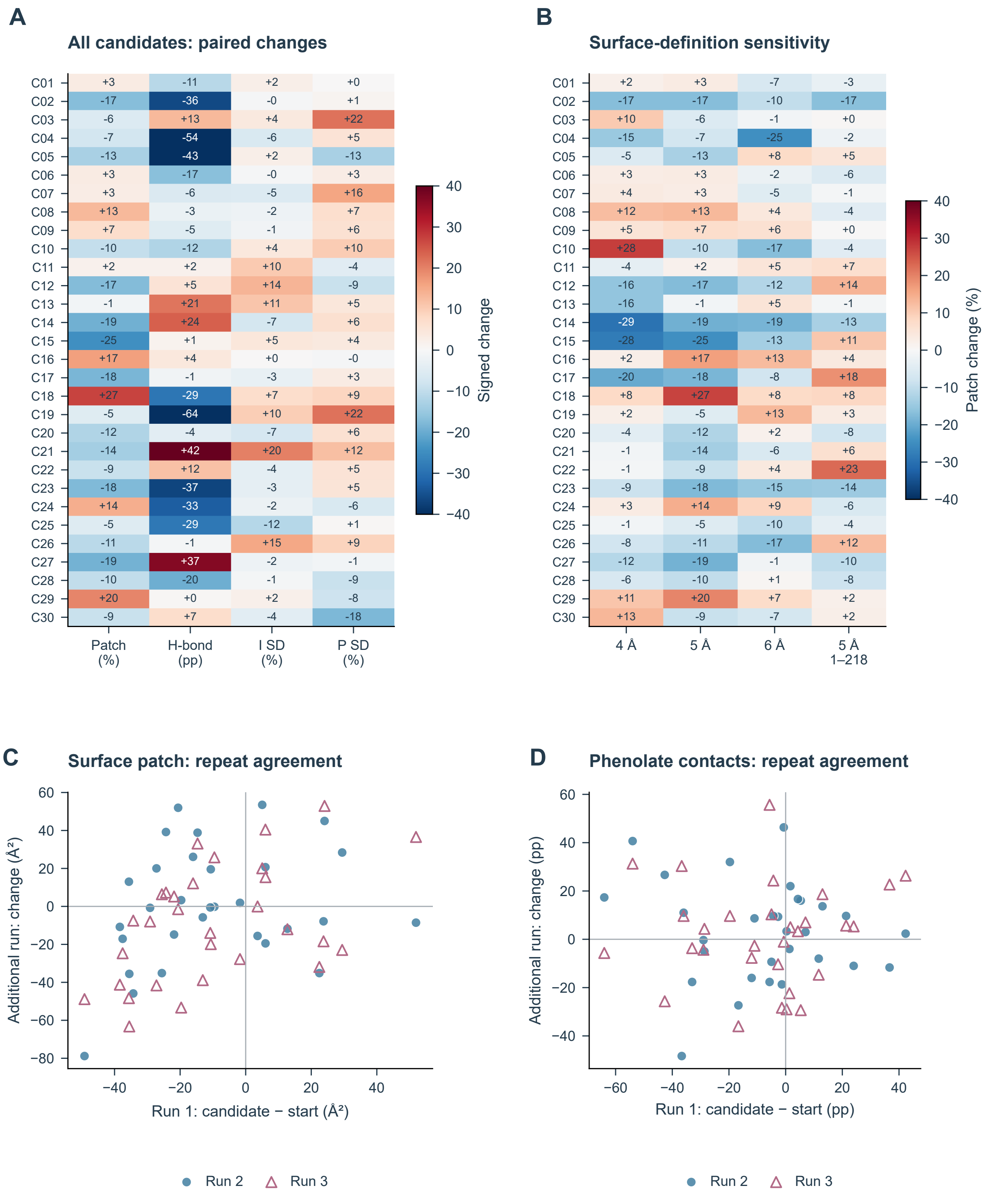


**Figure S14: Complete candidate changes and robustness checks**. **A.** First-run start-relative changes for all thirty candidates. Patch area and I/P circular SD are relative percentages; hydrogen-bond occupancy is a percentage-point difference. Colours show signed changes in each metric. The main text examines decreases in patch area and torsional spread and increases in phenolate occupancy. **B.** Patch changes for 4-, 5- and 6-angstrom radii and a 5-angstrom calculation restricted to resolved positions 1–218. The colour scale is clipped at ±40, while printed values are untruncated. **C,D.** Start-relative changes in runs 2 (circles) and 3 (triangles) against run 1 for all thirty candidates, showing surface patch area and phenolate occupancy separately. Each run has 3-ns production. Panel B uses run 1. Numerical results are in Supplementary Data 3–4.

#### S15 AI-assisted preparation

OpenAI Codex assisted with manuscript language editing, revision and figure preparation. Assistance with figure preparation covered plotting code, layout and graphical presentation.
